# Intrinsically disordered regions that drive phase separation form a robustly distinct protein class

**DOI:** 10.1101/2022.08.04.502866

**Authors:** Ayyam Y. Ibrahim, Nathan P. Khaodeuanepheng, Dhanush L. Amarasekara, John J. Correia, Karen A. Lewis, Nicholas C. Fitzkee, Loren E. Hough, Steven T. Whitten

## Abstract

Liquid-liquid phase separation (LLPS) of proteins is thought to be a primary driving force for the formation of membraneless organelles, which control a wide range of biological functions from stress response to ribosome biogenesis. LLPS of proteins in cells is primarily, though not exclusively, driven by intrinsically disordered (ID) domains. Accurate identification of ID regions (IDRs) that drive phase separation is important for testing the underlying mechanisms of phase separation, identifying biological processes that rely on phase separation, and designing sequences that modulate phase separation. To identify IDRs that drive phase separation, we first curated datasets of folded, ID, and phase-separating (PS) ID sequences. We then used these sequence sets to examine how broadly existing amino acids scales can be used to distinguish between the three classes of protein regions. We found that there are robust property differences between the classes and, consequently, that numerous combinations of amino acid property scales can be used to make robust predictions of LLPS. This result indicates that multiple, redundant mechanisms contribute to the formation of phase-separated droplets from IDRs. The top-performing scales were used to further optimize our previously developed predictor of PS IDRs, ParSe. We then modified ParSe to account for interactions between amino acids and obtained reasonable predictive power for mutations that have been designed to test the role of amino acid interactions in driving LLPS.

## Introduction

Many intracellular reactions occur within membrane-free compartments that form spontaneously from the cellular milieu (1). Examples of such compartments include P-bodies, Cajal bodies, the nucleolus, paraspeckles, and germ granules (2–4). The formation of membraneless organelles is facilitated primarily, though not exclusively (5, 6), by proteins that are intrinsically disordered (ID) or contain large ID regions (IDRs), collectively termed intrinsically disordered proteins (IDPs) (4, 7). Because these protein-rich droplets typically exist in dynamic, liquid-like states rather than as fixed complexes (1, 2), this transition is referred to as liquid-liquid phase separation (LLPS). By forming specific compartments and micro-environments, LLPS exerts control over the spatial organization and biochemical reactivity within cells (8, 9). Indeed, LLPS has been found to modulate chemical and biochemical reactions (10–12) and LLPS dysregulation has been associated with several human diseases (13–15).

Due to the critical role of protein LLPS in cell function and disease, significant efforts have been made to determine the physical mechanisms responsible for driving phase separation. Mutation and sequence analysis have demonstrated the importance of cation-π, π-π, π/sp^2^, and hydrophobic interactions (16–21). Groups of amino acids driving cohesive interactions are often characterized as “stickers” and are frequently interspaced with small polar residues acting as “spacers” (22–25). In addition, charge composition and patterning appear to contribute to the regulation of LLPS by IDRs (20, 26–29). Successfully predicting the relationship between primary sequence and phase separation behavior is key to understanding the molecular mechanisms underlying LLPS and to identifying the cellular processes that rely on LLPS. Effective predictive algorithms might also reveal how mutations affect LLPS-associated disease states.

Several methods have been developed to predict which protein sequences drive LLPS (30, 31). Algorithms including PLAAC, PSPredictor, and PSPer are based on the composition of databases of proteins that are known to drive LLPS (28, 32, 33). Others, including catGRANULE and CRAPome, are associated with cellular structures (34, 35). Uniquely, PScore was developed based on a specific mechanism thought to drive LLPS: the propensity of cation-π and π-π interactions to drive cohesive protein interactions (16, 36). Simulation models of IDRs have also been used to identify which protein domains drive LLPS as well as how mutations will affect LLPS behavior of those proteins (37–41). The diversity of successful approaches for predicting LLPS indicates that multiple complementary mechanisms may be responsible for this phenomenon.

We previously developed a predictive model of LLPS behavior, ParSe (“**Par**tition **Se**quence”), that identifies phase-separating (PS) IDRs starting from predictions of hydrodynamic size, which is indicative of the relative strength of intramolecular as compared to solvent interactions (42). The ParSe algorithm uses a sequence-calculated polymer scaling exponent, *v*_*model*_, to quantify hydrodynamic size (43, 44). When *v*_*model*_ is combined with a second sequence-based parameter, the intrinsic propensity for a sequence to form β-turns (45), the algorithm can distinguish between sequences belonging to one of three classes of protein regions: folded, ID, and PS ID (42). We proposed a physical mechanism whereby transient β-turn structures reduce the desolvation penalty of forming a protein-rich phase and increase exposure of atoms involved in π/sp^2^ valence electron interactions. In this mechanism, β-turns could act as energetically favored nucleation points, potentially explaining the observed higher propensity for turns in IDRs that drive phase separation *in vivo* (42, 45).

However, the prior study did not test whether the combination of *v*_*model*_ and β-turn propensity was uniquely able to distinguish folded, ID, and PS ID sequences, as would be required if this putative mechanism is necessary for LLPS. In the current study, we first curated the sequence training sets to expand the folded and ID categories. Our curated list of proteins that are ID but not thought to drive LLPS acts as a key negative control, enabling us to distinguish which features of IDRs in particular drive LLPS (31). Using the expanded sequence sets, we exhaustively tested all amino acid property scales found in the Amino Acid Index Database (46) for their ability to separate folded, ID, and PS ID sequences. We show that the three sequence sets are distinct in their means when quantified by the majority of amino acid scales, revealing that there are robust property differences between ID and PS ID sequences, not unlike the differences between folded and ID sequences.

We applied principal component analysis (PCA) to identify the extent of variability between our sequence sets and the optimal combinations of property scales that maximize the distinction between ID and PS ID sequences. The resulting predictor, ParSe version 2 (v2), uses sequence hydrophobicity to distinguish folded from ID, and, subsequently, *v*_*model*_ and a conformational parameter to distinguish ID from PS ID. In general, PS ID sequences exhibit enriched β-turn and depleted α-helix propensities. ParSe v2 more accurately predicts these regions from the amino acid sequence than the original version. We then compared our predicted propensity for LLPS with experimental results on mutant sequences designed to test the role of π- and charge-based interactions in LLPS behavior. We found that only by including effects representing interactions between amino acids could we accurately predict LLPS behavior of these mutants. Given the high fidelity of ParSe even in the absence of these interaction terms, it appears there are multiple diverse mechanisms that can drive LLPS and that PS ID sequences can be robustly identified through simple combinations of amino acid property scales.

## Results

### Construction of Protein Sequence Datasets

A limitation of the previous work, including our own (42), has been the relatively small set of sequences used to train predictors. We first sought to alleviate this problem by identifying additional sequences in our two negative control categories, folded proteins and IDRs, which are not thought to phase separate. The importance of well-defined negative control sets has been highlighted recently by Pansca et al (31) and Cai et al (47). For example, some negative control sets like the human proteome are known to contain many false negatives, which can lead to misassignments by the predictor.

We first expanded the set of folded proteins. Previously, we selected only folded regions found within known LLPS proteins. However, this selection may not be justified because it is not known whether folded regions within phase-separating proteins are biased differently in *v*_*model*_ and β-turn propensity compared to folded proteins in general. Subsequently, we expanded the previous folded set (comprised of 82 sequences) to include sequences from 122 human proteins with nonhomologous folded structures (48), 32 proteins with small (*N* = 36) to large (*N* = 415) folded structures (49), 54 folded extremophile proteins (50), 53 folded metamorphic proteins (51), and 90 folded membrane proteins (Table S1). Combined, these folded protein regions represent 421 unique sequences after removing duplicate entries. The folded sets were, overall, similar in both mean *v*_*model*_ (Figure 1A, Table 1) and mean β-turn propensity (Table 2) to the previous folded set obtained from known LLPS proteins (Tables S2, S3), indicating that folded regions within LLPS proteins are indeed similar to folded regions more generally.

**Table 1.**
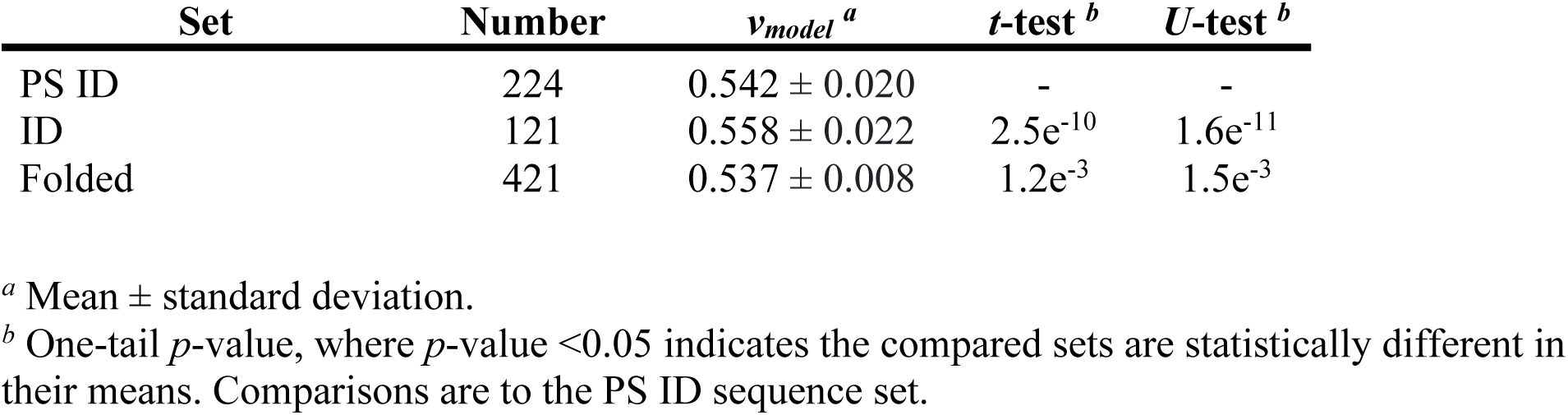
Summary of mean *v*_model_ in protein sequence sets.

**Table 2:**
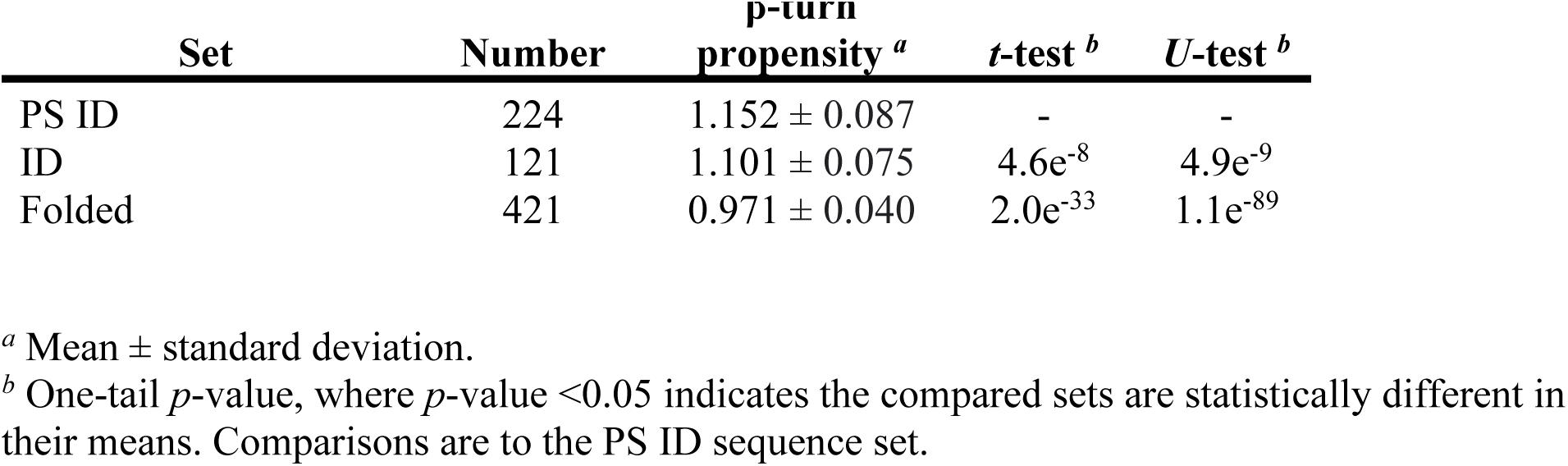
Summary of mean β-turn propensity in the protein sequence sets.

| Set | Number | β-turn<br>propensity <sup>a</sup> | *t*-test <sup>b</sup> | *U*-test <sup>b</sup> |
| --- | --- | --- | --- | --- |
| PS ID | 224 | 1.152 ± 0.087 | - | - |
| ID | 121 | 1.101 ± 0.075 | 4.6e<sup>-8</sup> | 4.9e<sup>-9</sup> |
| Folded | 421 | 0.971 ± 0.040 | 2.0e<sup>-33</sup> | 1.1e<sup>-89</sup> |

<sup>a</sup> Mean ± standard deviation.
<sup>b</sup> One-tail *p*-value, where *p*-value < 0.05 indicates the compared sets are statistically different in their means. Comparisons are to the PS ID sequence set.

**Figure 1.**
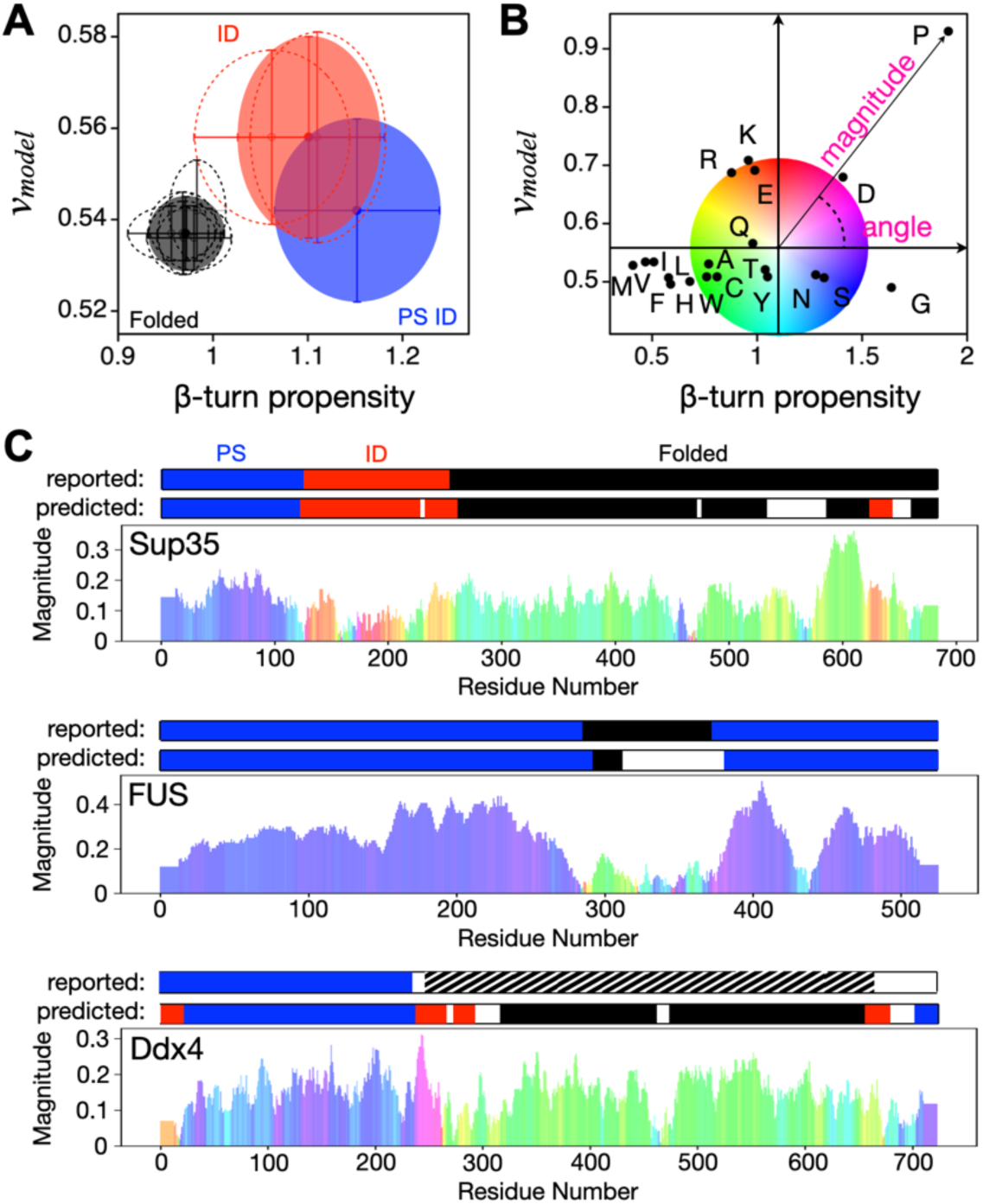
Sequence-calculated *v*_*model*_ and β-turn propensity separate protein regions by class. *A*, comparing *v*_*model*_ and β-turn propensity in each sequence set. Filled circles show the mean and standard deviation in *v*_*model*_ and β-turn propensity in the PS ID (blue), ID (red), and folded (black) sets. Open and dashed circles show the mean and standard deviation in individual subsets: previous ID and BMRB & DisProt (red); previous folded, human, small-to-large, extremophile, membrane, and metamorphic (black). *B*, comparing *v*_*model*_ and β-turn propensity in homopolymers (*N* = 100) where amino acid type is identified by its one-letter code. A centralized origin was mapped into this plot at the β-turn propensity and *v*_*model*_ values of 1.101 and 0.558, respectively, which are the means in the ID set. From this origin, every amino acid type can be represented by a distance magnitude and angular displacement; as shown for proline. A color wheel is used to convey angular displacement. *C*, magnitude/color plots are compared to the ParSe (original version) predictions for Sup35 (UniProt ID P05453), FUS (UniProt ID P35637), and Ddx4 (UniProt ID Q9NQI0), and to regions reported (i.e., identified) by experiment. Each figure shows the magnitude (y-axis) and color (angular displacement) by residue number (x-axis), as determined by amino acid type and its magnitude/color from panel B. ParSe predictions use blue (PS), red (ID), and black (folded). Striped represents ≥50% identity to a known folded protein.

Similarly, we expanded the set of IDR sequences not enriched for LLPS potential, called the “ID” set, by adding ID sequences found in the Biological Magnetic Resonance Bank (BMRB) (52) and DisProt (53, 54) databases. NMR experiments are typically performed at relatively high concentrations (≥100 μM), and so BMRB entries that do not explicitly address LLPS likely have a low propensity to phase separate. In addition, proteins known to drive LLPS are now annotated in DisProt; therefore, DisProt entries lacking such annotation are at least nominally depleted in LLPS drivers. Moreover, we only selected IDRs from DisProt that were both predicted to be disordered by MetaPredict (55) and were not highly homologous to proteins with folded structures in the Protein Data Bank (PDB) (56) using seqatoms (57). The combined ID set contains 121 unique protein domains (Table S4).

While these expanded datasets show slight differences in mean predicted *v*_*model*_ or β-turn propensity from the datasets used in our previous work (Tables 1, 2, S2, S3), the expanded sets reinforce our previous findings that there exist significant differences in *v*_*model*_ or β-turn propensity between the three classes of protein regions: folded, ID, and PS ID (Figure 1A). These results, as such, confirm that the two sequence-calculated metrics, *v*_*model*_ and β-turn propensity, can be used in combination, as done previously, to predict phase separating regions within proteins (42). Additionally, when *v*_*model*_ and β-turn propensity are calculated for homopolymers of the common amino acids and then presented in a β-turn propensity versus *v*_*model*_ plot (Figure 1B), the values obtained are consistent with previous characterizations of “order promoting” as compared to “disorder promoting” amino acids (Figure 1B). In particular, we find that homopolymers of Trp, Cys, Phe, Ile, Tyr, Val, Leu, Ala, His, Met, and Thr fall within the “folded” region of the β-turn versus *v*_*model*_ plot, and so are predicted to act as “order promoting” amino acids, while by similar analysis Arg, Gln, Pro, Glu, Lys, and Asp are “disorder promoting”, and Asn, Ser, and Gly are “phase separation promoting”. This result is similar to conclusions from analyses of protein structures (58, 59), where Trp, Cys, Phe, Ile, Tyr, Val, Leu, and Asn are enriched in folded proteins (“order promoting”), while Ala, Arg, Gln Pro, Glu, Lys, Gly, and Ser are enriched in IDPs (“disorder promoting”), and His, Met, Thr, and Asp are “ambiguous”.

The clear segregation of some amino acids into the PS ID sector of the β-turn propensity versus *v*_*model*_ plot motivated us to consider whether an approach as simple as color coding of the amino acids would enable identification of PS regions in proteins known to phase separate. Indeed, the PS driving regions of many proteins are visually apparent by our simple visualization tool based on the location of homopolymers in the β-turn propensity versus *v*_*model*_ plot (Figure 1B). The magnitude is related to the propensity and the color indicates the quadrant of the plot; therefore, a shaded bar chart predicts the propensity for a sequence to promote order, disorder, or phase separation. The rapid identification of PS regions in proteins (Figure 1C) such as Ddx4, FUS, and Sup35 (3, 17, 22, 60) led us to conclude that PS regions in proteins are distinctly different than other ID regions. We therefore sought to determine whether these classes of proteins were distinguishable by other amino acid property scales.

### Most amino acids property scales find significant differences between folded, ID, and phase-separating protein regions

We sought to determine if additional sequence-based intrinsic properties were significantly different between protein regions that are folded, ID, or ID with high potential for driving LLPS. To explore this idea, 566 scales of amino acid properties were obtained from the Amino Acid Index Database (46), which is a curated set of numerical indices representing various physicochemical and biochemical properties of the amino acids. This approach is similar to work done to improve coarse-grained models by testing multiple hydrophobicity scales (40). We added to these scales a newly developed hydrophobicity scale designed to predict sequences that drive protein LLPS (19), as well as *v*_*model*_. For each scale and for each sequence, we summed the amino acid scale for amino acids in the sequence, and divided by the length, *N*. Welch’s unequal variances *t*-test (61), given as a one-tail *p*-value, was used to find scales that show a statistical difference in the means of the sequence sets. Using the nonparametric Mann-Whitney *U*-test (62) gave overall similar results (Figure S1).

Figures 2A-C show that the different sequence sets have significantly different mean values for most scales when compared. For example, 81% of scales give *p*-values <0.05 (indicating means that are statistically different), when comparing ID and PS ID sequences (Figure 2A). Moreover, 13% and 22% of scales yield *p*-values smaller (thus showing a more significant difference) than the *p*-values obtained from *v*_*model*_ and β-turn propensity, respectively, used in ParSe (42). Each scale type (e.g., α-helix propensity, β-turn propensity, hydrophobicity, etc.) had some scales with very low *p*-values and some with *p*-values ≥0.05, suggesting that, overall, most, but not all, conformational and physicochemical based scales could substitute for *v*_*model*_ or β-turn propensity in ParSe and likely exhibit some ability for identifying PS IDRs from sequence. This analysis reveals that the physical differences between PS and conventional IDRs are robust across many different scales of amino acid properties (Figure 2A). We conclude that PS regions likely contain a variety of complementary, redundant sequence features that drive LLPS.

**Figure 2.**
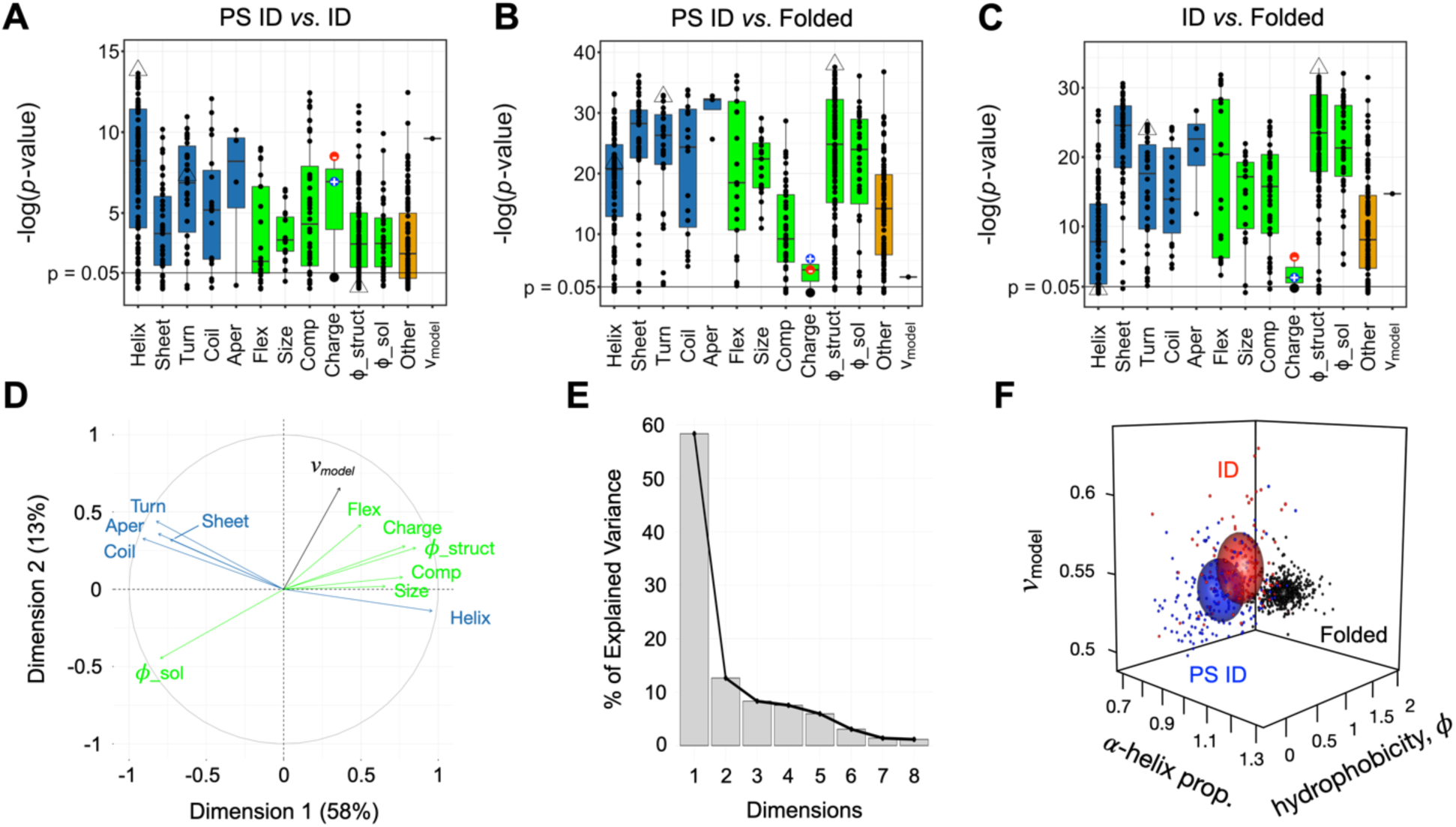
Robust differences in intrinsic sequence-calculated properties are found when comparing means by protein region class. *A-C, p*-values calculated by Welch’s unequal variances *t*-test, shown as -log(*p*-value), compares set means in 567 amino acid scales and *v*_*model*_. Conformation-based scales, highlighted by blue boxplots, are grouped by type according to α-helix (Helix), sheet or strand (Sheet), β-turn, tight turn, or reverse turn (Turn), coil or loop (Coil), and aperiodic (Aper) propensities. Physicochemical-based scales, highlighted by green box plots, are grouped by type according to flexibility (Flex), size (Size), composition (Comp), negative charge, positive charge, or net charge (Charge), and hydrophobicity (*ϕ*). Hydrophobicity scales were separated into two types: structure-based (*ϕ*_struct), where the scale is derived from a structural metric like burial or contact frequency in surveys of high-resolution protein structures, and solution-based (*ϕ*_sol), where the scale is obtained from solution studies like measuring the transfer free energy of the amino acids from water to an organic solvent. Scales (e.g., refractivity, crystal melting point) that did not easily map into a conformation- or physicochemical-based group were combined separately (Other). Boxplots show the dataset median (50^th^ percentile) with the central bar, and the vertical width spans the 25^th^ to 75^th^ percentiles. Open triangles highlight the smallest *p*-value when comparing means in the PS ID and ID sets (from an α-helix propensity scale), the smallest *p*-value when comparing means in either the PS ID or ID sets with the folded set (from a structure-based hydrophobicity scale), and the β-turn propensity scale used in ParSe. *D*, bidimensional plot from PCA showing the modes of variance in the combined ID set (PS ID and ID) arising from conformation- (blue arrows) and physicochemical-based (green arrows) scales relative to the two principal components of variance, given as Dimension 1 and Dimension 2. *E*, scree plot showing the percent of the total variance in the combined set of ID sequences that is captured by each principal component (i.e., dimension). *F*, sequence calculated *v*_*model*_, α-helix propensity, and hydrophobicity for the sequences in the PS ID (blue), ID (red), and folded (black) sets; spheres show the set mean ± σ.

The differences between folded and ID (both ID and PS ID) datasets are also robust to different scales of amino acid properties (Figures 2B-C). 95% and 93% of scales produced *p*-values <0.05 when means were compared between the folded and PS ID, and folded and ID sets, respectively. Almost all amino acid property scales yield statistically different means when comparing ID and folded sequences; the best performing scales were based on hydrophobicity. Those hydrophobicity scales with the lowest *p*-values when comparing means in the folded and ID sets had among the highest *p*-values when comparing means in the ID and PS ID sets (and *vice versa*), consistent with our previous findings that a single metric was insufficient to separate the three datasets.

### PCA identifies two principal modes of variation between proteins

We next sought to determine the degree to which amino acid scales could be combined without significant redundancy when comparing protein sequences. To do so, we used principal component analysis (PCA), which characterizes the variability in a dataset (63), in this case variability arising from different scales being applied to our sequences. Our primary focus is on distinguishing PS IDRs from conventional IDRs because many disordered predictors already exist to separate folded from disordered domains (55, 64, 65). We first selected the scale in each scale type (listed in Figure 2A) with the smallest *p*-value when comparing the ID and PS ID sets; that is representative scales from each type that are best able to separate ID and PS ID sequences. We additionally included *v*_*model*_, which we found previously to give complementary information to β-turn propensity. Each scale was then used to calculate sequence properties via a sliding 25-residue window applied to protein domains in a combined set including both the ID and PS ID datasets or the human proteome. We used a sliding window to avoid averaging properties between regions of proteins with different characteristics (42).

The results of the PCA indicate that most of the variability measured by high-performing scales within these datasets can be captured by 2-3 parameters (Figures 2D-E, S2). For both the combined ID dataset including ID and PS ID sequences and the human proteome, approximately 70% of the variability is captured by the first two principal components. Moreover, 58% of the variability in the combined ID set is captured by a single component. The variance arising from conformational propensity scales tend to cluster, as do those with physicochemical metrics like charge, hydrophobicity, and other compositional details. These results are robust to both the number of top-performing scales chosen and to the choice of reference set; we saw similar clustering when we extended this analysis to include the top three performing scales in each type and to the entire human proteome (Figure S2).

Within these two categories (conformational propensity and physicochemical metrics), high-performing scales function very similarly. As such, the predictive capabilities of amino acid scale combinations within each category are limited. In particular, turn and coil scales applied to protein sequences yield strongly correlated modes of variation that also are mostly anti-correlated with the variance produced from α-helix propensity scales (Figures 2D, S2). In our previous work, we proposed that β-turns could serve as a site for cohesive interactions between protein chains, driving LLPS (42). Our current results, while consistent with this hypothesis, show that this hypothesis cannot easily be distinguished from other structural hypotheses, e.g., that coils drive LLPS or helix inhibits LLPS, because the variation between these scales when applied to our datasets are all highly correlated. In contrast, the variances arising from hydrophobicity, charge, or *v*_*model*_ in our datasets have patterns that, in general, are different from the variances arising from turn, coil, and α-helix conformational propensities.

To illustrate the separation obtained when using complementary top-performing scales, we selected three scales to best separate our three datasets: 1) the top performing hydrophobicity scale for separating folded from either ID set (from Vendruscolo and coworkers (66)), 2) the top performing conformational scale in separating ID from PS ID sets, in this case one predicting α-helical propensity (from Tanaka and Scheraga (67)), and 3) *v*_*model*_ because it was most orthogonal to the latter helix scale in the PCA of our combined ID datasets. As can be seen in Figure 2F, significant separation is observed between our different datasets using these three intrinsic sequence properties. In general, the folded domains occupy a region with φ >0.08, and the greatest separation between the two disordered sets is observed in the α-helix/*v*_*model*_ plane.

When this approach is used to assess homopolymers of the common amino acids by their placement into a plot of hydrophobicity, α-helix propensity, and *v*_*model*_, the homopolymer results predict that Trp, Cys, Phe, Ile, Tyr, Val, Leu, His, and Met are “order promoting” amino acids, while Ala, Arg, Gln, Pro, Glu, Lys, and Asp are “disorder promoting”, and Asn, Ser, Thr, and Gly are “phase separation promoting” (Figure S3), similar to what we found previously (Figure 1B).

### Predicting folded, ID, and phase-separating protein regions from sequence

Next, we used the separation obtained from this method to identify protein sequences belonging to folded, ID or PS ID categories, analogous to what we did for ParSe. Our aim was to see if using these top-performing scales would provide better predictions of PS ID domains. We modified the algorithm making a second-generation version, ParSe version 2 (v2). In this version, as with the original (42), we apply a 25-residue window and then slide this window across a whole sequence in 1-residue steps (Figure 3A) to label individual amino acids as either P (for PS ID), D (for ID) or F (for folded), and then to regions that are at least 90% of any one of these labels (see Methods, Figure 3C). Both ParSe v1 and v2 accurately delineate regions of Sup35 that have been experimentally determined (60) to behave as ID, PS ID, or folded regions (Figure 3C), and good accuracy is similarly found for other well-studied proteins (3, 17, 22, 68–72) utilizing diverse reported mechanisms driving LLPS (Figure S4).

**Figure 3.**
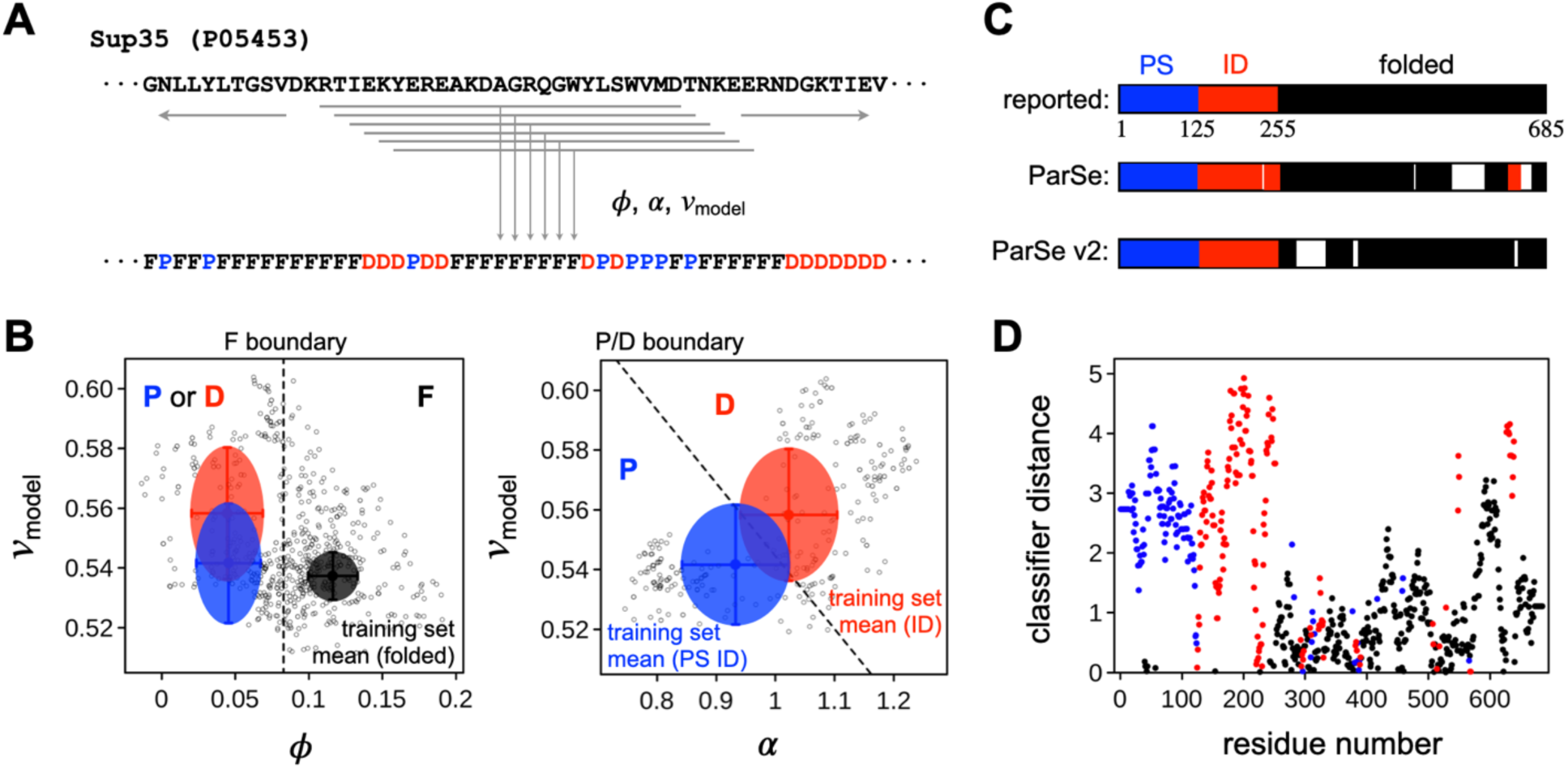
Predicting protein regions from sequence using the ParSe v2 algorithm. *A*, a sliding window algorithm is used to identify from sequence regions within a protein that match the PS ID, ID, and folded classes. Hydrophobicity (*ϕ*), α-helix propensity (α), and *v*_*model*_ are calculated for each contiguous stretch of 25-residues, or “window”, in the primary sequence. *B*, each window is assigned a label, F, P, or D, depending on the values of *ϕ*, α, and *v*_*model*_. In the left figure, open circles are *ϕ* and *v*_*model*_ calculated for each 25-residue window in the Sup35 sequence (UniProt ID P05453); filled circles are the mean ± σ in *ϕ* and *v*_*model*_ in the ID (red), PS ID (blue), and folded (black) sequence sets. Windows with *ϕ* ≥ the folded set mean - 2σ (dashed line) are labeled F. For windows with *ϕ* < the folded set mean - 2σ, the label is determined by α and *v*_*model*_; P for low α with low *v*_*model*_, or D for high α with high *v*_*model*_, as shown in the right figure. Filled circles show the mean ± σ in α and *v*_*model*_ in the ID (red) and PS ID (blue) sets. *C*, contiguous regions (*N* ≥20) in the Sup35 primary sequence that were 90% of only one label P, D, or F are colored blue, red, or black, respectively, to represent predicted PS, ID, or folded regions. Predictions from the original ParSe and ParSe v2 are compared to the reported regions identified by experiment. *D*, classifier distance of each window, assigned to the central residue of the window and then colored according to its label P (blue), D (red), or F (black).

One advantage of our algorithm is that it is very fast, and so can easily be applied to large datasets, e.g., the human proteome. We measured the prevalence of protein regions predicted by ParSe v2 to have LLPS potential in the human proteome (Figure 4) by two methods. First, as previously, we measured the longest predicted region with high LLPS potential (contiguous regions that are at least 90% labeled P). The results from ParSe v2 are mostly identical to results obtained previously using ParSe (42), whereby only ∼5% of proteins in the human proteome have a predicted P-labeled region that is at least 50 residues in length. Disordered regions taken from DisProt (minus the LLPS annotated IDPs) (53, 54) and folded regions taken from SCOPe (Structural Classification of Proteins extended, version 2.07) (73, 74) gave results mirroring the human proteome result in the sense that these sequences are mostly devoid of long regions predicted to have high LLPS potential. In contrast, the 43 proteins assembled by Vernon *et al* (16) that have been verified *in vitro* to exhibit homotypic phase separation behavior tend to contain long stretches labeled P by ParSe v2, with ∼90% of this set having predicted PS regions ≥50 residues in length. Only ∼63% of the 98 parent proteins from which the PS ID set was derived have predicted PS regions ≥50 residues, wherein not all in this set have been shown to phase separate as purified components.

**Figure 4.**
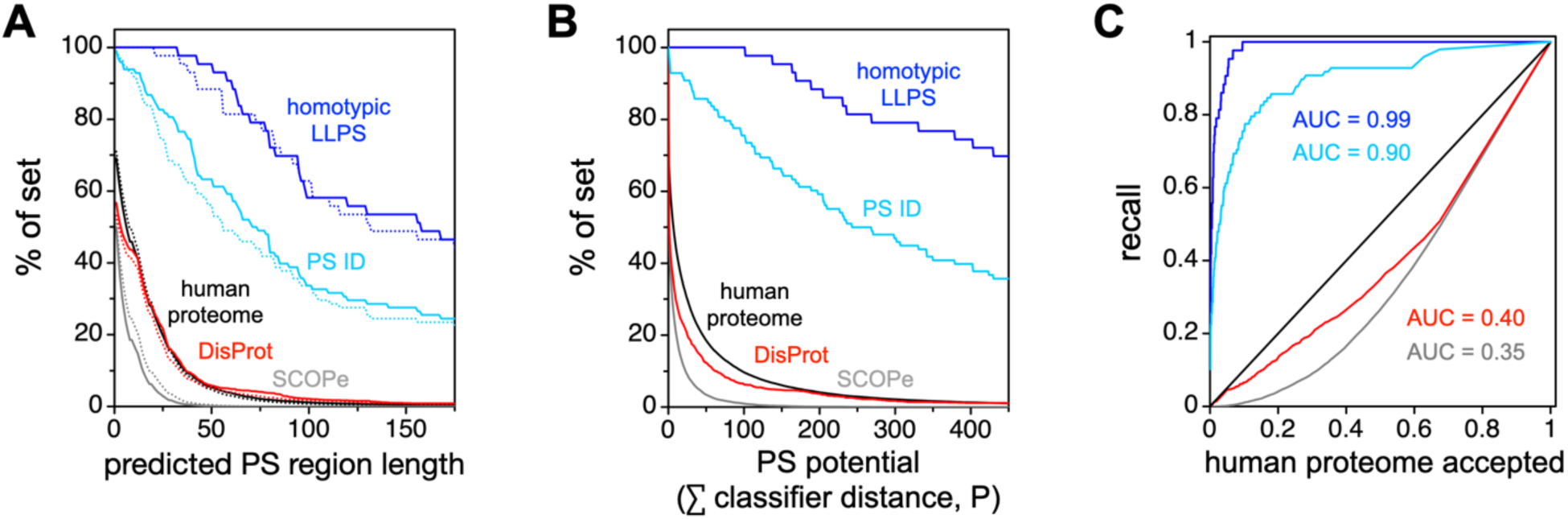
ParSe predicted PS regions are rarely found in the human proteome. *A*, ParSe (stippled lines) and ParSe v2 (solid lines) were used to identify regions in proteins that were ≥90% labeled P, which are referred to as phase-separating, PS, regions. Shown by the y-axis is the percent of proteins in a set with PS regions at least as long as the length indicated by the x-axis. The human proteome (UniProt reference proteome UP000005640) is given by black lines; DisProt (minus LLPS annotated entries) by red lines; SCOPe (version 2.07) by grey lines; a set of *in vitro* sufficient homotypic LLPS proteins by blue lines; and the full sequences of the proteins in the PS ID set by light blue lines. *B*, the summed P classifier distance was calculated by ParSe v2 for the protein sets in panel A. Shown by the y-axis is the percent of proteins in a set with a summed P classifier distance at least as much as the value indicated by the x-axis. Lines were colored using the same coloring scheme as in panel A. *C*, reproduction of the results in panel B wherein each set was directly compared to the human proteome result. Here, lines show the % of a set (using the same coloring scheme) plotted against the human proteome % of set for values of the summed P classifier distance.

Second, we developed a numerical score to give a quantitative measure of the confidence of our assignment of P, F and D labels, and to give a single metric to define the LLPS potential of every protein. Our justification for using a single numerical score is, in part, the dominance of a single principal component in the PCA of the combined ID set (Figure 2E), although we generalized this approach to F-labeled positions as well. In the combined ID datasets, most of the variability was in a single direction nearly orthogonal to the line separating P and D sectors in our plot. As such, we used the linear distance of a 25 amino-acid window into its classifier sector (i.e., F, D, or P sector), relative to the cutoff boundary and normalized by the distance to the boundary of the training set mean (Figure 3B). Values greater than 1 in this *classifier distance* indicate a window located at a distance further from the sector boundary than the distance of the training set mean, whereas values less than 1 indicate a window closer to the cutoff boundary than the training set mean and, as such, possibly with some uncertainty for its classifier label. Classifier distances calculated from the Sup35 sequence are shown in Figure 3D, wherein window values have been assigned to the central residue of the window, as we did with the window label.

We used the summed classifier distance for every window labeled P to obtain an overall score for each protein. This method is more robust to situations where multiple smaller regions drive phase separation (Figure S4), as compared to, e.g., Sup35, where a single domain drives phase separation. Windows labeled either F or D do not contribute to this sum. The assumption we are making is that single regions that promote phase separation are sufficient to drive phase separation of larger proteins. This is consistent with the observation that many PS proteins still undergo LLPS when they contain other protein regions or GFP tags (2, 75). As before, we found that only a small fraction of the human proteome consists of proteins with IDRs driving phase separation (42). Indeed, using a cutoff for the summed classifier distance of 100 retains 100% and 76% of the proteins in the Vernon et al *in vitro* sufficient set and the parent proteins of our PS ID set, respectively. In contrast, only 10% of human proteins are predicted to drive phase separation through their IDRs by this cutoff (Figure 4B). Because we are focused solely on IDRs which drive phase separation, excluding multivalent interactions that involve ordered domains, nucleic acids, or other drivers of LLPS, the total number of LLPS drivers is somewhat larger than this.

We used this whole protein metric (the summed classifier distance of P-labeled windows) to create a recall plot, used to assess prediction performance, for multiple datasets (Figures 4C, S5). The success in recall plots is typically quantified using the area under the curve (AUC), when comparing a test dataset to a comparison dataset (47, 76, 77). Here, in all cases, we used the human proteome as the comparison dataset. The SCOPe database and DisProt (excluding LLPS annotated entries) both have AUC values < 0.5 (Figure 4C), indicating that the human proteome does contain more proteins predicted to drive LLPS than these negative control groups. As a result, this approach likely gives a lower bound on the success of a predictor. As expected, our calculated AUC using ParSe v2 is highest on the *in vitro* sufficient LLPS drivers from Vernon *et al* (AUC = 0.99, Figure 4C), which constitute a significant fraction of our positive control dataset (i.e., the parent proteins of the PS ID set). This is likely both because this is the dataset we used for training and because it is also the most highly curated dataset. To further test its efficacy, we measured AUC values for ParSe v2 on datasets of LLPS drivers curated by other groups (16, 47, 76–78), and found it to perform quite well, with AUC values >0.8 (Figure S5).

Figures 4A and S6 suggest ParSe v2 is an improvement (i.e., slightly higher recall), albeit marginally, compared to the original ParSe. The strong performance of ParSe v1 is, in part, because even in the original version, we used scales that gave strong separation between datasets. Utilizing scales with weaker predictive value leads to a less efficient predictor, as expected (Figure S7). A comparison between ParSe v2 and ParSe v1 predictions reveals that the same patterning of P, D, and F regions appears for both predictors (Figure S8).

We then sought to compare ParSe to other published predictors. Although their data are not as highly curated as others, recent published work by Chen et al included predictions from multiple predictors on a publicly available dataset, facilitating comparison to other LLPS predictors (76). Of note, the negative control set in Chen et al contains, by our prediction, a higher fraction of IDRs driving phase separation than the human proteome (Figure S9D), although whether this is a problem with the database or with our prediction method is unclear. On their datasets, ParSe performs similarly as measured by AUC scores, to PScore (16), CatGranule (34), and PLAAC (32) in identifying proteins that drive LLPS (Figure S9A-C). The quality of the test one can make of these predictors depends significantly on the quality of the datasets, and so a true test of these predictors will require significantly more experimental data from both positive and negative controls (31, 47).

### Predicting the effects from mutation on phase separation behavior

Despite its simplicity, ParSe can predict the IDR(s) driving phase separation for a wide range of known phase-separating proteins, including FUS, Ddx4, LAF1, and A1. Several of these proteins have been the targets of mutagenesis studies implicating specific interactions between amino acids (i.e., cation-π or cation-anion) in the formation of phase-separated droplets. Cation-π interactions are thought to occur between different amino acids in the chain, and the balance of residues, e.g., Arg and Tyr, is thought to be important for LLPS (16, 22, 36). Similarly, net charge per residue, as opposed to simply the number of negative or positive charges (39), as well as the specific charge pattern (27), are also thought the be key determinants of LLPS.

Because ParSe is based only on the amino acid composition, and so does not include these higher-order effects involving combinations of amino acid types, we hypothesize that ParSe will have little predictive value for mutations that specifically alter the ratio of these pairwise interactions. More generally, we sought to determine if ParSe v2 could model the effects on phase separation behavior arising from mutations in the protein sequence. We hypothesize that sequence changes targeting P-labeled positions would have the greatest ability to modulate phase separation behavior. To assess this idea, we used the classifier distance whereby a phase separation “potential” was modeled as the summed classifier distance of P-labeled windows in the protein, as we did above in the recall plots. We compared the summed classifier distance with quantitative measures of LLPS behavior from four mutational studies involving three IDRs that individually exhibit LLPS behavior *in vitro* as purified components (3, 18, 27, 39) with sets of published mutations modulating either charge patterning or π-based interactions (Figures 5, S10).

**Figure 5.**
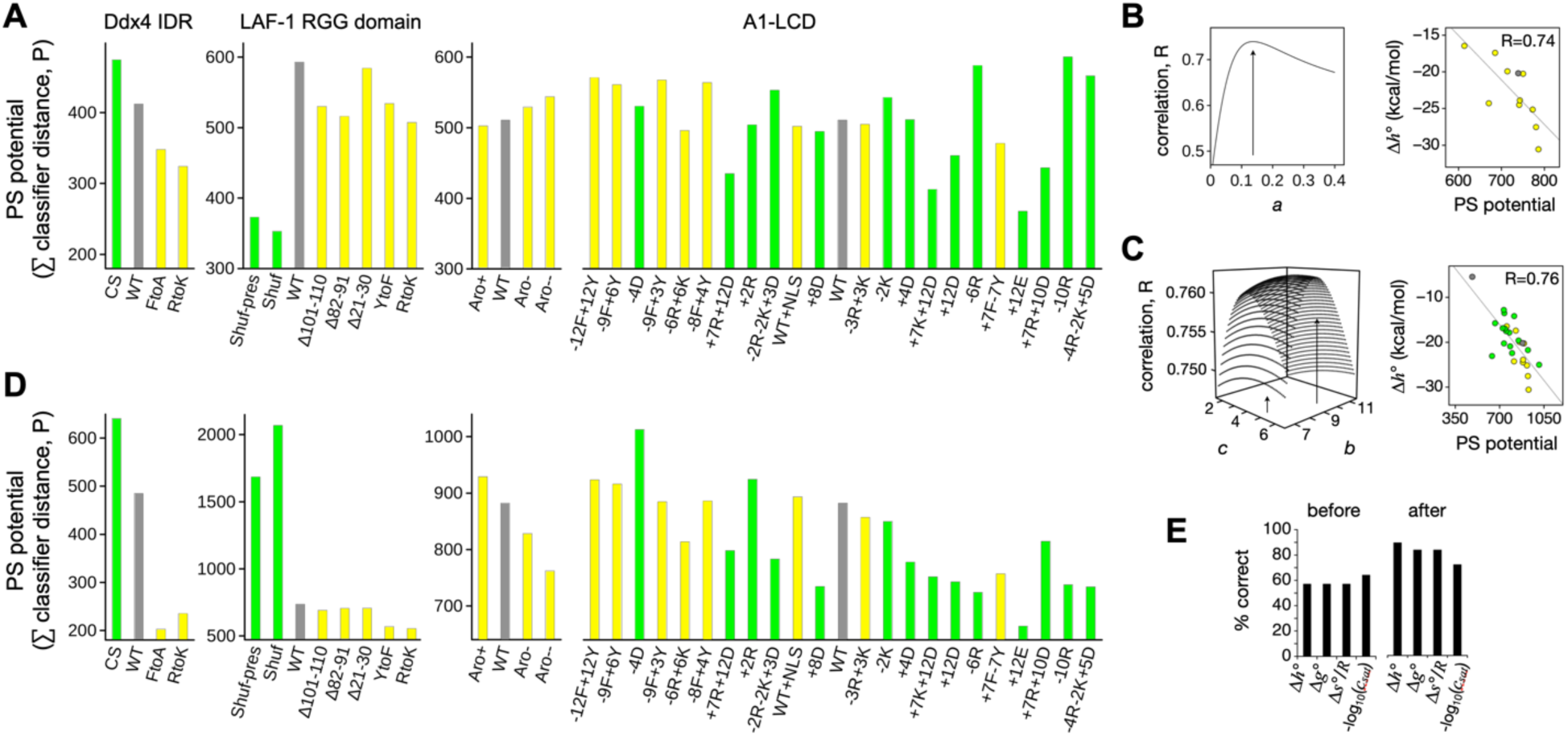
Predicting mutation effects on phase separation behavior. *A*, the summed classifier distance of P-labeled windows was used to calculate a phase-separating (PS) potential from sequence. Mutants were grouped by experimental study and colored grey for wildtype (WT), yellow for mutants with both *NCPR* and *SCD* identical to the WT values, and green otherwise (non-WT *NCPR* and *SCD*). Placement left-to-right within a study follows the reported PS potential in rank, from high-to-low, for comparison to the predicted PS potential. A1-LCD mutants used Δ*h°* and not *c*_*sat*_ to establish rank. *B*, A1-LCD mutants with *NCPR* and *SCD* matching the WT values were used to fix *a* in Equation 3 by optimizing the correlation of Parse-calculated PS potential (including *U*_*π*_) to Δ*h°*; the right figure shows the optimal correlation. *C*, similarly, all A1- LCD and Ddx4 mutants with experimental Δ*h°* were then used to fix *b* and *c* in Equation 4 by optimizing the correlation of ParSe-calculated PS potential (including *U*_*π*_ and *U*_*q*_) to Δ*h°*; the right figure shows the optimal correlation. *D*, ParSe-calculated PS potentials (including *U*_*π*_ and *U*_*q*_ optimized to Δ*h°*) for the mutant and WT sequences. *E*, percent of mutants correctly predicting an increase or decrease in PS potential relative to the WT before and after including *U*_*π*_ and *U*_*q*_ in the calculations. Results are binned according to experimental value that was used to fix *a, b*, and *c* in *U*_*π*_ and *U*_*q*_.

As the different studies used different metrics to quantitatively assess phase separation, we first began by simply asking whether the summed classifier distance could accurately reproduce the rank ordering of variants. In Figure 5A, we ordered, from left-to-right, in decreasing phase separation “potential” as reported within each individual study the mutant and wild type sequences. Shown is the summed classifier distance of P-labeled windows. In the LAF-1 RGG study (27), mutants forming phase separated droplets at elevated temperatures indicated increased phase separation potential, whereas changes in the saturation concentration, *c*_*sat*_, at a given temperature was used in studies with A1-LCD (18, 39). However, the mutant rank order in *c*_*sat*_ can change with the temperature; caused by differences in the standard molar enthalpy associated with phase separation, Δ*h°*, which reflects the temperature dependence to *c*_*sat*_. To manage this issue, mutant data were separated into two sets. One set corresponding to those mutants with experimental *c*_*sat*_ at 4 °C (Table S6), and a second corresponding to those mutants with experimental Δ*h°*, Δ*s°*, and Δ*g°* (Table S5). Figure 5A shows rank order in Δ*h°* for the A1-LCD mutants. Figure S10 ranks the A1-LCD mutants according to *c*_*sat*_ at 4 °C. The summed classifier distance (i.e., ParSe v2 predicted PS potential) of each mutant trended somewhat with the experimental rank order, correctly predicting an increase or decrease relative to the wild type in ∼60% of the mutants as presented in Figure 5 (i.e., with A1-LCD mutants ranked by Δ*h°*) and ∼65% in Figure S10 (i.e., with A1-LCD mutants ranked by *c*_*sat*_). Thus, ParSe is only moderately able to predict the effects of mutations designed to disrupt pairwise interactions between amino acids such as those arising from aromatic, cation-π, and charge-based interactions. This performance is similar to the performance of PScore, PLAAC, and catGranule (Figure S11).

To test the importance of pairwise interactions, we explicitly included different types of interactions in our model to try to account for these contributions and possibly improve the trend of calculated potential *versus* observed phase separation behavior. We expanded our calculation of LLPS potential to include both the summed P classifier distance and terms quantifying the effects of interactions between amino acids, termed *U*_*π*_ for π-π and cation-π interactions, and *U*_*q*_ for charge-based effects. The contribution of these terms toward predicting the effects of mutations can give information on the relative importance of the individual terms. We used *c*_*sat*_, Δ*h*°, Δ*s°*, and Δ*g°* separately to train this calculation; via 31 A1-LCD variants with *c*_*sat*_ and 27 A1-LCD and Ddx4 variants with Δ*h*, Δ*s°*, and Δ*g°* (Figures 5, S10). As *c*_*sat*_ is highly sensitive to the temperature (39), we expected the thermodynamic properties to be the more reliable metrics of LLPS. Indeed, we were best able to predict the effects of sequence changes on the measured Δ*h°* (Figure 5E). The predicted PS potential combining summed classifier distance with *U*_*π*_ and *U*_*q*_ correctly predicts the directional change relative to wild type in ∼90% of the mutants when *U*_*π*_ and *U*_*q*_ were trained against Δ*h*°, and the correlation between experimentally measured Δ*h°* and ParSe-calculated PS potential was reasonably high (R=0.76; Figure 5C). Thus, explicit consideration of interactions between amino acid types is important for determining PS potential in these mutational studies. It remains to be seen whether ParSe is able to accurately predict PS potential of mutants designed to test other aspects of LLPS, such as its dependence on the presence of partner molecules or on a specific set of solution conditions (e.g., pH, ionic strength, temperature).

Finally, we sought to determine what effect including these corrections to ParSe had on the identification of proteins driving phase separation. Overall, including *U*_*π*_ and *U*_*q*_ into ParSe increases the number of proteins identified that drive phase separation in both the PS sets and the human proteome (Figure S12). As a result, the AUC when comparing either our PS ID set or the Vernon highly curated set to the human proteome is slightly reduced. However, whether this is a result of correctly classifying more human proteins as driving LLPS, or whether we have simply increased the false negative rate remains to be seen.

## Discussion

In this work, we focused on identifying IDRs that drive phase separation, with a particular focus on separating PS IDRs from conventional IDRs that do not drive phase separation. Using carefully curated datasets of ID, PS ID and folded domains (Figures 1, 2), we developed a sequence-based predictor of phase separation (ParSe; Figure 3) which is fast enough to scan the entire human proteome in minutes on a single computer, and as or more accurate than other published predictors in identifying both proteins and regions within proteins that drive phase separation (Figures 3, S4, S5, S9, S11). We recognized that a wide variety of amino acid scales show significant differences between the ID and PS ID datasets, indicating that PS IDRs are a robustly different class of protein region than non-phase separating IDRs (Figure 2). We conclude that a redundant combination of molecular mechanisms driving cohesive interactions between amino acids is likely at play. This helps to explain why our general predictor of IDR hydrodynamic size (*v*_*model*_) is a strong indicator of LLPS potential, as we found previously (42). Moreover, by including interactions between amino acids thought to drive phase separation, we were able to match existing data on mutant sequences (Figure 5). This extension highlights the importance of pair-wise interactions in modulating phase separation.

While our approach has proved very successful, it, like other approaches to this problem, has significant limitations, including limitations in predicting responses to changes in solvent, limitations of the datasets, and limitations of the constraints of the approach chosen. The formation of phase separated droplets by polymer chains is a result, very generally, of interactions between chains that are stronger than the interactions of the chain with the solvent. As a result, LLPS is strongly dependent on the solution environment. Within cells, there are many proteins which assemble into membraneless organelles only within specific cellular conditions, e.g., upon lowering of pH (60). To accurately predict which solution conditions drive phase separation of any individual protein domain would require a detailed understanding of which mechanisms proteins use to drive phase separation, how those mechanisms are modulated by solutions conditions, and how cells modulate solution conditions in different cellular states. As a first step in this process, our aim is to simply improve identification of which IDRs and which potential mechanisms are used by IDRs to drive phase separation in a variety of cellular and solution conditions. Thus, although our predictor has high success in identifying proteins that have been seen experimentally to drive phase separation, we do not yet distinguish between responses to different cellular conditions, or, e.g., upper-versus lower-critical temperature. The temperature dependence of hydrophobicity scales as used by Dignon et al (79) could be a potential future approach to do this.

A primary limitation of our work, as well as others, is that even our well curated datasets have misidentified regions. For example, because the IDR in a protein that is responsible for phase separation has not always been identified, we simply used all IDRs from known phase-separating proteins. As a result, our PS ID set likely includes some IDRs which are not involved in phase separation. Similarly, our ID set was curated from proteins that have not yet been identified to phase separate, including those with experiments done at high protein concentration. However, the lack of observation of phase separation at any one experimental condition does not preclude its formation. Indeed, a long history of solution screening for crystallography would indicate that protein behavior can vary dramatically based on solution conditions (80). However, it appears that our PS ID and ID datasets are sufficiently enriched or depleted for PS IDRs for us to identify key properties of IDRs that drive phase separation. For example, the performance of our predictor is improved as the rigor with which the dataset was curated improves. ParSe gives the highest AUC on the dataset from Vernon et al containing only those proteins shown to drive homotypic LLPS *in vitro*, compared to datasets containing PS drivers more generally, and weaker still on datasets including both LLPS drivers and proteins recruited to existing droplets (Figure S5) (16, 76, 77).

Our approach is based primarily on sequence composition and not on sequence patterning or combinations of amino acids. It is surprising how effective this strategy is and how many different scales can be used to distinguish PS IDRs successfully. Nevertheless, our approach, while fast and effective, is unable to identify pairwise protein interactions that contribute to LLPS. In our analysis of mutants, we introduced a simple potential whereby amino acid pairs are counted, and this clearly improves the ability to predict the effects of mutation on phase separation (Figure 5). Pairwise interaction patterns are probably better identified by machine learning algorithms or simulation (27, 28, 40, 41, 47, 77). However, the efficacy of our approach appears to indicate that the primary determinant of whether any one sequence will phase separate depends on the overall amino composition, whereas rearrangements, mutations, or post-translational modifications of that base sequence will modulate that propensity for phase separation. Thus, it appears that the identification of sequences that have the potential to phase separate is an easier problem than identifying how mutation of a few residues will impact that phase separation potential. This result is not specific to our predictor, as none of the predictors tested here showed significantly better correlation with changes in phase separation potential upon mutation (Figure S11). We additionally note that different experimental measurements of LLPS potential give different ordering of mutants (Figures 5, S10), further compounding the issue.

Finally, our approach differs from several others in that we are focused solely on the problem of separating PS IDRs from IDRs that do not phase separate (47, 76). We are thus not able to identify proteins that utilize multivalent interactions between folded domains and other folded, ID, or nucleic acid binding domains as a primary mechanism for driving phase separation (5, 6, 24, 25). Moreover, we are primarily focused on IDRs that drive phase separation, as opposed to those that are recruited to existing phase separated droplets, a case which has been recently considered by Chen et al (76). Our motivation for this narrow focus is that a broader focus might obscure mechanisms used only by PS IDRs, and that interactions between folded domains are, in general, better understood than those between disordered domains.

The strong performance of ParSe on existing datasets, the robust nature of differences between PS IDRs and conventional IDRs, and the high correlation between ParSe and other predictors on databases of phase separating proteins all give confidence that ParSe is able to identify PS IDRs with significant accuracy. Because of its speed, ParSe can easily be applied to datasets of arbitrarily large size. As an example, we measured the summed classifier distance for the human proteome and found that only a small fraction of the human proteome is likely to drive phase separation (Figure 4B). Moreover, we identified the 500 proteins with the highest summed classifier distance in the human proteome, as well as their longest predicted PS ID region (Table S7). Many proteins involved in stress granule formation, RNA processing, and other functions known to be associated with membraneless organelles are identified in this process. However, many proteins are also identified that are not yet associated with a biological process driven by phase separation. This suggests that, while the fraction of human proteins driving phase separation may be small, not all of the biological processes relying on phase separation have yet been identified.

## Experimental procedures

### Protein databases

A set of 224 IDRs from proteins that exhibit LLPS behavior, used for the PS ID set, was obtained from our prior work (42). For the ID set, we started with 23 IDR sequences used previously (42), and then added all DisProt consensus ID sequences not having the disorder function ontology identifier for LLPS, IDPO:00041 (54). Protein sequences in the BMRB (52) with “disordered” or “IDP” as a keyword or in the entry title were also added to the ID set. BMRB obtained sequences were restricted to those with ≥70% of residue positions classified as disordered by Wishart’s random coil index, using an S^2^ cutoff of 0.6 (81). DisProt and BMRB sequences were culled by Metapredict (55), keeping only those predicted to be ID, and seqatoms (57), excluding those that were highly homologous to folded regions of proteins in the PDB. The folded set started with the 82 folded sequences used previously (42), and then added a set of human proteins with nonhomologous structures (48), proteins with small to large structures (49), extremophile proteins (50), metamorphic proteins (51), and membrane proteins that were found by searching the PDB (56) for the phrase “membrane protein.” Using the PISCES Server (82), the human, extremophile, metamorphic, and membrane proteins had a maximum of 50% sequence identity within each folded subset and only X-ray structures with a resolution better than 2.5 Å.

### Calculation of β-turn propensity and *v*_*model*_

The propensity to form β-turn structures was calculated by ∑ *scale*_*i*_/*N*, where *scale*_*i*_ is the value for amino acid type *i* in the normalized frequencies for β-turn from Levitt (83). The summation is over the protein sequence containing *N* number of amino acids. *v*_*model*_ was introduced previously (42) as a phenomenological substitute to the polymer scaling exponent (84, 85) and used to normalize protein hydrodynamic size to the chain length,

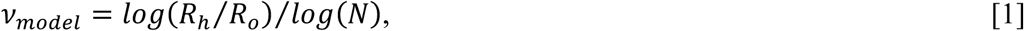

where *R*_*o*_ is a constant set to 2.16 Å, and the hydrodynamic radius, *R*_*h*_, is calculated from sequence using an equation found to be accurate for monomeric IDPs (43, 44, 86–88). The equation to calculate *R*_*h*_ for a disordered sequence is,

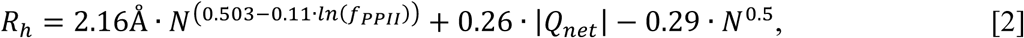

where *f*_*PPII*_ is the fractional number of residues in the PPII conformation, and *Q*_*net*_ is the net charge. *f*_*PPII*_ is estimated from ∑ *P*_*PPII,i*_/*N*, where *P*_*PPII,i*_ is the experimental PPII propensity determined for amino acid type *i* in unfolded peptides (89) and the summation is over the protein sequence. *Q*_*net*_ is determined from the number of lysine and arginine residues minus the number of glutamic acid and aspartic acid.

### Principal component analysis

The statistical program R (90) was used to perform PCA on the sequence sets, and the packages ggfortify, ggplot2, factoextra, MetBrewer, and tidyverse were used to render the results. In the PCA, the variables were shifted to be zero centered and scaled to unit variance.

### ParSe v2 algorithm

For an input primary sequence, whereby the amino acids are restricted to the 20 common types, ParSe v2 first reads the sequence to determine its length, *N*. Next, the algorithm uses a sliding window scheme (Figure 3A) to calculate *v*_*model*_, α-helix propensity, and *ϕ* for every 25-residue segment of the primary sequence. This window scheme can be applied to proteins with *N* >25. *R*_*h*_ is calculated by Equation 2, which in turn is used to determine *v*_*model*_ by Equation 1, by the same method used in the original ParSe described previously (42). α-helix propensity is calculated as the sequence sum divided by *N* using the scale by Tanaka and Scheraga (67). *ϕ* is calculated as the sequence sum divided by *N* using the hydrophobicity scale by Vendrusculo and coworkers (66). A window is labeled F if *ϕ* >0.08 (Figure 3B). If *ϕ* <0.08, a window is labeled P or D depending on the values of *v*_*model*_ and α-helix propensity. Windows with high α-helix propensity and high *v*_*model*_ are labeled D, while those with low α-helix propensity and low *v*_*model*_ are labeled P. The P/D boundary was determined by the line that bisects the overlapping distributions of *v*_*model*_ and α-helix propensity in the PS ID and ID sets, given by *v*_*model*_ = -0.244•α-helix propensity + 0.789. The window label is assigned to the central residue in that window. N- and C-terminal residues not belonging to a central window position are assigned the label of the central residue in the first and last window, respectively, of the whole sequence. Protein regions predicted by ParSe v2 to be PS, ID, or folded are determined by finding contiguous residue positions of length ≥20 that are ≥90% of only one label P, D, or F, respectively. When overlap occurs between adjacent predicted regions, owing to the up to 10% label mixing allowed, this overlap is split evenly between the two adjacent regions.

### Classifier distance calculation

The classifier distance is the normalized distance of a ParSe v2 generated window into its classifier sector (i.e., F, D, or P sector) and relative to the cutoff boundary (Figure 3B). For F labeled windows, the classifier distance is *ϕ* (of the window) minus the cutoff value of 0.08, and then normalized to distance of the folded set mean *ϕ* (0.1164) to the cutoff. Specifically, this is (*ϕ* – 0.08)/(0.1164 – 0.08). For P or D labeled windows, first we find the point on the P/D boundary (*v*_*model*_ = -0.244.α-helix propensity + 0.789) that makes a perpendicular bisector when paired with the window values of *v*_*model*_ and α-helix propensity. Then the distance between this point and the point defined by the window values of *v*_*model*_ and α-helix propensity is determined. Specifically, this distance is sqrt((α – x).(α – x) + (*v*_*model*_ – y).(*v*_*model*_ – y)) where α is the α-helix propensity, x is (α/0.244 + 0.789 – *v*_*model*_)/(0.244 + 1/0.244) and y is (x – α)/0.244 + *v*_*model*_. This distance is normalized by dividing by 0.019 (the distance from the boundary to either of the set means).

### PSCORE calculation

PSCORE, which is a phase separation propensity predictor (16), was calculated by computer algorithm using the Python script and associated database files available at https://doi.org/10.7554/eLife.31486.022.

### Granule propensity calculation

Granule propensity was calculated by using the catGranule (34) webtool available at http://www.tartaglialab.com.

### PLAAC LLR calculation

LLR score, which identifies prion-containing sequences (91), was calculated by using the webtool available at http://plaac.wi.mit.edu.

### Metapredict calculation

Metapredict score (55), which predicts the presence of ID in a sequence, was calculated by computer algorithm using the Python script available at http://metapredict.net.

### Calculation of *U*_*π*_

The relative contributions of aromatic and cation-π interactions to LLPS in our calculations followed the observed rank order by Wang *et al*: Tyr-Arg > Tyr-Lys ∼ Phe-Arg > Phe-Lys (22). To mimic this ranking, we assumed 3:2:1 weighting and, also, that Phe-Tyr interactions would contribute comparably to Phe-Lys interactions,

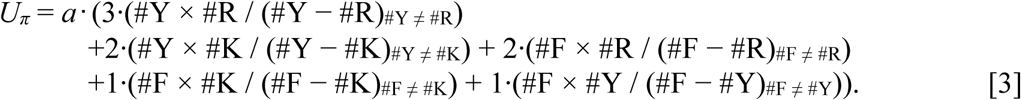

In Equation 3, #Y, #R, #F, and #K represent the number of Tyr, Arg, Phe, and Lys residues, respectively, in a sequence, calculated on a per-window basis, and *a* is a fitting parameter (see below). Thus, *U*_*π*_ increases with increasing Tyr, Arg, Phe, and Lys content, and more so when interaction partners are present at similar levels. When the divisor is zero (e.g., when #Y = #R), it is changed to 1 to avoid infinite potentials.

Window-specific *U*_*π*_ was added to the classifier distance at windows labeled P. Moreover, *U*_*π*_ was applied to D-labeled windows, allowing for the possibility of labels changing from D to P. This would occur when the value for *U*_*π*_ was larger than the classifier distance at a D-labeled window. Thus, protein regions that otherwise have characteristics more like the ID set, in *v*_*model*_ and α-helix propensity, could be labeled P if *U*_*π*_ was large enough. When this occurs, the given classifier distance was determined by the difference between *U*_*π*_ and the original classifier distance of the window formerly labeled D.

The parameter *a* in Equation 3 was determined by finding the optimal correlation of ParSe-calculated PS potential to Δ*h*° (finding *a* = 0.14; Figure 5B), Δ*s*° (finding *a* = 0.08), Δ*g*° (finding *a* = 0.11), or *c*_*sat*_ (finding *a* = 0.28; Figure S10B). In each case, the mutants used to fit *a* were limited to the subset with identical charge and charge patterns, determined by calculating the net charge per residue, *NCPR*, and sequence charge decoration, *SCD*, of each sequence. *NCPR* is the number of Lys and Arg residues minus the number of Glu and Asp residues, divided by *N. SCD* is calculated by *N*^-1^∑_*i*_∑_*j,j>i*_(*q*_*i*_*q*_*j*_)|*j-i*|^1/2^, where *q* is the amino acid specific charge (92).

### Calculation of *U*_*q*_

To model the contributions of charge-based interactions to LLPS, we build upon the observations by Schuster et al (27) and Bremer et al (39) that changes in *SCD* and *NCPR*, respectively, can affect phase separation potential. Accordingly, a simple charge-based potential was defined,

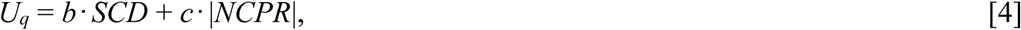

where *b* and *c* are fitting parameters, and *U*_*q*_ is calculated on a per-window basis. *U*_*q*_ is added to the classifier distance at each window labeled P, and is applied to windows labeled D, following the scheme described above for *U*_*π*_, again allowing for the possibility of labels changing from D to P. As with *a*, the parameters *b* and *c* were fixed by finding the optimal correlation of calculated PS potential and Δ*h*° (finding 8.4 and 5.6, respectively; Figure 5C), Δ*s*° (finding 4.6 and 7.0, respectively), Δ*g*° (finding 5.2 and 5.4, respectively), or *c*_*sat*_ (finding -16.0 and 33, respectively; Figure S10C).

### Calculation of Δ*h°*, Δ*s°*, and Δ*g°* from temperature dependence to *c*_*sat*_

For some Ddx4 and A1-LCD sequences, Δ*h°* and Δ*s°* (and thus Δ*g°*) were not available, but *c*_*sat*_ measured at different temperatures has been reported (3, 18). For these proteins, the standard molar chemical potential, *μ°*, was used to relate *c*_*sat*_ in the dilute and dense phases, *c*_*dilute*_ and *c*_*dense*_, respectively, to the standard molar enthalpy and entropy associated with phase separation (39),

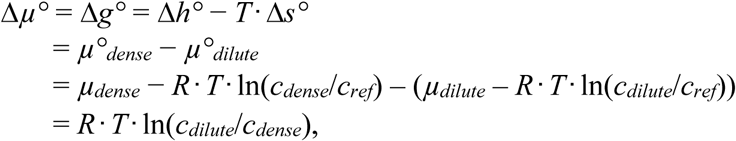

where *μ*_*dense*_ − *μ*_*dilute*_ is zero at equilibrium, *R* is the universal gas constant, and *T* is temperature. By plotting the natural logarithm of *c*_*sat*_ at different temperatures, a linear fit versus 1/*T* yields Δ*h°* and Δ*s°*. For A1-LCD mutants, 0.03 *M* was used for *c*_*dense*_ (39). For Ddx4 mutants, 0.01 *M* was used for *c*_*dense*_ (3). Δ*g°* was calculated from Δ*h°* − *T*.Δ*s°* and the standard temperature (273.15 K).

## Data availability

The Parse v2 algorithm written in Fortran, Parse_v2.f, can be downloaded at https://github.com/stevewhitten/ParSe_v2. A webtool version can be used at https://stevewhitten.github.io/Parse_v2_web.

## Author contributions

L. E. H. and S. T. W. conceptualization; A. Y. I., N. P. K., D. L. A., J. J. C., K. A. L., N. C. F., L. E. H., and S. T. W., formal analysis; K. A. L., N. C. F., L. E. H., and S. T. W., investigation; L. E. H and S. T. W., methodology; K.A.L., N. C. F., L. E. H, and S. T. W., writing–original draft; J. J. C., writing–review and editing.

## Funding and additional information

This work was supported by the National Institutes of Health under grants R25GM102783 (South Texas Doctoral Bridge Program; B. O. Oyajobi and S. T. W.), R35GM119755 (L. E. H.), and R01AI139479 (N. C. F.), as well as the National Science Foundation under grants 1818090 (N. C. F.) and 1943488 (L. E. H.). No nongovernmental sources were used to fund this project. The content is solely the responsibility of the authors and does not necessarily represent the official views of the NSF or NIH.

