## Supporting Information for "Intrinsically disordered regions that drive phase separation form a robustly distinct protein class"

Short title: Intrinsic properties of phase-separating IDRs

*Ayyam Y. Ibrahim, Nathan P. Khaodeuanepheng, Dhanush L. Amarasekara, John J. Correia, Karen A. Lewis, Nicholas C. Fitzkee, Loren E. Hough, and Steven T. Whitten*

### Contents:

Supporting Tables
S1. List of folded protein regions.
S2. Summary of mean $v_{model}$ in the ID and folded sequence subsets.
S3. Summary of mean β-turn propensity in the ID and folded sequence subsets.
S4. List of IDRs not known to exhibit phase separation behavior.
S5. Enthalpy, entropy, and free energy of phase separation of A1-LCD and Ddx4 mutants.
S6. Saturation concentration (at 4 °C) of A1-LCD mutants.
S7. List of 500 proteins with the highest summed P classifier distance in the human proteome.

Supporting Figures
S1. Comparing means in the sequence sets using a nonparametric test.
S2. Modes of variance in the sequence sets arising from different amino acid property scales.
S3. Comparing hydrophobicity, α-helix propensity, and $v_{model}$ in homopolymers.
S4. Predicting protein regions driving LLPS.
S5. LLPS driver sequences have AUC >0.8 when compared against the human proteome.
S6. ParSe v2 shows improved recall compared to the original version.
S7. ParSe v2 shows reduced recall when using scales with weaker predictive value.
S8. ParSe v2 sequence predictions exhibit the same LLPS patterns as ParSe v1 predictions.
S9. ParSe v2 shows similar predictive accuracy as other LLPS predictors.
S10. Predicting mutation effects on phase separation behavior by training against $c_{sat}$.
S11. ParSe v2 and other LLPS predictors show similar accuracy for predicting mutation effects.
S12. $U_{\pi}$ and $U_q$ effects on ParSe predicted PS regions.

Supporting References

### Supporting Tables

### Table S1. List of folded protein regions.

| Name | Database [a] | UniProt accession number | folded regions (*N*) [b] | PDB entries |
|---|---|---|---|---|
| PPP5C | Wang *et al* | P53041 | 19-177 (159) | 1a17.pdb |
| Galectin-3 | Wang *et al* | P17931** | 114-250 (137) | 1a3k.pdb |
| RB1 | Wang *et al* | P06400 | 378-562 (185) | 1ad6.pdb |
| CD40LG | Wang *et al* | P29965 | 116-261 (146) | 1aly.pdb |
| FABP5 | Wang *et al* | Q01469 | 3-135 (133) | 1b56.pdb |
| LALBA | Wang *et al* | P00709 | 20-142 (123) | 1b9o.pdb |
| CDKN2D | Wang *et al* | P55273 | 7-162 (156) | 1bd8.pdb |
| AMBP | Wang *et al* | P02760 | 230-339 (110) | 1bik.pdb |
| FKBP1A | Wang *et al* | P62942 | 2-108 (107) | 1bkf.pdb |
| SPTBN1 | Wang *et al* | Q01082 | 173-280 (108) | 1bkr.pdb |
| TIMP2 | Wang *et al* | P16035 | 27-208 (182) | 1br9.pdb |
| ZBTB16 | Wang *et al* | Q05516 | 6-126 (121) | 1buo.pdb |
| LGALS3BP | Wang *et al* | Q08380 | 19-127 (111) | 1by2.pdb |
| HSO90AA1 | Wang *et al* | P07900 | 11-223 (213) | 1byq.pdb |
| CRABP2 | Wang *et al* | P29373 | 2-138 (137) | 1cbs.pdb |
| CD4 | Wang *et al* | P01730 | 26-203 (178) | 1cdy.pdb |
| CALM1 | Wang *et al* | P0DP23 | 5-148 (144) | 1cll.pdb |
| HRAS | Wang *et al* | P01112 | 1-166 (166) | 1ctq.pdb |
| APAF1 | Wang *et al* | O14727 | 1-93 (93) | 1cy5.pdb |
| F5 | Wang *et al* | P12259 | 2066-2224 (159) | 1czt.pdb |
| F8 | Wang *et al* | P00451 | 2190-2348 (159) | 1d7p.pdb |
| ASGR1 | Wang *et al* | P07306 | 154-281 (128) | 1dv8.pdb |
| RNASE1 | Wang *et al* | P07998 | 35-154 (120) | 1e21.pdb |
| CD69 | Wang *et al* | Q07108 | 83-199 (117) | 1e87.pdb |
| PLEKHA1 | Wang *et al* | Q9HB21 | 190-293 (104) | 1eaz.pdb |

| | | | | |
|---|---|---|---|---|
| CCL8 | Wang *et al* | P80075 | 25-99 (75) | 1esr.pdb |
| DAPP1 | Wang *et al* | Q9UN19 | 162-261 (100) | 1fao.pdb |
| PFN1 | Wang *et al* | P07737 | 2-140 (139) | 1fil.pdb |
| AIMP1 | Wang *et al* | Q12904 | 150-313 (164) | 1fl0.pdb |
| FN1 | Wang *et al* | P02751 | 1543-1633 (91) | 1fna.pdb |
| FCGR3B | Wang *et al* | O75015 | 21-193 (173) | 1fnl.pdb |
| IGHE | Wang *et al* | P01854 | 217-424 (208) | 1fp5.pdb |
| GSTZ1 | Wang *et al* | O43708 | 5-212 (208) | 1fw1.pdb |
| SELE | Wang *et al* | P16581 | 22-178 (157) | 1g1t.pdb |
| CST3 | Wang *et al* | P01034* | 36-146 (111) | 1g96.pdb |
| MMP2 | Wang *et al* | P08253 | 461-660 (200) | 1gen.pdb |
| CALML3 | Wang *et al* | P27482 | 5-148 (144) | 1ggz.pdb |
| TXNL1 | Wang *et al* | O43396 | 2-108 (107) | 1gh2.pdb |
| CTSS | Wang *et al* | P25774 | 115-331 (217) | 1glo.pdb |
| GABARAP | Wang *et al* | O95166* | 1-117 (117) | 1gnu.pdb |
| IGF2R | Wang *et al* | P11717 | 1515-1647 (133) | 1gp0.pdb |
| RNASE2 | Wang *et al* | P10153 | 28-161 (134) | 1gqv.pdb |
| COL10A1 | Wang *et al* | Q03692 | 549-680 (132) | 1gr3.pdb |
| MADCAM1 | Wang *et al* | Q13477** | 23-227 (206) | 1gsm.pdb |
| NCF4 | Wang *et al* | Q15080 | 2-144 (143) | 1h6h.pdb |
| BLVRB | Wang *et al* | P30043 | 1-205 (205) | 1hdo.pdb |
| QDPR | Wang *et al* | P09417 | 9-244 (236) | 1hdr.pdb |
| FABP3 | Wang *et al* | P05413 | 2-132 (131) | 1hmt.pdb |
| GSTM2 | Wang *et al* | P28161 | 2-218 (217) | 1hna.pdb |
| MBL2 | Wang *et al* | P11226 | 108-248 (141) | 1hup.pdb |
| IL4 | Wang *et al* | P05112 | 25-153 (129) | 1hzi.pdb |
| PCMT1 | Wang *et al* | P22061 | 3-226 (224) | 1i1n.pdb |
| GTF2F1 | Wang *et al* | P35269 | 449-517 (73) | 1i27.pdb |
| UBR5 | Wang *et al* | O95071 | 2393-2453 (61) | 1i2t.pdb |

| | | | | |
|---|---|---|---|---|
| PRNP | Wang *et al* | P04156** | 119-226 (108) | 1i4m.pdb |
| LPA | Wang *et al* | P08519 | 1274-1355 (82) | 1i71.pdb |
| MMP8 | Wang *et al* | P22894 | 100-262 (163) | 1i76.pdb |
| ICAM1 | Wang *et al* | P05362 | 28-212 (185) | 1iam.pdb |
| ARHGEF1 | Wang *et al* | Q92888* | 44-233 (190) | 1iap.pdb |
| LMNA | Wang *et al* | P02545 | 436-544 (113) | 1ifr.pdb |
| FGF9 | Wang *et al* | P31371 | 52-208 (157) | 1ihk.pdb |
| LCK | Wang *et al* | P06239 | 123-226 (104) | 1ijr.pdb |
| FGF4 | Wang *et al* | P08620 | 79-206 (128) | 1ijt.pdb |
| HSD17B4 | Wang *et al* | P51659 | 622-736 (115) | 1ikt.pdb |
| ABHD14B | Wang *et al* | Q96IU4 | 2-209 (208) | 1imj.pdb |
| UBE2V2 | Wang *et al* | Q15819 | 7-145 (139) | 1j74.pdb |
| ANAPC10 | Wang *et al* | Q9UM13 | 2-162 (161) | 1jhj.pdb |
| MMP12 | Wang *et al* | P39900 | 106-263 (158) | 1jk3.pdb |
| LYZ | Wang *et al* | P61626 | 19-148 (130) | 1jsf.pdb |
| TCL1A | Wang *et al* | P56279 | 4-114 (111) | 1jsg.pdb |
| GGA1 | Wang *et al* | Q9UJY5 | 7-145 (139) | 1jwf.pdb |
| MATK | Wang *et al* | P42679 | 117-213 (97) | 1jwo.pdb |
| PTK2 | Wang *et al* | Q05397 | 908-1049 (142) | 1k04.pdb |
| BCL3 | Wang *et al* | P20749 | 133-360 (228) | 1k1b.pdb |
| ANG | Wang *et al* | P03950 | 26-147 (122) | 1k59.pdb |
| RAP2A | Wang *et al* | P10114 | 1-167 (167) | 1kao.pdb |
| GSN | Wang *et al* | P06396 | 185-288 (104) | 1kcq.pdb |
| NRP1 | Wang *et al* | O14786 | 273-427 (155) | 1kex.pdb |
| DHFR | Wang *et al* | P00374 | 2-187 (186) | 1kmv.pdb |
| HINT1 | Wang *et al* | P49773 | 16-126 (111) | 1kpf.pdb |
| COL6A3 | Wang *et al* | P12111 | 3108-3165 (58) | 1kth.pdb |
| PROCR | Wang *et al* | Q9UNN8 | 25-194 (170) | 1l8j.pdb |
| GNLY | Wang *et al* | P22749 | 63-136 (74) | 1l9l.pdb |

| | | | | |
|---|---|---|---|---|
| CLC | Wang *et al* | Q05315 | 2-142 (141) | 1lcl.pdb |
| B2M | Wang *et al* | P61769** | 21-116 (96) | 1lds.pdb |
| PCTP | Wang *et al* | Q9UKL6 | 8-210 (203) | 1ln1.pdb |
| RBP7 | Wang *et al* | Q96R05 | 2-134 (133) | 1lpj.pdb |
| THBS1 | Wang *et al* | P07996 | 434-546 (113) | 1lsl.pdb |
| RND3 | Wang *et al* | P61587 | 22-200 (179) | 1m7b.pdb |
| TGFBR2 | Wang *et al* | P37173 | 49-153 (105) | 1m9z.pdb |
| SOD1 | Wang *et al* | P00441* | 2-154 (153) | 1mfm.pdb |
| RAC1 | Wang *et al* | P63000 | 2-181 (183) | 1mh1.pdb |
| NT5M | Wang *et al* | Q9NPB1 | 34-227 (194) | 1mh9.pdb |
| SUOX | Wang *et al* | P51687 | 81-160 (80) | 1mj4.pdb |
| APP | Wang *et al* | P05067** | 28-123 (96) | 1mwp.pdb |
| RAB5A | Wang *et al* | P20339 | 15-181 (167) | 1n6h.pdb |
| KIR2DL1 | Wang *et al* | P43626 | 27-221 (195) | 1nkr.pdb |
| FKBP3 | Wang *et al* | Q00688 | 109-224 (116) | 1pbk.pdb |
| CYTH2 | Wang *et al* | Q99418 | 52-246 (195) | 1pbv.pdb |
| PIK3R1 | Wang *et al* | P27986 | 3-85 (83) | 1pht.pdb |
| PLA2GRA | Wang *et al* | P14555 | 21-144 (124) | 1pod.pdb |
| CDC25B | Wang *et al* | P30305 | 388-565 (178) | 1qb0.pdb |
| REG1A | Wang *et al* | P05451 | 23-166 (144) | 1qdd.pdb |
| ESR1 | Wang *et al* | P03372** | 304-551 (248) | 1qkt.pdb |
| ACTN2 | Wang *et al* | P35609 | 391-635 (248) | 1quu.pdb |
| RBP4 | Wang *et al* | P02753 | 19-193 (175) | 1rbp.pdb |
| PLA2G4A | Wang *et al* | P47712 | 17-141 (126) | 1rlw.pdb |
| SPARC | Wang *et al* | P09486 | 153-303 (151) | 1sra.pdb |
| TNC | Wang *et al* | P24821 | 802-891 (90) | 1ten.pdb |
| CLEC3B | Wang *et al* | P05452 | 66-202 (137) | 1tn3.pdb |
| ITGAL | Wang *et al* | P20701 | 153-333 (181) | 1zon.pdb |
| ICAM2 | Wang *et al* | P13598 | 25-216 (192) | 1zxq.pdb |

| | | | | |
|---|---|---|---|---|
| ABL1 | Wang *et al* | P00519 | 57-218 (163) | 2abl.pdb |
| PPIA | Wang *et al* | P62937 | 2-165 (164) | 2cpl.pdb |
| FCGR2B | Wang *et al* | P31994 | 46-218 (173) | 2fcb.pdb |
| FTH1 | Wang *et al* | P02794 | 6-177 (172) | 2fha.pdb |
| IL10 | Wang *et al* | P22301 | 24-178 (155) | 2ilk.pdb |
| S100A7 | Wang *et al* | P31151 | 2-97 (96) | 2psr.pdb |
| TGFB2 | Wang *et al* | P61812 | 303-414 (112) | 2tgi.pdb |
| FGG | Wang *et al* | P02679 | 170-418 (249) | 3fib.pdb |
| CXCL8 | Wang *et al* | P10145 | 32-99 (68) | 3il8.pdb |
| ACP1 | Wang *et al* | P24666 | 2-158 (157) | 5pnt.pdb |
| VIL1 | Fitzkee & Rose | P02640 | 792-826 (36) | 1vii.pdb |
| Prkcd | Fitzkee & Rose | P28867 | 231-280 (50) | 1ptq.pdb |
| spg | Fitzkee & Rose | P06654 | 228-282 (56) | 2gb1.pdb |
| FYN | Fitzkee & Rose | P06241 | 84-142 (59) | 1shfA.pdb |
| cspB | Fitzkee & Rose | P32081 | 1-67 (67) | 1csp.pdb |
| UBC | Fitzkee & Rose | P0CG48 | 609-684 (76) | 1ubq.pdb |
| cI | Fitzkee & Rose | P03034 | 7-93 (87) | 1lmb.pdb |
| Barstar | Fitzkee & Rose | P11540 | 2-90 (89) | 1a19A.pdb |
| ACYP1 | Fitzkee & Rose | P41500 | 4-101 (98) | 2acy.pdb |
| PETE | Fitzkee & Rose | P00299 | 70-168 (99) | 2pcy.pdb |
| CYCS | Fitzkee & Rose | P00004 | 2-105 (104) | 1hrc.pdb |
| Pik3r1 | Fitzkee & Rose | Q63787 | 321-431 (111) | 1fu6A.pdb |
| Hemerythrin | Fitzkee & Rose | P02246 | 1-113 (113) | 2hmqA.pdb |
| LALBA | Fitzkee & Rose | P00711 | 20-141 (122) | 1f6sA.pdb |
| RNASE1 | Fitzkee & Rose | P61823 | 27-150 (124) | 1xptA.pdb |
| cheY | Fitzkee & Rose | P0AE67 | 2-129 (128) | 1ehc.pdb |
| LYZ | Fitzkee & Rose | P00698 | 19-147 (129) | 1hel.pdb |
| Fabp2 | Fitzkee & Rose | P02693 | 2-132 (131) | 1ifb.pdb |
| nuc | Fitzkee & Rose | P00644 | 83-223 (141) | 2sns.pdb |

| | | | | |
|---|---|---|---|---|
| CALM | Fitzkee & Rose | P62157 | 5-147 (143) | 1cm1A.pdb |
| MB | Fitzkee & Rose | P02185 | 2-154 (153) | 1mbo.pdb |
| rnhA | Fitzkee & Rose | P0A7Y4 | 1-155 (155) | 2rn2.pdb |
| gag-pol | Fitzkee & Rose | O92956 | 1331-1487 (162) | 1asu.pdb |
| E (endolysin) | Fitzkee & Rose | P00720 | 1-164 (164) | 2lzm.pdb |
| DFR1 | Fitzkee & Rose | P22906 | 1-192 (192) | 1ai9A.pdb |
| mutY | Fitzkee & Rose | P17802 | 1-225 (225) | 1mun.pdb |
| Triosephosphate isomerase | Fitzkee & Rose | P04789 | 2-250 (249) | 5timA.pdb |
| HAGH | Fitzkee & Rose | Q16775 | 49-308 (260) | 1qh3A.pdb |
| ecoRIR | Fitzkee & Rose | P00642 | 17-277 (261) | 1eriA.pdb |
| galE | Fitzkee & Rose | P09147 | 1-338 (338) | 1nah.pdb |
| CKMT1A | Fitzkee & Rose | P12532 | 39-417 (379) | 1qk1A.pdb |
| PGK1 | Fitzkee & Rose | P00560 | 2-415 (415) | 3pgk.pdb |
| apr | Panja *et al* | P00782 | 108-382 (274) | 1a2q.pdb |
| adk | Panja *et al* | P69441 | 1-214 (214) | 1ake.pdb |
| amy | Panja *et al* | P29957 | 25-472 (448) | 1aqm.pdb |
| hip | Panja *et al* | P00260 | 38-122 (85) | 1b0y.pdb |
| amyE | Panja *et al* | P00691 | 42-466 (425) | 1bag.pdb |
| FGF2 | Panja *et al* | P09038 | 161-285 (125) | 1bas.pdb |
| amyS | Panja *et al* | P06278 | 32-512 (481) | 1bli.pdb |
| axe-2 | Panja *et al* | O59893 | 28-234 (207) | 1bs9.pdb |
| sodB | Panja *et al* | Q9X6W9 | 3-213 (211) | 1coj.pdb |
| fer1 | Panja *et al* | P00217 | 2-129 (128) | 1doi.pdb |
| phnA | Panja *et al* | Q51782 | 2-407 (404) | 1ei6.pdb |
| cyp119 | Panja *et al* | Q55080 | 1-367 (367) | 1f4t.pdb |
| atsA | Panja *et al* | P51691 | 3-527 (524) | 1hdh.pdb |
| hip2 | Panja *et al* | P38524 | 1-71 (71) | 1hpi.pdb |
| katG2 | Panja *et al* | O59651 | 18-731 (707) | 1itk.pdb |
| aspC | Panja *et al* | Q8RR70 | 1-388 (388) | 1j32.pdb |

| | | | | |
|---|---|---|---|---|
| SSO2706 | Panja *et al* | P50389 | 3-236 (226) | 1jds.pdb |
| mtnN | Panja *et al* | P0AF12 | 1-230 (226) | 1jys.pdb |
| rpiA | Panja *et al* | O50083 | 1-229 (229) | 1lk5.pdb |
| VNG_1446H | Panja *et al* | Q9HPW4 | 11-77 (67) | 1mog.pdb |
| speE | Panja *et al* | Q5SK28 | 1-312 (309) | 1uir.pdb |
| Endoglucanase | Panja *et al* | P06564 | 578-761 (181) | 1uww.pdb |
| acyP | Panja *et al* | P84142 | 2-91 (90) | 1v3z.pdb |
| mdh | Panja *et al* | O59028 | 2-360 (337) | 1v9n.pdb |
| serC | Panja *et al* | Q9RME2 | 2-361 (360) | 1w23.pdb |
| amyA | Panja *et al* | Q8GPL8 | 28-515 (488) | 1wza.pdb |
| APE_2278 | Panja *et al* | Q9Y9L0 | 2-245 (240) | 1x0r.pdb |
| mvaS | Panja *et al* | Q9FD71 | 1-383 (383) | 1x9e.pdb |
| Rv1264 | Panja *et al* | P9WMU9 | 14-376 (360) | 1y10.pdb |
| adk | Panja *et al* | P27142 | 1-217 (217) | 1zin.pdb |
| Rv1885c | Panja *et al* | P9WIB9 | 35-199 (165) | 2ao2.pdb |
| ndk | Panja *et al* | P61136 | 4-158 (155) | 2az1.pdb |
| gdh | Panja *et al* | Q977U7 | 1-357 (355) | 2b5v.pdb |
| tdh | Panja *et al* | O58389 | 3-347 (327) | 2d8a.pdb |
| Lysozyme 1 | Panja *et al* | Q7YT16 | 20-141 (122) | 2fbd.pdb |
| Cat-1 | Panja *et al* | Q24940 | 17-326 (306) | 2o6x.pdb |
| oxc | Panja *et al* | P0AFI0 | 5-551 (547) | 2q27.pdb |
| Thioredoxin-dependent peroxiredoxin | Panja *et al* | G1K3P1 | 1-76 (156) | 2xhf.pdb |
| sod | Panja *et al* | Q9Y8H8 | 1-212 (212) | 3ak1.pdb |
| Alkaline serine protease ver112 | Panja *et al* | Q68GV9 | 104-382 (279) | 3f7m.pdb |
| dapE | Panja *et al* | P44514 | 1-376 (370) | 3ic1.pdb |
| pepQ | Panja *et al* | Q44238 | 1-440 (425) | 3l24.pdb |
| sodB | Panja *et al* | P84612 | 1-192 (192) | 3lio.pdb |
| Enpp2 | Panja *et al* | Q9R1E6 | 51-855 (805) | 3nkm.pdb |

| | | | | |
|---|---|---|---|---|
| phoK | Panja *et al* | A1YYW7 | 31-556 (526) | 3q3q.pdb |
| cheC1 | Panja *et al* | Q5V4K4 | 2-206 (200) | 3qta.pdb |
| FOXG_17421 | Panja *et al* | B3A0S5 | 1-327 (327) | 3u7b.pdb |
| Alkaline phosphatase | Panja *et al* | B5BP20 | 31-527 (497) | 3wbh.pdb |
| LGMN | Panja *et al* | Q99538 | 26-288 (267) | 4aw9.pdb |
| LMRG_02624 | Panja *et al* | A0A0H3GD84 | 39-526 (488) | 4cdb.pdb |
| bop | Panja *et al* | Q5UXY6 | 3-238 (236) | 4pxk.pdb |
| mdh | Panja *et al* | A9W386 | 2-320 (319) | 4ror.pdb |
| patA | Panja *et al* | P42588 | 7-459 (453) | 4uox.pdb |
| cysQ | Panja *et al* | P9WKJ1 | 10-267 (266) | 5djf.pdb |
| F | Chen *et al* | P11209 | 480-515 (36) | 1g2cF.pdb |
| HA | Chen *et al* | P03437 | 385-498 (114) | 1htmB.pdb |
| SERPINB14 | Chen *et al* | P01012 | 2-386 (381) | 1jtiB.pdb |
| Plk4 | Chen *et al* | Q64702 | 845-919 (75) | 1mbyA.pdb |
| PVC01_130047600 | Chen *et al* | O60989 | 76-450 (375) | 1miqB.pdb |
| MATALPHA2 | Chen *et al* | P0CY08 | 113-189 (77) | 1mnmC.pdb |
| colG | Chen *et al* | Q9X721 | 1005-1118 (111) | 1nqdA.pdb |
| SRP102 | Chen *et al* | P36057 | 36-244 (191) | 1nrjB.pdb |
| PDE5A | Chen *et al* | O76074 | 535-860 (311) | 1rkpA.pdb |
| cobB | Chen *et al* | P75960 | 40-274 (225) | 1s5pA.pdb |
| F | Chen *et al* | P04849 | 122-183 (62) | 1svfC.pdb |
| SOD1 | Chen *et al* | P00441** | 2-154 (153) | 1uxmK.pdb |
| F | Chen *et al* | O89342 | 143-205 (63) | 1wp8C.pdb |
| S | Chen *et al* | P59594 | 892-981 (124) | 1wyyB.pdb |
| tll0464 | Chen *et al* | Q8DLM0 | 1-112 (102) | 1x0gA.pdb |
| hlyA | Chen *et al* | P09545 | 46-741 (663) | 1xezA.pdb |
| SAR-endolysin | Chen *et al* | Q37875 | 9-185 (170) | 1xjtA.pdb |
| Relb | Chen *et al* | Q04863 | 276-378 (110) | 1zk9A.pdb |
| ftsH | Chen *et al* | Q9WZ49 | 150-606 (421) | 2ce7C.pdb |

| | | | | |
|---|---|---|---|---|
| suhB | Chen *et al* | O33832 | 1-254 (254) | 2p3vA.pdb |
| Polyprotein | Chen *et al* | O36607 | 3-230 (227) | 2pbk.pdb |
| NRP2 | Chen *et al* | O60462 | 276-595 (315) | 2qqjA.pdb |
| MAD2L1 | Chen *et al* | Q13257 | 1-205 (202) | 2vfxL.pdb |
| prgI | Chen *et al* | P41784 | 19-80 (62) | 2x9cA.pdb |
| FN1 | Chen *et al* | P02751 | 516-606 (91) | 3ejhA.pdb |
| CST3 | Chen *et al* | P01034** | 38-146 (107) | 3gaxA.pdb |
| R | Chen *et al* | P27359 | 1-165 (165) | 3hdeA.pdb |
| FBP2 | Chen *et al* | O00757 | 9-337 (326) | 3ifaA.pdb |
| PRIM2 | Chen *et al* | P49643 | 272-457 (167) | 3l9qB.pdb |
| B2M | Chen *et al* | P61769* | 21-119 (99) | 3lowA.pdb |
| gag-pol | Chen *et al* | P04585 | 588-1139 (552) | 3meeA.pdb |
| gp-C | Chen *et al* | Q9ICW1 | 313-422 (103) | 3mkoA.pdb |
| rsmH | Chen *et al* | P60390 | 8-313 (283) | 3tkaA.pdb |
| CWC2 | Chen *et al* | Q12046 | 3-227 (225) | 3tp2A.pdb |
| PR | Chen *et al* | Q3L181 | 1-336 (311) | 3uyiA.pdb |
| macA | Chen *et al* | Q74FY6** | 23-346 (320) | 4aalA.pdb |
| Diphtheria toxin | Chen *et al* | P00588 | 37-567 (499) | 4ae0A.pdb |
| SUN2 | Chen *et al* | Q9UH99 | 522-717 (196) | 4dxrA.pdb |
| bcp | Chen *et al* | Q9YA14 | 2-160 (160) | 4gqcB.pdb |
| PRNP | Chen *et al* | Q95211 | 125-221 (97) | 4hlsA.pdb |
| Grem2 | Chen *et al* | O88273 | 50-160 (111) | 4jphB.pdb |
| pimA | Chen *et al* | A0QWG6 | 1-373 (359) | 4n9wA.pdb |
| plyB | Chen *et al* | Q5W9E8 | 53-519 (465) | 4ov8A.pdb |
| KWL1 | Chen *et al* | P85261 | 48-213 (158) | 4pmkA.pdb |
| COMT | Chen *et al* | P21964 | 54-266 (207) | 4pyiA.pdb |
| MJ1213 | Chen *et al* | Q58610 | 1-109 (109) | 4qhfA.pdb |
| gbs1529 | Chen *et al* | Q8E473 | 494-642 (141) | 4rmbA.pdb |
| OAS1 | Chen *et al* | Q29599 | 1-349 (349) | 4rwnA.pdb |

| | | | | |
|---|---|---|---|---|
| ply | Chen *et al* | Q7ZAK5 | 1-471 (471) | 5aoeB.pdb |
| malE | Chen *et al* | P0AEX9 | 27-393 (402) | 5b3zA.pdb |
| TRAP1 | Chen *et al* | Q12931 | 82-294 (205) | 5f3kA.pdb |
| G | Chen *et al* | P0C2X0 | 1-409 (409) | 5i2mA.pdb |
| DVL2 | Chen *et al* | O14641 | 416-509 (92) | 5suzA.pdb |
| MADCAM1 | membrane protein | Q13477* | 23-231 (209) | 1bqsA.pdb |
| MSN | membrane protein | P26038 | 4-297 (289) | 1ef1A.pdb |
| FCGR2A | membrane protein | P12318 | 37-207 (171) | 1fcgA.pdb |
| SELP | membrane protein | P16109 | 42-199 (158) | 1g1sA.pdb |
| EEA1 | membrane protein | Q15075 | 1289-1411 (123) | 1jocA.pdb |
| GGA1 | membrane protein | Q9UJY5 | 494-639 (146) | 1na8A.pdb |
| SDCBP | membrane protein | O00560 | 197-273 (82) | 1r6jA.pdb |
| CLIC1 | membrane protein | O00299 | 22-234 (213) | 1rk4A.pdb |
| NGF | membrane protein | P01138 | 132-236 (99) | 1sg1A.pdb |
| ANTXR2 | membrane protein | P58335 | 38-218 (181) | 1shuX.pdb |
| IL1RAPL1 | membrane protein | Q9NZN1 | 403-561 (147) | 1t3gA.pdb |
| PGLYRP3 | membrane protein | Q96LB9 | 177-341 (165) | 1twqA.pdb |
| CD3E | membrane protein | P07766 | 33-123 (91) | 1xiwA.pdb |
| CFTR | membrane protein | P13569 | 388-671 (267) | 1xmiA.pdb |
| TRPV2 | membrane protein | Q9Y5S1 | 71-318 (244) | 2f37A.pdb |
| SYNJ2BP | membrane protein | P57105 | 5-98 (100) | 2jikA.pdb |
| GRIP1 | membrane protein | Q9Y3R0 | 148-239 (94) | 2jilA.pdb |
| SELENOS | membrane protein | Q9BQE4 | 52-121 (69) | 2q2fA.pdb |
| CD59 | membrane protein | P13987 | 26-102 (78) | 2uwrA.pdb |
| RAMP2 | membrane protein | O60895 | 58-135 (78) | 2xvtA.pdb |
| ARHGEF1 | membrane protein | Q92888** | 22-233 (165) | 3ab3D.pdb |
| CNKSR2 | membrane protein | Q8WXI2 | 6-80 (74) | 3bs5B.pdb |
| HLA-DRA | membrane protein | P01903 | 28-205 (178) | 3c5jA.pdb |
| AGER | membrane protein | Q15109 | 23-240 (219) | 3cjjA.pdb |

| | | | | |
|---|---|---|---|---|
| IQGAP1 | membrane protein | P46940 | 962-1339 (369) | 3fayA.pdb |
| GRIK1 | membrane protein | P39086 | 445-820 (256) | 3fvoA.pdb |
| ADAM22 | membrane protein | Q9P0K1 | 233-718 (486) | 3g5cA.pdb |
| AQP4 | membrane protein | P55087 | 32-254 (223) | 3gd8A.pdb |
| RHCG | membrane protein | Q9UBD6 | 2-443 (403) | 3hd6A.pdb |
| TRIM72 | membrane protein | Q6ZMU5 | 278-470 (193) | 3kb5A.pdb |
| MPP1 | membrane protein | Q00013 | 282-458 (180) | 3neyA.pdb |
| GLIPR1 | membrane protein | P48060 | 22-214 (193) | 3q2uA.pdb |
| PLXNA2 | membrane protein | O75051 | 1490-1600 (102) | 3q3jA.pdb |
| GORASP2 | membrane protein | Q9H8Y8 | 7-208 (200) | 3rleA.pdb |
| MAPKAP1 | membrane protein | Q9BPZ7 | 372-490 (116) | 3voqA.pdb |
| PILRA | membrane protein | Q9UKJ1 | 32-150 (120) | 3wuzA.pdb |
| macA | membrane protein | Q74FY6* | 24-346 (323) | 4aanA.pdb |
| PMP2 | membrane protein | P02689 | 1-132 (132) | 4bvmA.pdb |
| DYSF | membrane protein | O75923 | 1-124 (127)<br>943-1051 (109) | 4iqhA.pdb<br>4caiA.pdb |
| BECN1 | membrane protein | Q14457 | 248-447 (195) | 4ddpA.pdb |
| STING1 | membrane protein | Q86WV6 | 155-337 (173) | 4emtA.pdb |
| DLG1 | membrane protein | Q12959 | 310-406 (97) | 4g69A.pdb |
| FOLR1 | membrane protein | P15328 | 30-233 (206) | 4km6A.pdb |
| SLC4A1 | membrane protein | P02730 | 57-350 (276) | 4ky9A.pdb |
| MR1 | membrane protein | Q95460 | 23-291 (262) | 4l4vA.pdb |
| HLA-B | membrane protein | P01889 | 25-298 (274) | 4lcyA.pdb |
| PRNP | membrane protein | P04156** | 118-224 (107) | 4n9oA.pdb |
| ESYT2 | membrane protein | A0FGR8 | 363-659 (292) | 4npjA.pdb |
| PVDR | membrane protein | P22290 | 211-508 (282) | 4nuuA.pdb |
| LGR4 - fusion | membrane protein | Q9BXB1 | 27-399 (443) | 4qxeA.pdb |
| PLK1 | membrane protein | P53350 | 372-599 (223) | 4rcpA.pdb |
| TOR1AIP1 | membrane protein | Q5JTV8 | 360-583 (224) | 4tvsA.pdb |
| VAMP8 | membrane protein | Q9BV40 | 11-74 (64) | 4wy4A.pdb |

| | | | | |
|---|---|---|---|---|
| PGRMC1 | membrane protein | O00264 | 72-179 (112) | 4x8yA.pdb |
| GPC1 | membrane protein | P35052 | 25-473 (411) | 4ywtA.pdb |
| GLP1R | membrane protein | P43220 | 29-128 (100) | 5e94H.pdb |
| SCN2B | membrane protein | O60939 | 30-148 (122) | 5febA.pdb |
| ADORA2A - fusion | membrane protein | P29274 | 2-305 (387) | 5iu4A.pdb |
| ADIPOR2 | membrane protein | Q86V24 | 99-380 (282) | 5lx9A.pdb |
| ZMPSTE24 | membrane protein | O75844 | 10-474 (444) | 5sytA.pdb |
| PTGES | membrane protein | O14684 | 5-152 (147) | 5tl9A.pdb |
| CHRM2 - fusion | membrane protein | P08172 | 16-458 (384) | 5zkcA.pdb |
| SLMAP | membrane protein | Q14BN4 | 2-135 (134) | 6ar2A.pdb |
| FZD4 - fusion | membrane protein | Q9ULV1 | 181-513 (379) | 6bd4A.pdb |
| C5AR1 - fusion | membrane protein | P21730 | 30-327 (370) | 6c1rB.pdb |
| GRM5 - fusion | membrane protein | P41594 | 569-836 (409) | 6ffiA.pdb |
| CCDC90B - fusion | membrane protein | Q9GZT6 | 62-126 (94) | 6h9mA.pdb |
| TACR1 - fusion | membrane protein | P25103 | 27-327 (483) | 6hlpA.pdb |
| MCOLN2 | membrane protein | Q8IZK6 | 92-282 (173) | 6hrrA.pdb |
| KDELR2 | membrane protein | Q5ZKX9 | 1-207 (207) | 6i6hA.pdb |
| DHODH | membrane protein | Q02127 | 29-395 (367) | 6idjA.pdb |
| MPLZL1 | membrane protein | O95297 | 38-158 (119) | 6igwA.pdb |
| MFN2 | membrane protein | O95140 | 24-418 (428) | 6jfkA.pdb |
| GPR52 - fusion | membrane protein | Q9Y2T5 | 21-338 (441) | 6li0A.pdb |
| AQP7 | membrane protein | O14520 | 33-279 (247) | 6qziA.pdb |
| LTC4S | membrane protein | Q16873 | 2-144 (143) | 6r7dA.pdb |
| PTCH1 | membrane protein | Q13635 | 149-423 (277)<br>842-935 (94) | 6rtwA.pdb<br>6rvcA.pdb |
| ERVW-1 | membrane protein | Q9UQF0 | 345-433 (89) | 6rx1A.pdb |
| ERVFRD-1 | membrane protein | P60508 | 380-468 (89) | 6rx3A.pdb |
| CYSLTR2 - fusion | membrane protein | Q9NS75 | 29-322 (365) | 6rz6A.pdb |
| SLC2A1 | membrane protein | P11166 | 8-455 (448) | 6thaA.pdb |
| HCRTR1 | membrane protein | O43613 | 45-346 (301) | 6todA.pdb |

| | | | | |
|---|---|---|---|---|
| KCNMA1 | membrane protein | Q12791 | 408-1121 (594) | 6v5aA.pdb |
| SCN4B | membrane protein | Q8IWT1 | 37-154 (115) | 6vsvA.pdb |
| JAGN1 - fusion | membrane protein | Q8N5M9 | 2-183 (397) | 6wvdA.pdb |
| malE - fusion | membrane protein | P0AEX9 | 26-392 (571) | 6zhoA.pdb |
| DDR2 - fusion | membrane protein | Q16832 | 561-849 (275) | 7aymA.pdb |
| GABARAP - fusion | membrane protein | O95166** | 1-116 (132) | 7brqA.pdb |

[a] The Protein Data Bank (1) was used to identify folded regions within proteins. Originally, we searched for folded regions within proteins known to exhibit phase separation behavior, finding 82 folded regions (2). The phase-separating proteins were obtained from lists compiled by Vernon et al (3), the PhaSePro database (4), and the DisProt database (5). A complete list of these 82 folded regions has been published elsewhere (2). To that list, we added folded regions from 122 human proteins with nonhomologous structures obtained from Wang et al (6), 32 proteins with small to large structures obtained from Fitzkee and Rose (7), 54 extremophile proteins obtained from Panja et al (8), 53 metamorphic proteins obtained from Chen et al (9), and 90 membrane proteins that were found by searching the Protein Data Bank for the phrase "membrane protein." Duplicate entries were removed from the combined list. For example, human Galectin-3 (UniProt accession number P17931) is found in both the PhaSePro database of phase-separating proteins and the list of human proteins with nonhomologous structures from Wang et al. Duplicate entries in the combined list are identified by an asterisk at the end of the UniProt accession number; two asterisks indicate the duplicate that was removed from the final folded set. Protein names with "- fusion" indicate a protein that is fused to another protein in the crystallographic structure, which is found among a few classified as "membrane protein".

[b] Residue positions with resolved atomic coordinates in a PDB structure (x-ray or NMR) were used to verify regions ($N \geq 20$) that fold. Unresolved residues were not included in folded regions. Protein sequences were extracted from the referenced PDB file and thus may contain substitutions, deletions, and/or insertions (excluding histidine affinity tags) compared to the UniProt sequence. The value of $N$ in parenthesis is the length of the extracted sequence.

**Table S2. Summary of mean $v_{model}$ in the ID and folded sequence subsets.**

| Set | Number | $v_{model}$ [a] | *t*-test [b] | *U*-test [b] |
|---|---|---|---|---|
| Previous ID | 23 | 0.558 ± 0.019 | - | - |
| *BMRB & DisProt* | 98 | 0.558 ± 0.023 | 0.44 | 0.48 |
| Previous Folded | 82 | 0.536 ± 0.008 | - | - |
| *Human* | 122 | 0.536 ± 0.007 | 0.40 | 0.32 |
| *Small-to-large* | 32 | 0.537 ± 0.009 | 0.36 | 0.41 |
| *Extremophile* | 54 | 0.542 ± 0.011 | $1.2e^{-4}$ | $2.4e^{-4}$ |
| *Membrane* | 90 | 0.537 ± 0.006 | 0.17 | 0.21 |
| *Metamorphic* | 53 | 0.537 ± 0.006 | 0.15 | 0.18 |

[a] Mean ± standard deviation.

[b] One-tail *p*-value, where values $<0.05$ indicate the compared sets are statistically different in their means. Comparisons are to the previous set; BMRB & DisProt to the Previous ID, and Human, Small-to-large, Extremophile, Membrane, and Metamorphic to the Previous Folded.

**Table S3. Summary of mean β-turn propensity in the ID and folded sequence subsets.**

| Set | Number | β-turn propensity [a] | *t*-test [b] | *U*-test [b] |
|---|---|---|---|---|
| Previous ID | 23 | 1.062 ± 0.082 | - | - |
| *BMRB & DisProt* | 98 | 1.110 ± 0.071 | 6.5e$^{-3}$ | 9.3e$^{-4}$ |
| Previous Folded Set | 82 | 0.969 ± 0.039 | - | - |
| *Human* | 122 | 0.980 ± 0.039 | 0.03 | 0.07 |
| *Small-to-large* | 32 | 0.968 ± 0.027 | 0.42 | 0.34 |
| *Extremophile* | 54 | 0.983 ± 0.030 | 0.01 | 0.03 |
| *Membrane* | 90 | 0.956 ± 0.046 | 0.02 | 0.02 |
| *Metamorphic* | 53 | 0.972 ± 0.040 | 0.30 | 0.48 |

[a] Mean ± standard deviation.
[b] One-tail *p*-value, where values <0.05 indicate the compared sets are statistically different in their means. Comparisons are to the previous set; BMRB & DisProt to the Previous ID, and Human, Small-to-large, Extremophile, Membrane, and Metamorphic to the Previous Folded.

**Table S4. List of IDRs not known to exhibit phase separation behavior.**

| Name | Database [a] | Entry number | UniProt accession number | ID region ($N$) |
|---|---|---|---|---|
| pknG | BMRB | 26027 | P9WI73 | 1-75 (75) |
| HCK | BMRB | 27554 | P08631 | 2-79 |
| SIC1 | BMRB | 16657 | P38634 | 1-90 (90) |
| SLC9A1 | BMRB | 26557 | P19634 | 680-815 (136) |
| ERD14 | BMRB | 16876 | P42763 | 1-185 (185) |
| Spp1 | DisProt | DP01448 | P10923 | 17-294 (278) |
| PAGE4 | DisProt | DP01435 | O60829 | 1-102 (102) |
| MAP2K4 | DisProt | DP01400 | P45985 | 1-86 (86) |
| Sufu | DisProt | DP01397 | Q9Z0P7 | 279-359 (81) |
| HCN1 | DisProt | DP01317 | O60741 | 1-93 (93) |
| SUFU | DisProt | DP01312 | Q9UMX1 | 279-360 (82) |
| PQBP1 | DisProt | DP01308 | O60828 | 82-265 (184) |
| HIRD11 | DisProt | DP01300 | Q9SLJ2 | 1-98 (98) |
| LEA18 | DisProt | DP01299 | Q96273 | 1-97 (97) |
| PSEN1 | DisProt | DP01292 | P49768 | 1-77 (77) |
| Prothymosin a14 | DisProt | DP01228 | Q9UMZ1 | 1-101 (101) |
| Ppp1r10 | DisProt | DP01202 | O55000 | 309-433 (125) |
| NOLC1 | DisProt | DP01178 | Q14978 | 1-699 (699) |
| Gja4 | DisProt | DP01175 | P28235 | 233-333 (101) |
| DCLRE1C | DisProt | DP01162 | Q96SD1 | 480-575 (96) |
| ptkA | DisProt | DP01160 | P9WPI9 | 1-81 (81) |
| H1-0 | DisProt | DP01156 | P07305 | 105-194 (90) |
| CHZ1 | DisProt | DP01135 | P40019 | 1-153 (153) |
| Caskin1 | DisProt | DP01127 | Q8VHK2 | 603-1430 (828) |
| Ttn-1 | DisProt | DP01090 | A0A2I2LG13 | 2793-6678 (3886) |
| PM28 | DisProt | DP01088 | Q9XES8 | 1-89 (89) |

| | | | | |
|---|---|---|---|---|
| YRB2 | DisProt | DP01079 | P40517 | 1-203 (203) |
| Ahn-1 | DisProt | DP01074 | Q7YUB9 | 1-86 (86) |
| MSA2 | DisProt | DP01067 | P19599 | 21-238 (218) |
| LMP2A | DisProt | DP01060 | A8CDV5 | 1-118 (118) |
| Omega gliadin storage protein | DisProt | DP01040 | Q9FUW7 | 1-280 (280) |
| SLE2 | DisProt | DP01036 | I1JLC8 | 1-105 (105) |
| pscP | DisProt | DP00993 | Q9I332 | 1-253 (253) |
| Small delta antigen | DisProt | DP00965 | P0C6L3 | 60-195 (136) |
| SBDS-like protein | DisProt | DP00957 | C0J347 | 264-464 (201) |
| GAP43 | DisProt | DP00955 | P06836 | 1-242 (242) |
| N | DisProt | DP00948 | P59595 | 182-259 (78) |
| Ppp1r9b | DisProt | DP00943 | O35274 | 1-154 (154) |
| BASP1 | DisProt | DP00930 | P80723 | 1-227 (227) |
| NABP2 | DisProt | DP00864 | Q9BQ15 | 110-211 (102) |
| trm10 | DisProt | DP00798 | O14214 | 1-83 (83) |
| CNGB1 | DisProt | DP00768 | Q28181-4 | 14-99 (86)<br>272-590 (319) |
| Smtnl1 | DisProt | DP00742 | Q99LM3 | 1-341 (341) |
| dre4 | DisProt | DP00721 | Q8IRG6 | 889-1044 (156) |
| Ssrp | DisProt | DP00720 | Q05344 | 437-554 (118)<br>625-723 (99) |
| N | DisProt | DP00698 | O89339 | 400-532 (133) |
| RYBP | DisProt | DP00694 | Q8N488 | 1-228 (228) |
| L1CAM | DisProt | DP00666 | P32004 | 1144-1257 (114) |
| GMPM1 | DisProt | DP00664 | Q01417 | 1-173 (173) |
| ALB3 | DisProt | DP00662 | Q8LBP4 | 339-462 (124) |
| MAC-41A | DisProt | DP00659 | P16458 | 233-385 (153) |
| COR47 | DisProt | DP00657 | P31168 | 1-265 (265) |
| N | DisProt | DP00640 | Q89933 | 400-525 (126) |
| ERD10 | DisProt | DP00606 | P42759 | 1-260 (260) |
| Genome polyprotein | DisProt | DP00588 | P27958 | 1-82 (82) |

| | | | | |
|---|---|---|---|---|
| stm | DisProt | DP00584 | A2VD23 | 1-613 (613) |
| SEPTIN4 | DisProt | DP00537 | O43236 | 1-119 (119) |
| DHN1 | DisProt | DP00530 | P12950 | 1-168 (168) |
| MYOM1 | DisProt | DP00517 | P52179 | 836-931 (96) |
| NUPR1 | DisProt | DP00510 | O60356 | 1-82 (82) |
| UBA2 | DisProt | DP00486 | Q9UBT2 | 551-640 (90) |
| HY5 | DisProt | DP00469 | O24646 | 1-77 (77) |
| cna | DisProt | DP00461 | P08083 | 1-90 (90) |
| Chm | DisProt | DP00458 | P37727 | 108-208 (101) |
| PPP1R1B | DisProt | DP00421 | P07516 | 1-202 (202) |
| JAG1 | DisProt | DP00418 | P78504 | 1094-1218 (125) |
| URE1 | DisProt | DP00353 | P23202 | 1-90 (90) |
| DNAJC6 | DisProt | DP00351 | Q27974 | 547-813 (267) |
| col | DisProt | DP00342 | P09883 | 1-83 (83) |
| Trl | DisProt | DP00328 | Q08605 | 368-444 (77) |
| PPP1R1A | DisProt | DP00325 | P01099 | 1-166 (166) |
| ADD2 | DisProt | DP00241 | P35612 | 409-726 (318) |
| ADD1 | DisProt | DP00240 | P35611 | 430-737 (308) |
| SSB | DisProt | DP00229 | P05455 | 326-408 (83) |
| Nucleoplasmin | DisProt | DP00217 | P05221 | 120-200 (81) |
| CAST | DisProt | DP00196 | P20810 | 137-277 (141) |
| HMGN2 | DisProt | DP00195 | P02313 | 1-89 (89) |
| Late embryogenesis abundant protein 1 | DisProt | DP00186 | Q95V77 | 1-143 (143) |
| CTDP1 | DisProt | DP00177 | Q9Y5B0 | 879-961 (83) |
| TCF7L2 | DisProt | DP00175 | Q9NQB0 | 1-130 (130) |
| zipA | DisProt | DP00161 | P77173 | 86-185 (100) |
| RAD23A | DisProt | DP00156 | P54725 | 79-160 (82) |
| NEFL | DisProt | DP00151 | P02547 | 444-549 (106) |
| Slbp | DisProt | DP00144 | Q9VAN6 | 97-175 (79) |

| | | | | |
|---|---|---|---|---|
| PTHLH | DisProt | DP00138 | P12272 | 68-144 (77) |
| H1-4 | DisProt | DP00136 | P15865 | 1-217 (217) |
| PRB4 | DisProt | DP00119 | P10163 | 17-310 (294) |
| Desiccation-related protein clone PCC6-19 | DisProt | DP00112 | P22239 | 1-155 (155) |
| H1-0 | DisProt | DP00097 | P10922 | 96-193 (98) |
| TOP2 | DisProt | DP00076 | P06786 | 1178-1428 (251) |
| TOP1 | DisProt | DP00075 | P11387 | 1-214 (214) |
| Structural polyprotein | DisProt | DP03350 | P03316 | 1-113 (113) |
| RPA1 | DisProt | DP00061 | P27694 | 105-180 (76) |
| HMGA1 | DisProt | DP00040 | P17096 | 1-107 (107) |
| HMGN2 | DisProt | DP00039 | P05204 | 1-90 (90) |
| RAP1 | DisProt | DP00020 | P11938 | 1-123 (123) |

[a] The Biological Magnetic Resonance Data Bank (BMRB) (10) and DisProt (5) databases were used to identify IDRs that are not known to exhibit phase separation behavior. This list of verified IDRs, wherein duplicates have been removed, was combined with a list of 23 IDRs that have been identified and reported elsewhere (2).

**Table S5. Enthalpy, entropy, and free energy of phase separation of A1-LCD and Ddx4 mutants.**

| IDR | Mutant | Primary Sequence | $\Delta h^{\circ\,a}$ | $\Delta s^{\circ}/R^{\,b}$ | $\Delta g^{\circ\,c}$ |
|---|---|---|---|---|---|
| Ddx4 | CS | MGDRDWRAEINPHMSSYVPIFEKDRYSGENGRNFNDTP<br>ASSSEMRDGPSERDHFMKSGFASGDNFGNRDAGKCNER<br>DNTSTMGGFGVGKSFGNEGFSNSRFERGDSSGFWRESS<br>NDCRDNPTRNDGFSDRGGYEKGNNSEASGPYERGGRGS<br>FDGCRGGFGLGSPNNRLDPRECMQRTGGLFGSDRPVLS<br>GTGNGDTSQSRSGSGSERGGYKGLNEKVITGSGENSWK<br>SEARGGES | -23.09 | 38.82 | -44.16 |
| Ddx4 | WT | MGDEDWEAEINPHMSSYVPIFEKDRYSGENGDNFNRTP<br>ASSSEMDDGPSRRDHFMKSGFASGRNFGNRDAGECNKR<br>DNTSTMGGFGVGKSFGNRGFSNSRFEDGDSSGFWRESS<br>NDCEDNPTRNRGFSKRGGYRDGNNSEASGPYRRGGRGS<br>FRGCRGGFGLGSPNNDLDPDECMQRTGGLFGSRRPVLS<br>GTGNGDTSQSRSGSGSERGGYKGLNEEVITGSGKNSWK<br>SEAEGGES | -5.43 | 8.47 | -10.03 |
| A1-LCD | Aro+ | GSMAFASSFQRGRYGSGNFGGGRGGGFGGNDNFGRGGN<br>FSGRGGFGGSRGGGGYGGSGDGYNGFGNDGSNFGGGGS<br>YNDFGNYNNQSSNFGPMKGGNFGGRSSGGSYGGGQYFA<br>KPRNQGGYGGSSFSSSYGSGRRF | -30.58 | 43.50 | -54.18 |
| A1-LCD | Aro- | GSMASASSSQRGRSGSGNSGGGRGGGFGGNDNFGRGGN<br>SSGRGGFGGSRGGGGYGGSGDGYNGFGNDGSNSGGGGS<br>SNDFGNYNNQSSNFGPMKGGNFGGRSSGGSGGGGQYSA<br>KPRNQGGYGGSSSSSSSGSGRRF | -17.44 | 27.00 | -32.10 |
| A1-LCD | -12F+12Y | GSMASASSSQRGRSGSGNYGGGRGGGYGGNDNYGRGGN<br>YSGRGGYGGSRGGGGYGGSGDGYNGYGNDGSNYGGGGS<br>YNDYGNYNNQSSNYGPMKGGNYGGRSSGGSGGGGQYYA<br>KPRNQGGYGGSSSSSSYGSGRRY | -27.55 | 40.89 | -49.74 |
| A1-LCD | -9F+6Y | GSMASASSSQRGRSGSGNFGGGRGGGYGGNDNYGRGGN<br>YSGRGGFGGSRGGGGYGGSGDGYNGGGNDGSNYGGGGS<br>YNDSGNYNNQSSNFGPMKGGNYGGRSSGGSGGGGQYGA<br>KPRNQGGYGGSSSSSSYGSGRRY | -25.16 | 38.78 | -46.21 |
| A1-LCD | -4D | GSMASASSSQRGRSGSGNFGGGRGGGFGGNGNFGRGGN<br>FSGRGGFGGSRGGGGYGGSGGGYNGFGNSGSNFGGGGS<br>YNGFGNYNNQSSNFGPMKGGNFGGRSSGPYGGGGQYFA<br>KPRNQGGYGGSSSSSSYGSGRRF | -25.05 | 39.72 | -46.61 |
| A1-LCD | -9F+3Y | GSMASASSSQRGRSGSGNFGGGRGGGYGGNDNGGRGGN<br>YSGRGGFGGSRGGGGYGGSGDGYNGGGNDGSNYGGGGS<br>YNDSGNGNNQSSNFGPMKGGNYGGRSSGGSGGGGQYGA<br>KPRNQGGYGGSSSSSSYGSGRRS | -24.51 | 38.89 | -45.62 |
| A1-LCD | -6R+6K | GSMASASSSQKGKSGSGNFGGGRGGGFGGNDNFGKGGN<br>FSGRGGFGGSKGGGGYGGSGDGYNGFGNDGSNFGGGGS<br>YNDFGNYNNQSSNFGPMKGGNFGGKSSGGSGGGGQYFA<br>KPRNQGGYGGSSSSSSYGSGRKF | -24.29 | 39.95 | -45.98 |
| A1-LCD | -8F+4Y | GSMASASSSQRGRSGSGNFGGGRGGGYGGNDNGGRGGN<br>YSGRGGFGGSRGGGGYGGSGDGYNGGGNDGSNYGGGGS<br>YNDSGNYNNQSSNFGPMKGGNYGGRSSGGSGGGGQYGA<br>KPRNQGGYGGSSSSSSYGSGRRF | -23.91 | 37.37 | -44.19 |
| A1-LCD | +7R+12D | GSMASADSSQRDRDDRGNFGDGRGGGFGGNDNFGRGGN<br>FSDRGGFGGSRGDGRYGGDGDRYNGFGNDGRNFGGGGS<br>YNDFGNYNNQSSNFDPMKGGNFRDRSSGPYDRGGQYFA<br>KPRNQGGYGGSSSSRSYGSDRRF | -22.45 | 30.44 | -38.97 |
| A1-LCD | +2R | GSMASASSSQRGRSGSGNFGGGRGGGFGGNDNFGRGGN<br>FSGRGGFGGSRGGGGYGGSGDGYNGFRNDGSNFGGGGR<br>YNDFGNYNNQSSNFGPMKGGNFGGRSSGPYGGGGQYFA<br>KPRNQGGYGGSSSSSSYGSGRRF | -21.74 | 31.96 | -39.09 |
| A1-LCD | -2R-2K+3D | GSMASASSSQDGRSGSGNFGGGRGGGFGGNDNFGRGGN<br>FSGRGGFGGSRGGGGYGGSGDGYNGFGNDGSNFGGGGS<br>YNDFGNYNNQSSNFGPMDGGNFGGRSSGPYGGGGQYFA<br>DPRNQGGYGGSSSSSSYGSGGRF | -20.92 | 29.85 | -37.12 |

| | | | | | |
|---|---|---|---|---|---|
| A1-LCD | WT+NLS | GSMASASSSQRGRSGSGNFGGGRGGGFGGNDNFGRGGN<br>FSGRGGFGGSRGGGGYGGSGDGYNGFGNDGSNFGGGGS<br>YNDFGNYNNQSSNFGPMKGGNFGGRSSGPYGGGGQYFA<br>KPRNQGGYGGSSSSSSYGSGRRF | -20.27 | 28.79 | -35.90 |
| A1-LCD | +8D | GSMASASSSQRDRSGSGNFGGGRDGGFGGNDNFGRGDN<br>FSGRGDFGGSRDGGGYGGSGDGYNGFGNDGSNFGGGGS<br>YNDFGNYNNQSSNFGPMKGGNFGGRSSDPYGGGGQYFA<br>KPRNQDGYGGSSSSSSYDSGRRF | -20.27 | 29.50 | -36.28 |
| A1-LCD | WT | GSMASASSSQRGRSGSGNFGGGRGGGFGGNDNFGRGGN<br>FSGRGGFGGSRGGGGYGGSGDGYNGFGNDGSNFGGGGS<br>YNDFGNYNNQSSNFGPMKGGNFGGRSSGGSGGGGQYFA<br>KPRNQGGYGGSSSSSSYGSGRRF | -20.22 | 28.91 | -35.91 |
| A1-LCD | -3R+3K | GSMASASSSQRGKSGSGNFGGGRGGGFGGNDNFGRGGN<br>FSGRGGFGGSKGGGGYGGSGDGYNGFGNDGSNFGGGGS<br>YNDFGNYNNQSSNFGPMKGGNFGGRSSGGSGGGGQYFA<br>KPRNQGGYGGSSSSSSYGSGRKF | -19.95 | 30.44 | -36.47 |
| A1-LCD | -2K | GSMASASSSQRGRSGSGNFGGGRGGGFGGNDNFGRGGN<br>FSGRGGFGGSRGGGGYGGSGDGYNGFGNDGSNFGGGGS<br>YNDFGNYNNQSSNFGPMGGGNFGGRSSGPYGGGGQYFA<br>GPRNQGGYGGSSSSSSYGSGRRF | -19.62 | 26.09 | -33.78 |
| A1-LCD | +4D | GSMASASSSQRDRSGSGNFGGGRGGGFGGNDNFGRGGN<br>FSGRGDFGGSRGGGGYGGSGDGYNGFGNDGSNFGGGGS<br>YNDFGNYNNQSSNFGPMKGGNFGGRSSDPYGGGGQYFA<br>KPRNQGGYGGSSSSSSYDSGRRF | -17.88 | 23.62 | -30.70 |
| A1-LCD | +7K+12D | GSMASADSSQRDRDDKGNFGDGRGGGFGGNDNFGRGGN<br>FSDRGGFGGSRGDGKYGGDGDKYNGFGNDGKNFGGGGS<br>YNDFGNYNNQSSNFDPMKGGNFKDRSSGPYDKGGQYFA<br>KPRNQGGYGGSSSSKSYGSDRRF | -17.55 | 25.50 | -31.39 |
| A1-LCD | +12D | GSMASADSSQRDRDDSGNFGDGRGGGFGGNDNFGRGGN<br>FSDRGGFGGSRGDGGYGGDGDGYNGFGNDGSNFGGGGS<br>YNDFGNYNNQSSNFDPMKGGNFGDRSSGPYDGGGQYFA<br>KPRNQGGYGGSSSSSSYGSDRRF | -17.01 | 25.15 | -30.66 |
| A1-LCD | -6R | GSMASASSSQGGRSGSGNFGGGRGGGFGGNDNFGGGGN<br>FSGSGGFGGSRGGGGYGGSGDGYNGFGNDGSNFGGGGS<br>YNDFGNYNNQSSNFGPMKGGNFGGSSSGPYGGGGQYFA<br>KPGNQGGYGGSSSSSSYGSGGRF | -16.90 | 22.33 | -29.02 |
| A1-LCD | +7F-7Y | GSMASASSSQRGRSGSGNFGGGRGGGFGGNDNFGRGGN<br>FSGRGGFGGSRGGGGFGGSGDGFNGFGNDGSNFGGGGS<br>FNDFGNFNNQSSNFGPMKGGNFGGRSSGGSGGGGQFFA<br>KPRNQGGFGGSSSSSSFGSGRRF | -16.47 | 23.39 | -29.17 |
| A1-LCD | +12E | GSMASAESSQREREESGNFGEGRGGGFGGNDNFGRGGN<br>FSERGGFGGSRGEGGYGGEGDGYNGFGNDGSNFGGGGS<br>YNDFGNYNNQSSNFEPMKGGNFGERSSGPYEGGGQYFA<br>KPRNQGGYGGSSSSSSYGSERRF | -15.76 | 23.62 | -28.58 |
| A1-LCD | +7R+10D | GSMASADSSQRDRDGRGNFGDGRGGGFGGNDNFGRGGN<br>FSDRGGFGGSRGGGRYGGDGDRYNGFGNDGRNFGGGGS<br>YNDFGNYNNQSSNFDPMKGGNFRDRSSGPYDRGGQYFA<br>KPRNQGGYGGSSSSRSYGSDRRF | -14.18 | 17.05 | -23.43 |
| A1-LCD | -10R | GSMASASSSQGGSSGSGNFGGGGGGGFGGNDNFGGGGN<br>FSGSGGFGGSGGGGGYGGSGDGYNGFGNDGSNFGGGGS<br>YNDFGNYNNQSSNFGPMKGGNFGGSSSGPYGGGGQYFA<br>KPGNQGGYGGSSSSSSYGSGGGF | -13.53 | 19.16 | -23.93 |
| A1-LCD | -4R-2K+5D | GSMASASSSQDGRSGSGNFGGGDGGGFGGNDNFGRGGN<br>FSGGGGFGGSRGGGGYGGSGDGYNGFGNDGSNFGGGGS<br>YNDFGNYNNQSSNFGPMDGGNFGGRSSGPYGGGGQYFA<br>DPRNQGGYGGSSSSSSYGSGDRF | -12.83 | 17.63 | -22.40 |

[a] Standard molar enthalpy ($\Delta h°$) in units of kcal/mol. Values for Ddx4 CS, Ddx4 WT, A1-LCD Aro+, and A1-LCD Aro- were calculated from the temperature dependence to $c_{sat}$ (see Methods) using $c_{sat}$ values digitally extracted from Figures 1C-D in Brady et al (11) and Figure 3F in Martin et al (12). Values for all other IDRs in this table were digitally extracted from Supplementary Figure 7D in Bremer et al (13).

[b] Standard molar entropy ($\Delta s°$) divided by the universal gas constant, and thus dimensionless. Values for Ddx4 CS, Ddx4 WT, A1-LCD Aro+, and A1-LCD Aro- were calculated from the temperature dependence to $c_{sat}$ (see Methods) using $c_{sat}$ values digitally extracted from Figures 1C-D in Brady et al (11) and Figure 3F in Martin et al (12). Values for all other IDRs in this table were digitally extracted from Supplementary Figure 7E in Bremer et al (13).
[c] Standard molar free energy ($\Delta g°$) in units of kcal/mol. Values were calculated from $\Delta h°$ and $\Delta s°$ using the equation, $\Delta g° = \Delta h° - T\Delta s°$, where $T$ is the standard temperature (273.15 K).

**Table S6. Saturation concentration (at 4 °C) of A1-LCD mutants.**

| Mutant | Primary Sequence | $c_{sat}$ [a] |
|---|---|---|
| +23G-23S-12F+12Y | GSMAGAGGGQRGRGGGGNYGGGRGGGYGGNDNYGRGGNYGGRGGYGGGRG<br>GGGYGGGGDGYNGYGNDGGNYGGGGGYNDYGNYNNQGGNYGPMKGGNYGG<br>RGGGGGGGGGQYYAKPRNQGGYGGGGGGGGYGGGRRY | 4.86E-07 |
| +7R+12D | GSMASADSSQRDRDDRGNFGDGRGGGFGGNDNFGRGGNFSDRGGFGGSRG<br>DGRYGGDGDRYNGFGNDGRNFGGGGSYNDFGNYNNQSSNFDPMKGGNFRD<br>RSSGPYDRGGQYFAKPRNQGGYGGSSSSRSYGSDRRF | 7.49E-07 |
| -12F+12Y | GSMASASSSQRGRSGSGNYGGGRGGGYGGNDNYGRGGNYSGRGGYGGSRG<br>GGGYGGSGDGYNGYGNDGSNYGGGGSYNDYGNYNNQSSNYGPMKGGNYGG<br>RSSGGSGGGGQYYAKPRNQGGYGGSSSSSSYGSGRRY | 2.74E-06 |
| +4D | GSMASASSSQRDRSGSGNFGGGRGGGFGGNDNFGRGGNFSGRGDFGGSRG<br>GGGYGGSGDGYNGFGNDGSNFGGGGSYNDFGNYNNQSSNFGPMKGGNFGG<br>RSSDPYGGGGQYFAKPRNQGGYGGSSSSSSYDSGRRF | 4.04E-06 |
| -6R | GSMASASSSQGGRSGSGNFGGGRGGGFGGNDNFGGGGNFSGSGGFGGSRG<br>GGGYGGSGDGYNGFGNDGSNFGGGGSYNDFGNYNNQSSNFGPMKGGNFGG<br>SSSGPYGGGGQYFAKPGNQGGYGGSSSSSSYGSGGRF | 7.34E-06 |
| -2R-2K+3D | GSMASASSSQDGRSGSGNFGGGRGGGFGGNDNFGRGGNFSGRGGFGGSRG<br>GGGYGGSGDGYNGFGNDGSNFGGGGSYNDFGNYNNQSSNFGPMDGGNFGG<br>RSSGPYGGGGQYFADPRNQGGYGGSSSSSSYGSGGRF | 9.91E-06 |
| -20G+20S-12F+12Y | GSMASASSSQRSRSGSGNYSGSRSGSYSGNDNYGRSGNYSGRSGYGGSRS<br>GGGYSGSGDSYNSYGNDGSNYSGSGSYNDYGNYNNQSSNYGPMKSGNYGG<br>RSSGSSGGSGQYYAKPRNQGSYSGSSSSSSYGSSRRY | 1.22E-05 |
| WT+NLS | GSMASASSSQRGRSGSGNFGGGRGGGFGGNDNFGRGGNFSGRGGFGGSRG<br>GGGYGGSGDGYNGFGNDGSNFGGGGSYNDFGNYNNQSSNFGPMKGGNFGG<br>RSSGPYGGGGQYFAKPRNQGGYGGSSSSSSYGSGRRF | 1.25E-05 |
| WT | GSMASASSSQRGRSGSGNFGGGRGGGFGGNDNFGRGGNFSGRGGFGGSRG<br>GGGYGGSGDGYNGFGNDGSNFGGGGSYNDFGNYNNQSSNFGPMKGGNFGG<br>RSSGGSGGGGQYFAKPRNQGGYGGSSSSSSYGSGRRF | 1.25E-05 |
| -30G+30S-12F+12Y | GSMASASSSQRSRSSSGNYSGSRSGSYSGNDNYGRSGNYSGRSGYSGSRS<br>GSGYSGSSDSYNSYGNDSSNYSGSSSYNDYGNYNNQSSNYGPMKSGNYSG<br>RSSSSSGSSGQYYAKPRNQGSYSGSSSSSSYSSSRRY | 1.47E-05 |
| +2R | GSMASASSSQRGRSGSGNFGGGRGGGFGGNDNFGRGGNFSGRGGFGGSRG<br>GGGYGGSGDGYNGFRNDGSNFGGGRYNDFGNYNNQSSNFGPMKGGNFGG<br>RSSGPYGGGGQYFAKPRNQGGYGGSSSSSSYGSGRRF | 1.81E-05 |
| +8D | GSMASASSSQRDRSGSGNFGGGRDGGFGGNDNFGRGDNFSGRGDFGGSRD<br>GGGYGGSGDGYNGFGNDGSNFGGGGSYNDFGNYNNQSSNFGPMKGGNFGG<br>RSSDPYGGGGQYFAKPRNQDGYGGSSSSSSYDSGRRF | 1.84E-05 |
| -10G+10S | GSMASASSSQRSRSGSGNFGGGRSGGFGGNDNFGRSGNFSGRGGFGGSRG<br>GGGYGGSGDSYNGFGNDGSNFGGSGSYNDFGNYNNQSSNFGPMKSGNFGG<br>RSSGSSGGSGQYFAKPRNQGSYSGSSSSSSYGSGRRF | 2.76E-05 |
| -9F+6Y | GSMASASSSQRGRSGSGNFGGGRGGGYGGNDNYGRGGNYSGRGGFGGSRG<br>GGGYGGSGDGYNGGGNDGSNYGGGGSYNDSGNYNNQSSNFGPMKGGNYGG<br>RSSGGSGGGGQYGAKPRNQGGYGGSSSSSSYGSGRRY | 2.80E-05 |
| +7K+12D | GSMASADSSQRDRDDKGNFGDGRGGGFGGNDNFGRGGNFSDRGGFGGSRG<br>DGKYGGDGDKYNGFGNDGKNFGGGGSYNDFGNYNNQSSNFDPMKGGNFKD<br>RSSGPYDKGGQYFAKPRNQGGYGGSSSSKSYGSDRRF | 4.31E-05 |
| +7F-7Y | GSMASASSSQRGRSGSGNFGGGRGGGFGGNDNFGRGGNFSGRGGFGGSRG<br>GGGFGGSGDGFNGFGNDGSNFGGGGSFNDFGNFNNQSSNFGPMKGGNFGG<br>RSSGGSGGGGQFFAKPRNQGGFGGSSSSSSFGSGRRF | 4.94E-05 |
| -20G+20S | GSMASASSSQRSRSGSGNFSGSRSGSFSGNDNFGRSGNFSGRSGFGGSRS<br>GGGYSGSGDSYNSFGNDGSNFSGSGSYNDFGNYNNQSSNFGPMKSGNFGG<br>RSSGSSGGSGQYFAKPRNQGSYSGSSSSSSYGSSRRF | 5.39E-05 |
| -8F+4Y | GSMASASSSQRGRSGSGNFGGGRGGGYGGNDNGGRGGNYSGRGGFGGSRG<br>GGGYGGSGDGYNGGGNDGSNYGGGGSYNDSGNYNNQSSNFGPMKGGNYGG<br>RSSGGSGGGGQYGAKPRNQGGYGGSSSSSSYGSGRRF | 6.26E-05 |
| +23G-23S+7F-7Y | GSMAGAGGGQRGRGGGGNFGGGRGGGFGGNDNFGRGGNFGGRGGFGGGRG<br>GGGFGGGGDGFNGFGNDGGNFGGGGGFNDFGNFNNQGGNFGPMKGGNFGG<br>RGGGGGGGGGQFFAKPRNQGGFGGGGGGGGFGGGRRF | 7.63E-05 |

| | | |
|---|---|---|
| -3R+3K | GSMASASSSQRGKSGSGNFGGGRGGGFGGNDNFGRGGNFSGRGGFGGSKG GGGYGGSGDGYNGFGNDGSNFGGGGSYNDFGNYNNQSSNFGPMKGGNFGG RSSGGSGGGGQYFAKPRNQGGYGGSSSSSSYGSGRKF | 8.30E-05 |
| -4D | GSMASASSSQRGRSGSGNFGGGRGGGFGGNGNFGRGGNFSGRGGFGGSRG GGGYGGSGGGYNGFGNSGSNFGGGGSYNGFGNYNNQSSNFGPMKGGNFGG RSSGPYGGGGQYFAKPRNQGGYGGSSSSSSYGSGRRF | 8.69E-05 |
| -30G+30S+7F-7Y | GSMASASSSQRSRSSSGNFSGSRSGSFSGNDNFGRSGNFSGRSGFSGSRS GSGFSGSSDSFNSFGNDSSNFSGSSSFNDFGNFNNQSSNFGPMKSGNFSG RSSSSSGSSGQFFAKPRNQGSFSGSSSSSSFSSSRRF | 8.98E-05 |
| -20G+20S+7F-7Y | GSMASASSSQRSRSGSGNFSGSRSGSFSGNDNFGRSGNFSGRSGFGGSRS GGGFSGSGDSFNSFGNDGSNFSGSGSFNDFGNFNNQSSNFGPMKSGNFGG RSSGSSGGSGQFFAKPRNQGSFSGSSSSSSFGSSRRF | 9.87E-05 |
| +12D | GSMASADSSQRDRDDSGNFGDGRGGGFGGNDNFGRGGNFSDRGGFGGSRG DGGYGGDGDGYNGFGNDGSNFGGGGSYNDFGNYNNQSSNFDPMKGGNFGD RSSGPYDGGGQYFAKPRNQGGYGGSSSSSSYGSDRRF | 9.96E-05 |
| -4R-2K+5D | GSMASASSSQDGRSGSGNFGGGDGGGFGGNDNFGRGGNFSGGGGFGGSRG GGGYGGSGDGYNGFGNDGSNFGGGGSYNDFGNYNNQSSNFGPMDGGNFGG RSSGPYGGGGQYFADPRNQGGYGGSSSSSSYGSGDRF | 1.06E-04 |
| -9F+3Y | GSMASASSSQRGRSGSGNFGGGRGGGYGGNDNGGRGGNYSGRGGFGGSRG GGGYGGSGDGYNGGGNDGSNYGGGGSYNDSGNGNNQSSNFGPMKGGNYGG RSSGGSGGGGQYGAKPRNQGGYGGSSSSSSYGSGRRS | 1.13E-04 |
| -10R | GSMASASSSQGGSSGSGNFGGGGGGGFGGNDNFGGGGNFSGSGGFGGSGG GGGYGGSGDGYNGFGNDGSNFGGGGSYNDFGNYNNQSSNFGPMKGGNFGG SSSGPYGGGGQYFAKPGNQGGYGGSSSSSSYGSGGGF | 1.34E-04 |
| +7R | GSMASASSSQRGRSGRGNFGGGRGGGFGGNDNFGRGGNFSGRGGFGGSRG GGRYGGSGDRYNGFGNDGRNFGGGGSYNDFGNYNNQSSNFGPMKGGNFRG RSSGPYGRGGQYFAKPRNQGGYGGSSSSRSYGSGRRF | 1.78E-04 |
| +12E | GSMASAESSQREREESGNFGEGRGGGFGGNDNFGRGGNFSERGGFGGSRG EGGYGGEGDGYNGFGNDGSNFGGGGSYNDFGNYNNQSSNFEPMKGGNFGE RSSGPYEGGGQYFAKPRNQGGYGGSSSSSSYGSERRF | 2.12E-04 |
| Aro- | GSMASASSSQRGRSGSGNSGGGRGGGFGGNDNFGRGGNSSGRGGFGGSRG GGGYGGSGDGYNGFGNDGSNSGGGGSSNDFGNYNNQSSNFGPMKGGNFGG RSSGGSGGGGQYSAKPRNQGGYGGSSSSSSSGSGRRF | 3.01E-04 |
| -6R+6K | GSMASASSSQKGKSGSGNFGGGRGGGFGGNDNFGKGGNFSGRGGFGGSKG GGGYGGSGDGYNGFGNDGSNFGGGGSYNDFGNYNNQSSNFGPMKGGNFGG KSSGGSGGGGQYFAKPRNQGGYGGSSSSSSYGSGRKF | 4.96E-04 |

[a] Saturation concentration ($c_{sat}$) in molarity ($M$). Values were digitally extracted from Figures 1-5 and Supplementary Figures 2, 4, 6, in Bremer et al (13) and Figure 3F in Martin et al (12).

**Table S7. List of 500 proteins with the highest summed P classifier distance in the human proteome.**

| ∑ P class. dist. | longest PS IDR | first residue | last residue | UniProt ID and protein |
|---|---|---|---|---|
| 14249.16 | 2913 | 1714 | 4626 | Q7Z5P9\|MUC19_HUMAN Mucin-19 OS=Homo sapiens OX=9606 GN=MUC19 PE=1 |
| 10117.86 | 5705 | 995 | 6699 | A0A0G2JR97\|A0A0G2JR97_HUMAN Mucin-4 OS=Homo sapiens OX=9606 GN=MUC |
| 10109.08 | 5693 | 995 | 6687 | A0A0G2JS65\|A0A0G2JS65_HUMAN Mucin-4 OS=Homo sapiens OX=9606 GN=MUC |
| 10099.64 | 5441 | 995 | 6435 | A0A0G2JR46\|A0A0G2JR46_HUMAN Mucin-4 OS=Homo sapiens OX=9606 GN=MUC |
| 10092.09 | 5693 | 995 | 6687 | A0A0G2JQK9\|A0A0G2JQK9_HUMAN Mucin-4 OS=Homo sapiens OX=9606 GN=MUC |
| 10045.46 | 5453 | 995 | 6447 | A0A0G2JRD8\|A0A0G2JRD8_HUMAN Mucin-4 OS=Homo sapiens OX=9606 GN=MUC |
| 10040.92 | 5455 | 995 | 6449 | A0A0G2JRY3\|A0A0G2JRY3_HUMAN Mucin-4 OS=Homo sapiens OX=9606 GN=MUC |
| 10030.54 | 5469 | 995 | 6463 | A0A0G2JRJ6\|A0A0G2JRJ6_HUMAN Mucin-4 OS=Homo sapiens OX=9606 GN=MUC |
| 10021.77 | 5411 | 995 | 6405 | A0A0G2JRS2\|A0A0G2JRS2_HUMAN Mucin-4 OS=Homo sapiens OX=9606 GN=MUC |
| 10021.77 | 5521 | 995 | 6515 | A0A0G2JQI2\|A0A0G2JQI2_HUMAN Mucin-4 OS=Homo sapiens OX=9606 GN=MUC |
| 9926.97 | 2689 | 161 | 2849 | Q86YZ3\|HORN_HUMAN Hornerin OS=Homo sapiens OX=9606 GN=HRNR PE=1 SV |
| 7899.66 | 222 | 8676 | 8897 | Q8WXI7\|MUC16_HUMAN Mucin-16 OS=Homo sapiens OX=9606 GN=MUC16 PE=1 |
| 7632.47 | 5095 | 251 | 5345 | Q9UKN1\|MUC12_HUMAN Mucin-12 OS=Homo sapiens OX=9606 GN=MUC12 PE=1 |
| 6761.51 | 3574 | 963 | 4536 | A0A0G2JQT8\|A0A0G2JQT8_HUMAN Mucin-4 (Fragment) OS=Homo sapiens OX= |
| 6752.73 | 3571 | 963 | 4533 | A0A0G2JRA1\|A0A0G2JRA1_HUMAN Mucin-4 (Fragment) OS=Homo sapiens OX= |
| 6744.49 | 3441 | 963 | 4403 | A0A0G2JS91\|A0A0G2JS91_HUMAN Mucin-4 (Fragment) OS=Homo sapiens OX= |
| 6735.74 | 3571 | 963 | 4533 | A0A0G2JQC6\|A0A0G2JQC6_HUMAN Mucin-4 (Fragment) OS=Homo sapiens OX= |
| 6690.45 | 3453 | 963 | 4415 | A0A0G2JSB4\|A0A0G2JSB4_HUMAN Mucin-4 (Fragment) OS=Homo sapiens OX= |
| 6685.78 | 3455 | 963 | 4417 | A0A0G2JSD9\|A0A0G2JSD9_HUMAN Mucin-4 (Fragment) OS=Homo sapiens OX= |
| 6676.71 | 2716 | 1884 | 4599 | Q02817\|MUC2_HUMAN Mucin-2 OS=Homo sapiens OX=9606 GN=MUC2 PE=1 SV= |
| 6675.41 | 3469 | 963 | 4431 | A0A0G2JS19\|A0A0G2JS19_HUMAN Mucin-4 (Fragment) OS=Homo sapiens OX= |
| 6666.63 | 3521 | 963 | 4483 | A0A0G2JR43\|A0A0G2JR43_HUMAN Mucin-4 (Fragment) OS=Homo sapiens OX= |
| 6666.63 | 3411 | 963 | 4373 | A0A0G2JRE6\|A0A0G2JRE6_HUMAN Mucin-4 (Fragment) OS=Homo sapiens OX= |
| 6658.20 | 3565 | 990 | 4554 | E7ENC5\|E7ENC5_HUMAN Mucin-4 OS=Homo sapiens OX=9606 GN=MUC4 PE=1 S |
| 6649.42 | 3562 | 990 | 4551 | E9PDY6\|E9PDY6_HUMAN Mucin-4 OS=Homo sapiens OX=9606 GN=MUC4 PE=1 S |
| 6649.42 | 3565 | 963 | 4527 | A0A0G2JMX1\|A0A0G2JMX1_HUMAN Mucin-4 (Fragment) OS=Homo sapiens OX= |
| 6642.42 | 3556 | 891 | 4446 | A0A0G2JM16\|A0A0G2JM16_HUMAN Mucin-4 OS=Homo sapiens OX=9606 GN=MUC |
| 6640.64 | 3562 | 963 | 4524 | A0A0G2JNM3\|A0A0G2JNM3_HUMAN Mucin-4 (Fragment) OS=Homo sapiens OX= |
| 6633.80 | 3440 | 990 | 4429 | E7EQG8\|E7EQG8_HUMAN Mucin-4 OS=Homo sapiens OX=9606 GN=MUC4 PE=1 S |
| 6632.43 | 3562 | 990 | 4551 | E7EWN1\|E7EWN1_HUMAN Mucin-4 OS=Homo sapiens OX=9606 GN=MUC4 PE=1 S |
| 6626.06 | 3562 | 964 | 4525 | A0A0G2JN54\|A0A0G2JN54_HUMAN Mucin-4 OS=Homo sapiens OX=9606 GN=MUC |

| | | | | |
|---|---|---|---|---|
| 6625.01 | 3430 | 963 | 4392 | A0A0G2JQA9\|A0A0G2JQA9_HUMAN Mucin-4 (Fragment) OS=Homo sapiens OX= |
| 6625.01 | 3440 | 963 | 4402 | A0A0G2JS42\|A0A0G2JS42_HUMAN Mucin-4 (Fragment) OS=Homo sapiens OX= |
| 6623.64 | 3562 | 963 | 4524 | A0A0G2JRT1\|A0A0G2JRT1_HUMAN Mucin-4 (Fragment) OS=Homo sapiens OX= |
| 6585.80 | 3452 | 990 | 4441 | E7ERK0\|E7ERK0_HUMAN Mucin-4 OS=Homo sapiens OX=9606 GN=MUC4 PE=1 S |
| 6581.26 | 3454 | 990 | 4443 | E7EUL9\|E7EUL9_HUMAN Mucin-4 OS=Homo sapiens OX=9606 GN=MUC4 PE=1 S |
| 6577.01 | 3452 | 963 | 4414 | A0A0G2JRW6\|A0A0G2JRW6_HUMAN Mucin-4 (Fragment) OS=Homo sapiens OX= |
| 6572.47 | 3454 | 963 | 4416 | A0A0G2JRV5\|A0A0G2JRV5_HUMAN Mucin-4 (Fragment) OS=Homo sapiens OX= |
| 6570.88 | 3468 | 990 | 4457 | E7EQT2\|E7EQT2_HUMAN Mucin-4 OS=Homo sapiens OX=9606 GN=MUC4 PE=1 S |
| 6562.10 | 3520 | 990 | 4509 | E7ETT5\|E7ETT5_HUMAN Mucin-4 OS=Homo sapiens OX=9606 GN=MUC4 PE=1 S |
| 6562.10 | 3410 | 990 | 4399 | E7EW47\|E7EW47_HUMAN Mucin-4 OS=Homo sapiens OX=9606 GN=MUC4 PE=1 S |
| 6562.10 | 3468 | 963 | 4430 | A0A0G2JQN9\|A0A0G2JQN9_HUMAN Mucin-4 (Fragment) OS=Homo sapiens OX= |
| 6553.32 | 3520 | 963 | 4482 | A0A0G2JRK4\|A0A0G2JRK4_HUMAN Mucin-4 (Fragment) OS=Homo sapiens OX= |
| 6553.32 | 3410 | 963 | 4372 | A0A0G2JRW3\|A0A0G2JRW3_HUMAN Mucin-4 (Fragment) OS=Homo sapiens OX= |
| 6366.10 | 1339 | 2223 | 3561 | P98088\|MUC5A_HUMAN Mucin-5AC OS=Homo sapiens OX=9606 GN=MUC5AC PE= |
| 6226.32 | 2259 | 132 | 2390 | Q5D862\|FILA2_HUMAN Filaggrin-2 OS=Homo sapiens OX=9606 GN=FLG2 PE= |
| 5454.25 | 3764 | 297 | 4060 | P20930\|FILA_HUMAN Filaggrin OS=Homo sapiens OX=9606 GN=FLG PE=1 SV |
| 4726.82 | 689 | 4233 | 4921 | Q9HC84\|MUC5B_HUMAN Mucin-5B OS=Homo sapiens OX=9606 GN=MUC5B PE=1 |
| 4219.06 | 477 | 2087 | 2563 | Q02505\|MUC3A_HUMAN Mucin-3A OS=Homo sapiens OX=9606 GN=MUC3A PE=1 |
| 4033.40 | 857 | 442 | 1298 | Q685J3\|MUC17_HUMAN Mucin-17 OS=Homo sapiens OX=9606 GN=MUC17 PE=1 |
| 4032.16 | 857 | 442 | 1298 | E7EPM4\|E7EPM4_HUMAN Mucin-17 OS=Homo sapiens OX=9606 GN=MUC17 PE=1 |
| 2867.01 | 1659 | 1254 | 2912 | Q02388\|CO7A1_HUMAN Collagen alpha-1(VII) chain OS=Homo sapiens OX= |
| 2854.13 | 1531 | 54 | 1584 | P02462\|CO4A1_HUMAN Collagen alpha-1(IV) chain OS=Homo sapiens OX=9 |
| 2815.86 | 1565 | 33 | 1597 | P29400\|CO4A5_HUMAN Collagen alpha-5(IV) chain OS=Homo sapiens OX=9 |
| 2778.12 | 30 | 4145 | 4174 | P08519\|APOA_HUMAN Apolipoprotein(a) OS=Homo sapiens OX=9606 GN=LPA |
| 2585.77 | 596 | 933 | 1528 | A8MXH5\|A8MXH5_HUMAN Collagen alpha-6(IV) chain OS=Homo sapiens OX= |
| 2562.66 | 596 | 917 | 1512 | Q14031\|CO4A6_HUMAN Collagen alpha-6(IV) chain OS=Homo sapiens OX=9 |
| 2532.07 | 582 | 916 | 1497 | F5H851\|F5H851_HUMAN Collagen alpha-6(IV) chain OS=Homo sapiens OX= |
| 2521.38 | 1215 | 337 | 1551 | P53420\|CO4A4_HUMAN Collagen alpha-4(IV) chain OS=Homo sapiens OX=9 |
| 2514.09 | 581 | 916 | 1496 | A0A087WZY5\|A0A087WZY5_HUMAN Collagen alpha-6(IV) chain OS=Homo sap |
| 2504.77 | 1256 | 302 | 1557 | P08572\|CO4A2_HUMAN Collagen alpha-2(IV) chain OS=Homo sapiens OX=9 |
| 2457.82 | 544 | 916 | 1459 | F5H3Q5\|F5H3Q5_HUMAN Collagen alpha-6(IV) chain OS=Homo sapiens OX= |
| 2405.02 | 1265 | 91 | 1355 | P02461\|CO3A1_HUMAN Collagen alpha-1(III) chain OS=Homo sapiens OX= |
| 2372.37 | 1056 | 492 | 1547 | Q01955\|CO4A3_HUMAN Collagen alpha-3(IV) chain OS=Homo sapiens OX=9 |
| 2262.45 | 244 | 435 | 678 | A0A1B0GU24\|A0A1B0GU24_HUMAN Trinucleotide repeat-containing gene 6 |
| 2231.96 | 325 | 391 | 715 | Q8NDV7\|TNR6A_HUMAN Trinucleotide repeat-containing gene 6A protein |

| | | | | |
|---|---|---|---|---|
| 2231.54 | 1302 | 92 | 1393 | P05997\|CO5A2_HUMAN Collagen alpha-2(V) chain OS=Homo sapiens OX=96 |
| 2194.23 | 712 | 355 | 1066 | Q8N7X1\|RMXL3_HUMAN RNA-binding motif protein, X-linked-like-3 OS=H |
| 2129.73 | 1220 | 28 | 1247 | P08123\|CO1A2_HUMAN Collagen alpha-2(I) chain OS=Homo sapiens OX=96 |
| 2126.77 | 1214 | 31 | 1244 | A0A087WTA8\|A0A087WTA8_HUMAN Collagen alpha-2(I) chain OS=Homo sapi |
| 2068.48 | 426 | 212 | 637 | Q9UPQ9\|TNR6B_HUMAN Trinucleotide repeat-containing gene 6B protein |
| 2064.13 | 243 | 1188 | 1430 | Q12816\|TROP_HUMAN Trophinin OS=Homo sapiens OX=9606 GN=TRO PE=1 SV |
| 2023.20 | 380 | 1309 | 1688 | A0A6Q8NVI4\|A0A6Q8NVI4_HUMAN AT-rich interactive domain-containing |
| 2009.55 | 244 | 225 | 468 | Q9HCJ0\|TNR6C_HUMAN Trinucleotide repeat-containing gene 6C protein |
| 1956.85 | 1232 | 106 | 1337 | P02452\|CO1A1_HUMAN Collagen alpha-1(I) chain OS=Homo sapiens OX=96 |
| 1956.80 | 1263 | 93 | 1355 | P02458\|CO2A1_HUMAN Collagen alpha-1(II) chain OS=Homo sapiens OX=9 |
| 1921.32 | 531 | 564 | 1094 | Q9UMD9\|COHA1_HUMAN Collagen alpha-1(XVII) chain OS=Homo sapiens OX |
| 1910.96 | 593 | 1011 | 1603 | Q07092\|COGA1_HUMAN Collagen alpha-1(XVI) chain OS=Homo sapiens OX= |
| 1909.43 | 380 | 1269 | 1648 | A0A3F2YNW7\|A0A3F2YNW7_HUMAN AT-rich interactive domain-containing |
| 1890.60 | 1125 | 490 | 1614 | A0A0G2JL35\|A0A0G2JL35_HUMAN COL11A2 OS=Homo sapiens OX=9606 GN=COL |
| 1890.43 | 1125 | 490 | 1614 | A0A140TA43\|A0A140TA43_HUMAN COL11A2 OS=Homo sapiens OX=9606 GN=COL |
| 1884.98 | 1125 | 490 | 1614 | P13942\|COBA2_HUMAN Collagen alpha-2(XI) chain OS=Homo sapiens OX=9 |
| 1884.14 | 1125 | 490 | 1614 | A0A0C4DFS1\|A0A0C4DFS1_HUMAN COL11A2 OS=Homo sapiens OX=9606 GN=COL |
| 1880.29 | 1125 | 377 | 1501 | A0A140T9I7\|A0A140T9I7_HUMAN Collagen alpha-2(XI) chain (Fragment) |
| 1880.29 | 1125 | 404 | 1528 | Q4VXY6\|Q4VXY6_HUMAN Collagen alpha-2(XI) chain OS=Homo sapiens OX= |
| 1880.12 | 1125 | 404 | 1528 | A0A140T9N1\|A0A140T9N1_HUMAN Collagen alpha-2(XI) chain OS=Homo sap |
| 1879.97 | 1125 | 383 | 1507 | H0YIS1\|H0YIS1_HUMAN Collagen alpha-2(XI) chain OS=Homo sapiens OX= |
| 1879.80 | 1125 | 383 | 1507 | A0A140TA54\|A0A140TA54_HUMAN Collagen alpha-2(XI) chain OS=Homo sap |
| 1877.32 | 380 | 1173 | 1552 | Q8NFD5\|ARI1B_HUMAN AT-rich interactive domain-containing protein 1 |
| 1872.94 | 405 | 595 | 999 | O14497\|ARI1A_HUMAN AT-rich interactive domain-containing protein 1 |
| 1830.52 | 338 | 1752 | 2089 | P35658\|NU214_HUMAN Nuclear pore complex protein Nup214 OS=Homo sap |
| 1827.89 | 338 | 1740 | 2077 | A0A494C1F2\|A0A494C1F2_HUMAN Nuclear pore complex protein Nup214 OS |
| 1813.71 | 311 | 1 | 311 | P23490\|LORI_HUMAN Loricrin OS=Homo sapiens OX=9606 GN=LORICRIN PE= |
| 1812.15 | 1105 | 476 | 1580 | P25940\|CO5A3_HUMAN Collagen alpha-3(V) chain OS=Homo sapiens OX=96 |
| 1809.72 | 1103 | 562 | 1664 | P20908\|CO5A1_HUMAN Collagen alpha-1(V) chain OS=Homo sapiens OX=96 |
| 1798.87 | 1127 | 532 | 1658 | P12107\|COBA1_HUMAN Collagen alpha-1(XI) chain OS=Homo sapiens OX=9 |
| 1789.69 | 1143 | 483 | 1625 | Q8NFW1\|COMA1_HUMAN Collagen alpha-1(XXII) chain OS=Homo sapiens OX |
| 1777.68 | 554 | 1311 | 1864 | A0A0G2JN42\|A0A0G2JN42_HUMAN Mucin-6 OS=Homo sapiens OX=9606 GN=MUC |
| 1776.98 | 554 | 1311 | 1864 | Q6W4X9\|MUC6_HUMAN Mucin-6 OS=Homo sapiens OX=9606 GN=MUC6 PE=1 SV= |
| 1768.42 | 266 | 326 | 591 | Q92804\|RBP56_HUMAN TATA-binding protein-associated factor 2N OS=Ho |
| 1767.36 | 213 | 1650 | 1862 | A0A0G2JNJ8\|A0A0G2JNJ8_HUMAN Mucin-6 OS=Homo sapiens OX=9606 GN=MUC |

| | | | | |
|---|---|---|---|---|
| 1705.57 | 338 | 1181 | 1518 | A0A0A0MSW3\|A0A0A0MSW3_HUMAN Nuclear pore complex protein Nup214 OS |
| 1666.66 | 963 | 1 | 963 | A0A087WYX9\|A0A087WYX9_HUMAN Collagen alpha-2(V) chain OS=Homo sapi |
| 1665.08 | 1071 | 489 | 1559 | Q17RW2\|COOA1_HUMAN Collagen alpha-1(XXIV) chain OS=Homo sapiens OX |
| 1662.48 | 606 | 29 | 634 | E2RYF6\|MUC22_HUMAN Mucin-22 OS=Homo sapiens OX=9606 GN=MUC22 PE=1 |
| 1657.27 | 171 | 1 | 171 | P35527\|K1C9_HUMAN Keratin, type I cytoskeletal 9 OS=Homo sapiens O |
| 1657.11 | 375 | 1 | 375 | H0Y720\|H0Y720_HUMAN Trinucleotide repeat-containing gene 6B protei |
| 1593.26 | 314 | 1 | 314 | P35637\|FUS_HUMAN RNA-binding protein FUS OS=Homo sapiens OX=9606 G |
| 1588.95 | 313 | 1 | 313 | H3BPE7\|H3BPE7_HUMAN RNA-binding protein FUS OS=Homo sapiens OX=960 |
| 1571.71 | 338 | 578 | 915 | B7ZAV2\|B7ZAV2_HUMAN Nuclear pore complex protein Nup214 OS=Homo sa |
| 1566.62 | 163 | 1 | 163 | P13645\|K1C10_HUMAN Keratin, type I cytoskeletal 10 OS=Homo sapiens |
| 1517.96 | 385 | 512 | 896 | A0A0U1RQI7\|KLF18_HUMAN Kruppel-like factor 18 OS=Homo sapiens OX=9 |
| 1506.20 | 787 | 625 | 1411 | Q8IZC6\|CORA1_HUMAN Collagen alpha-1(XXVII) chain OS=Homo sapiens O |
| 1431.95 | 157 | 1 | 157 | P04264\|K2C1_HUMAN Keratin, type II cytoskeletal 1 OS=Homo sapiens |
| 1401.00 | 94 | 1455 | 1548 | Q9UPA5\|BSN_HUMAN Protein bassoon OS=Homo sapiens OX=9606 GN=BSN PE |
| 1390.15 | 290 | 605 | 894 | H0Y837\|H0Y837_HUMAN Nuclear pore complex protein Nup214 (Fragment) |
| 1374.30 | 130 | 2108 | 2237 | Q8NEZ4\|KMT2C_HUMAN Histone-lysine N-methyltransferase 2C OS=Homo s |
| 1352.52 | 291 | 1184 | 1474 | P49790\|NU153_HUMAN Nuclear pore complex protein Nup153 OS=Homo sap |
| 1347.59 | 274 | 467 | 740 | A0A0G2JNL3\|A0A0G2JNL3_HUMAN Mucin-4 OS=Homo sapiens OX=9606 GN=MUC |
| 1343.38 | 581 | 27 | 607 | A0A140T8X8\|A0A140T8X8_HUMAN Mucin-21 OS=Homo sapiens OX=9606 GN=MU |
| 1342.89 | 405 | 214 | 618 | A0A1B0GTU5\|A0A1B0GTU5_HUMAN AT-rich interactive domain-containing |
| 1342.05 | 274 | 467 | 740 | A0A0G2JPA4\|A0A0G2JPA4_HUMAN Mucin-4 OS=Homo sapiens OX=9606 GN=MUC |
| 1339.93 | 405 | 212 | 616 | H0Y488\|H0Y488_HUMAN AT-rich interactive domain-containing protein |
| 1339.80 | 207 | 137 | 343 | Q99102\|MUC4_HUMAN Mucin-4 OS=Homo sapiens OX=9606 GN=MUC4 PE=1 SV= |
| 1323.56 | 475 | 1495 | 1969 | P24928\|RPB1_HUMAN DNA-directed RNA polymerase II subunit RPB1 OS=H |
| 1317.29 | 791 | 510 | 1300 | Q9NZW4\|DSPP_HUMAN Dentin sialophosphoprotein OS=Homo sapiens OX=96 |
| 1298.72 | 193 | 1 | 193 | P52948\|NUP98_HUMAN Nuclear pore complex protein Nup98-Nup96 OS=Hom |
| 1293.42 | 119 | 2552 | 2670 | O14686\|KMT2D_HUMAN Histone-lysine N-methyltransferase 2D OS=Homo s |
| 1281.04 | 606 | 1397 | 2002 | A0A0G2JR65\|A0A0G2JR65_HUMAN Mucin-2 OS=Homo sapiens OX=9606 GN=MUC |
| 1279.85 | 626 | 69 | 694 | Q49AM6\|Q49AM6_HUMAN COL4A5 protein OS=Homo sapiens OX=9606 GN=COL4 |
| 1273.45 | 210 | 1512 | 1721 | Q5H9R4\|ARMX4_HUMAN Armadillo repeat-containing X-linked protein 4 |
| 1264.84 | 404 | 212 | 615 | A0A087WUV6\|A0A087WUV6_HUMAN AT-rich interactive domain-containing |
| 1264.40 | 627 | 417 | 1043 | Q14993\|COJA1_HUMAN Collagen alpha-1(XIX) chain OS=Homo sapiens OX= |
| 1263.12 | 755 | 728 | 1482 | P39060\|COIA1_HUMAN Collagen alpha-1(XVIII) chain OS=Homo sapiens O |
| 1259.34 | 547 | 27 | 573 | A0A0G2JKD1\|A0A0G2JKD1_HUMAN Mucin-21 OS=Homo sapiens OX=9606 GN=MU |
| 1243.81 | 113 | 1 | 113 | Q03164\|KMT2A_HUMAN Histone-lysine N-methyltransferase 2A OS=Homo s |

| | | | | |
|---|---|---|---|---|
| 1238.63 | 193 | 1 | 193 | A0A3B3ITD8\|A0A3B3ITD8_HUMAN Nuclear pore complex protein Nup98-Nup |
| 1238.18 | 273 | 39 | 311 | Q15517\|CDSN_HUMAN Corneodesmosin OS=Homo sapiens OX=9606 GN=CDSN P |
| 1231.60 | 157 | 1 | 157 | P35908\|K22E_HUMAN Keratin, type II cytoskeletal 2 epidermal OS=Hom |
| 1231.48 | 477 | 26 | 502 | A0A182DWF7\|A0A182DWF7_HUMAN Mucin-3A (Fragment) OS=Homo sapiens OX |
| 1225.83 | 273 | 39 | 311 | G8JLG2\|G8JLG2_HUMAN Corneodesmosin OS=Homo sapiens OX=9606 GN=CDSN |
| 1223.35 | 380 | 695 | 1074 | H0Y7H8\|H0Y7H8_HUMAN AT-rich interactive domain-containing protein |
| 1223.15 | 273 | 39 | 311 | Q2L6G8\|Q2L6G8_HUMAN Corneodesmosin OS=Homo sapiens OX=9606 GN=CDSN |
| 1209.74 | 114 | 642 | 755 | Q9UGU0\|TCF20_HUMAN Transcription factor 20 OS=Homo sapiens OX=9606 |
| 1199.54 | 82 | 2246 | 2327 | Q96JG9\|ZN469_HUMAN Zinc finger protein 469 OS=Homo sapiens OX=9606 |
| 1192.65 | 82 | 2274 | 2355 | H3BS19\|H3BS19_HUMAN Zinc finger protein 469 OS=Homo sapiens OX=960 |
| 1185.46 | 485 | 27 | 511 | A0A0G2JJF7\|A0A0G2JJF7_HUMAN Mucin-21 OS=Homo sapiens OX=9606 GN=MU |
| 1180.20 | 485 | 27 | 511 | Q5SSG8\|MUC21_HUMAN Mucin-21 OS=Homo sapiens OX=9606 GN=MUC21 PE=1 |
| 1178.88 | 487 | 27 | 513 | A0A0G2JHX4\|A0A0G2JHX4_HUMAN Mucin-21 OS=Homo sapiens OX=9606 GN=MU |
| 1177.93 | 109 | 597 | 705 | A0A0A0MTL4\|A0A0A0MTL4_HUMAN Neuron navigator 2 OS=Homo sapiens OX= |
| 1164.24 | 487 | 27 | 513 | A0A140TA38\|A0A140TA38_HUMAN Mucin-21 OS=Homo sapiens OX=9606 GN=MU |
| 1160.96 | 109 | 620 | 728 | Q8IVL1\|NAV2_HUMAN Neuron navigator 2 OS=Homo sapiens OX=9606 GN=NA |
| 1160.96 | 109 | 620 | 728 | A0A0A0MTE8\|A0A0A0MTE8_HUMAN Neuron navigator 2 OS=Homo sapiens OX= |
| 1141.13 | 374 | 1 | 374 | Q01844\|EWS_HUMAN RNA-binding protein EWS OS=Homo sapiens OX=9606 G |
| 1132.79 | 157 | 863 | 1019 | Q10571\|MN1_HUMAN Transcriptional activator MN1 OS=Homo sapiens OX= |
| 1131.54 | 61 | 388 | 448 | A0A0J9YXN7\|A0A0J9YXN7_HUMAN Perilipin-4 OS=Homo sapiens OX=9606 GN |
| 1129.72 | 337 | 100 | 436 | A0A494C0Y1\|A0A494C0Y1_HUMAN Nuclear pore complex protein Nup214 (F |
| 1127.65 | 61 | 373 | 433 | Q96Q06\|PLIN4_HUMAN Perilipin-4 OS=Homo sapiens OX=9606 GN=PLIN4 PE |
| 1102.34 | 84 | 1157 | 1240 | Q9Y566\|SHAN1_HUMAN SH3 and multiple ankyrin repeat domains protein |
| 1102.34 | 84 | 1165 | 1248 | H9KV90\|H9KV90_HUMAN SH3 and multiple ankyrin repeat domains protei |
| 1099.42 | 109 | 519 | 627 | P12035\|K2C3_HUMAN Keratin, type II cytoskeletal 3 OS=Homo sapiens |
| 1098.30 | 114 | 642 | 755 | A0A6Q8PH68\|A0A6Q8PH68_HUMAN Transcription factor 20 (Fragment) OS= |
| 1097.91 | 70 | 1571 | 1640 | Q68DE3\|USF3_HUMAN Basic helix-loop-helix domain-containing protein |
| 1081.47 | 352 | 1 | 352 | B0QYK0\|B0QYK0_HUMAN RNA-binding protein EWS OS=Homo sapiens OX=960 |
| 1072.95 | 1034 | 93 | 1126 | A0A087WWM1\|A0A087WWM1_HUMAN Mucin-1 OS=Homo sapiens OX=9606 GN=MUC |
| 1067.22 | 262 | 1 | 262 | H3BNZ4\|H3BNZ4_HUMAN RNA-binding protein FUS OS=Homo sapiens OX=960 |
| 1060.10 | 64 | 3492 | 3555 | A2VEC9\|SSPO_HUMAN SCO-spondin OS=Homo sapiens OX=9606 GN=SSPOP PE= |
| 1058.36 | 341 | 1495 | 1835 | A0A6Q8PGB0\|A0A6Q8PGB0_HUMAN DNA-directed RNA polymerase subunit OS |
| 1056.51 | 140 | 1073 | 1212 | Q15648\|MED1_HUMAN Mediator of RNA polymerase II transcription subu |
| 1047.86 | 1039 | 93 | 1131 | P15941\|MUC1_HUMAN Mucin-1 OS=Homo sapiens OX=9606 GN=MUC1 PE=1 SV= |
| 1044.87 | 130 | 366 | 495 | Q9Y6Q9\|NCOA3_HUMAN Nuclear receptor coactivator 3 OS=Homo sapiens |

| | | | | |
|---|---|---|---|---|
| 1041.03 | 120 | 518 | 637 | Q01546\|K22O_HUMAN Keratin, type II cytoskeletal 2 oral OS=Homo sap |
| 1040.79 | 662 | 27 | 688 | Q14055\|CO9A2_HUMAN Collagen alpha-2(IX) chain OS=Homo sapiens OX=9 |
| 1040.11 | 304 | 462 | 765 | P20849\|CO9A1_HUMAN Collagen alpha-1(IX) chain OS=Homo sapiens OX=9 |
| 1034.45 | 115 | 2011 | 2125 | O75179\|ANR17_HUMAN Ankyrin repeat domain-containing protein 17 OS= |
| 1034.23 | 242 | 133 | 374 | Q6E0U4\|DMKN_HUMAN Dermokine OS=Homo sapiens OX=9606 GN=DMKN PE=1 S |
| 1030.70 | 652 | 32 | 683 | Q14050\|CO9A3_HUMAN Collagen alpha-3(IX) chain OS=Homo sapiens OX=9 |
| 1024.56 | 284 | 529 | 812 | Q14157\|UBP2L_HUMAN Ubiquitin-associated protein 2-like OS=Homo sap |
| 1016.98 | 153 | 189 | 341 | Q92793\|CBP_HUMAN CREB-binding protein OS=Homo sapiens OX=9606 GN=C |
| 1014.49 | 67 | 4936 | 5002 | Q9Y6V0\|PCLO_HUMAN Protein piccolo OS=Homo sapiens OX=9606 GN=PCLO |
| 1012.35 | 244 | 426 | 669 | Q14686\|NCOA6_HUMAN Nuclear receptor coactivator 6 OS=Homo sapiens |
| 1002.13 | 64 | 4088 | 4151 | Q2LD37\|K1109_HUMAN Transmembrane protein KIAA1109 OS=Homo sapiens |
| 999.38 | 174 | 831 | 1004 | Q86UU0\|BCL9L_HUMAN B-cell CLL/lymphoma 9-like protein OS=Homo sapi |
| 999.37 | 190 | 1039 | 1228 | O00512\|BCL9_HUMAN B-cell CLL/lymphoma 9 protein OS=Homo sapiens OX |
| 996.58 | 174 | 794 | 967 | A0A087WZX0\|A0A087WZX0_HUMAN B-cell CLL/lymphoma 9-like protein OS= |
| 993.55 | 70 | 1034 | 1103 | Q8IVL0\|NAV3_HUMAN Neuron navigator 3 OS=Homo sapiens OX=9606 GN=NA |
| 986.36 | 284 | 540 | 823 | F8W726\|F8W726_HUMAN Ubiquitin-associated protein 2-like OS=Homo sa |
| 985.36 | 278 | 616 | 893 | Q12906\|ILF3_HUMAN Interleukin enhancer-binding factor 3 OS=Homo sa |
| 976.33 | 115 | 1048 | 1162 | Q9UQ35\|SRRM2_HUMAN Serine/arginine repetitive matrix protein 2 OS= |
| 975.22 | 319 | 1 | 319 | C9JGE3\|C9JGE3_HUMAN EWS RNA-binding protein variant 6 OS=Homo sapi |
| 973.33 | 184 | 1038 | 1221 | A8CG34\|P121C_HUMAN Nuclear envelope pore membrane protein POM 121C |
| 971.05 | 182 | 2733 | 2914 | Q99715\|COCA1_HUMAN Collagen alpha-1(XII) chain OS=Homo sapiens OX= |
| 958.46 | 177 | 2733 | 2909 | D6RGG3\|D6RGG3_HUMAN Collagen alpha-1(XII) chain OS=Homo sapiens OX |
| 958.16 | 459 | 245 | 703 | Q6XPR3\|RPTN_HUMAN Repetin OS=Homo sapiens OX=9606 GN=RPTN PE=1 SV= |
| 952.77 | 54 | 2383 | 2436 | Q9P2P6\|STAR9_HUMAN StAR-related lipid transfer protein 9 OS=Homo s |
| 951.29 | 102 | 656 | 757 | P35568\|IRS1_HUMAN Insulin receptor substrate 1 OS=Homo sapiens OX= |
| 946.89 | 106 | 883 | 988 | A0A2R8Y4T1\|A0A2R8Y4T1_HUMAN Tensin-1 OS=Homo sapiens OX=9606 GN=TN |
| 946.05 | 200 | 607 | 806 | Q5T6F2\|UBAP2_HUMAN Ubiquitin-associated protein 2 OS=Homo sapiens |
| 945.89 | 106 | 837 | 942 | A0A494C067\|A0A494C067_HUMAN Tensin-1 (Fragment) OS=Homo sapiens OX |
| 932.70 | 225 | 77 | 301 | Q09472\|EP300_HUMAN Histone acetyltransferase p300 OS=Homo sapiens |
| 931.41 | 72 | 1629 | 1700 | Q15911\|ZFHX3_HUMAN Zinc finger homeobox protein 3 OS=Homo sapiens |
| 930.65 | 317 | 623 | 939 | Q96QC0\|PP1RA_HUMAN Serine/threonine-protein phosphatase 1 regulato |
| 930.13 | 380 | 476 | 855 | A0A1B0GVK1\|A0A1B0GVK1_HUMAN AT-rich interactive domain-containing |
| 930.02 | 130 | 1815 | 1944 | Q2M2H8\|MGAL_HUMAN Probable maltase-glucoamylase 2 OS=Homo sapiens |
| 929.68 | 293 | 1 | 293 | A0A0D9SFL3\|A0A0D9SFL3_HUMAN RNA-binding protein EWS OS=Homo sapien |
| 929.14 | 238 | 20 | 257 | Q17RH7\|TPRXL_HUMAN Putative protein TPRXL OS=Homo sapiens OX=9606 |

| | | | | |
|---|---|---|---|---|
| 923.64 | 54 | 1238 | 1291 | P98160\|PGBM_HUMAN Basement membrane-specific heparan sulfate prote |
| 918.36 | 608 | 1 | 608 | A0A0G2JNG3\|A0A0G2JNG3_HUMAN Mucin-4 OS=Homo sapiens OX=9606 GN=MUC |
| 918.13 | 580 | 245 | 824 | Q2UY09\|COSA1_HUMAN Collagen alpha-1(XXVIII) chain OS=Homo sapiens |
| 917.35 | 569 | 35 | 603 | A0A0G2JLU8\|A0A0G2JLU8_HUMAN Mucin-4 OS=Homo sapiens OX=9606 GN=MUC |
| 911.60 | 225 | 77 | 301 | A0A669KB12\|A0A669KB12_HUMAN Histone acetyltransferase OS=Homo sapi |
| 909.19 | 158 | 232 | 389 | K7EQQ3\|K7EQQ3_HUMAN Keratin, type I cytoskeletal 9 OS=Homo sapiens |
| 906.71 | 33 | 2560 | 2592 | O60494\|CUBN_HUMAN Cubilin OS=Homo sapiens OX=9606 GN=CUBN PE=1 SV= |
| 905.15 | 106 | 737 | 842 | E9PGF5\|E9PGF5_HUMAN Tensin-1 OS=Homo sapiens OX=9606 GN=TNS1 PE=1 |
| 904.25 | 62 | 1430 | 1491 | I3L2J0\|I3L2J0_HUMAN Protein capicua homolog OS=Homo sapiens OX=960 |
| 904.15 | 106 | 737 | 842 | Q9HBL0\|TENS1_HUMAN Tensin-1 OS=Homo sapiens OX=9606 GN=TNS1 PE=1 S |
| 904.14 | 106 | 737 | 842 | E9PF55\|E9PF55_HUMAN Tensin-1 OS=Homo sapiens OX=9606 GN=TNS1 PE=1 |
| 903.24 | 55 | 194 | 248 | P48634\|PRC2A_HUMAN Protein PRRC2A OS=Homo sapiens OX=9606 GN=PRRC2 |
| 900.93 | 63 | 5827 | 5889 | Q09666\|AHNK_HUMAN Neuroblast differentiation-associated protein AH |
| 900.11 | 106 | 388 | 493 | A0A087WWW7\|A0A087WWW7_HUMAN Tensin-1 OS=Homo sapiens OX=9606 GN=TN |
| 896.87 | 287 | 195 | 481 | A0A6I8PTU7\|A0A6I8PTU7_HUMAN AT-rich interactive domain-containing |
| 895.36 | 132 | 653 | 784 | Q9H4A3\|WNK1_HUMAN Serine/threonine-protein kinase WNK1 OS=Homo sap |
| 892.75 | 119 | 2323 | 2441 | P25054\|APC_HUMAN Adenomatous polyposis coli protein OS=Homo sapien |
| 887.57 | 115 | 1895 | 2009 | H0YM23\|H0YM23_HUMAN Ankyrin repeat domain-containing protein 17 (F |
| 885.26 | 178 | 36 | 213 | A0A075B7F4\|A0A075B7F4_HUMAN TATA-binding protein-associated factor |
| 880.06 | 129 | 432 | 560 | Q9Y4H2\|IRS2_HUMAN Insulin receptor substrate 2 OS=Homo sapiens OX= |
| 878.19 | 182 | 1544 | 1725 | A0A087X0A8\|A0A087X0A8_HUMAN Collagen alpha-1(XII) chain OS=Homo sa |
| 875.82 | 157 | 1059 | 1215 | Q96HA1\|P121A_HUMAN Nuclear envelope pore membrane protein POM 121 |
| 871.82 | 296 | 650 | 945 | A6NCT7\|A6NCT7_HUMAN Collagen alpha-1(XVI) chain OS=Homo sapiens OX |
| 869.26 | 210 | 422 | 631 | Q9ULL5\|PRR12_HUMAN Proline-rich protein 12 OS=Homo sapiens OX=9606 |
| 868.87 | 31 | 34 | 64 | A0A3B3ISX9\|A0A3B3ISX9_HUMAN Tenascin-X OS=Homo sapiens OX=9606 GN= |
| 865.35 | 539 | 55 | 593 | Q03692\|COAA1_HUMAN Collagen alpha-1(X) chain OS=Homo sapiens OX=96 |
| 861.45 | 61 | 2132 | 2192 | Q5JSZ5\|PRC2B_HUMAN Protein PRRC2B OS=Homo sapiens OX=9606 GN=PRRC2 |
| 861.04 | 181 | 191 | 371 | P09651\|ROA1_HUMAN Heterogeneous nuclear ribonucleoprotein A1 OS=Ho |
| 855.43 | 521 | 80 | 600 | P25067\|CO8A2_HUMAN Collagen alpha-2(VIII) chain OS=Homo sapiens OX |
| 854.14 | 92 | 910 | 1001 | Q5VT52\|RPRD2_HUMAN Regulation of nuclear pre-mRNA domain-containin |
| 850.48 | 516 | 578 | 1093 | Q9C0J8\|WDR33_HUMAN pre-mRNA 3' end processing protein WDR33 OS=Hom |
| 848.95 | 382 | 1397 | 1778 | A0A0G2JM87\|A0A0G2JM87_HUMAN Mucin-2 OS=Homo sapiens OX=9606 GN=MUC |
| 841.89 | 145 | 653 | 797 | F5GWT4\|F5GWT4_HUMAN Non-specific serine/threonine protein kinase O |
| 839.73 | 35 | 917 | 951 | A0A140T902\|A0A140T902_HUMAN Tenascin-X OS=Homo sapiens OX=9606 GN= |
| 838.59 | 35 | 917 | 951 | A0A140T9C0\|A0A140T9C0_HUMAN Tenascin-X OS=Homo sapiens OX=9606 GN= |

| | | | | |
|---|---|---|---|---|
| 836.87 | 521 | 15 | 535 | E9PP49\|E9PP49_HUMAN Collagen alpha-2(VIII) chain OS=Homo sapiens O |
| 833.67 | 380 | 432 | 811 | A0A1B0GTJ8\|A0A1B0GTJ8_HUMAN AT-rich interactive domain-containing |
| 833.45 | 37 | 2223 | 2259 | Q9Y6R7\|FCGBP_HUMAN IgGFc-binding protein OS=Homo sapiens OX=9606 G |
| 833.44 | 98 | 492 | 589 | P13647\|K2C5_HUMAN Keratin, type II cytoskeletal 5 OS=Homo sapiens |
| 831.79 | 176 | 234 | 409 | Q99081\|HTF4_HUMAN Transcription factor 12 OS=Homo sapiens OX=9606 |
| 828.84 | 510 | 436 | 945 | Q96P44\|COLA1_HUMAN Collagen alpha-1(XXI) chain OS=Homo sapiens OX= |
| 827.75 | 35 | 917 | 951 | A0A140T8Y3\|A0A140T8Y3_HUMAN Tenascin-X OS=Homo sapiens OX=9606 GN= |
| 824.81 | 219 | 1010 | 1228 | A0A2R8Y651\|A0A2R8Y651_HUMAN PDZ domain-containing protein GIPC3 OS |
| 824.62 | 30 | 843 | 872 | Q8TCU4\|ALMS1_HUMAN Alstrom syndrome protein 1 OS=Homo sapiens OX=9 |
| 824.07 | 141 | 39 | 179 | A0A1B0GVR6\|A0A1B0GVR6_HUMAN Transcription factor 4 OS=Homo sapiens |
| 821.72 | 169 | 209 | 377 | P51991\|ROA3_HUMAN Heterogeneous nuclear ribonucleoprotein A3 OS=Ho |
| 821.72 | 101 | 475 | 575 | Q9BVL2\|NUP58_HUMAN Nucleoporin p58/p45 OS=Homo sapiens OX=9606 GN= |
| 821.66 | 212 | 387 | 598 | Q15596\|NCOA2_HUMAN Nuclear receptor coactivator 2 OS=Homo sapiens |
| 820.61 | 205 | 1143 | 1347 | Q15788\|NCOA1_HUMAN Nuclear receptor coactivator 1 OS=Homo sapiens |
| 820.52 | 35 | 917 | 951 | A0A140TA33\|A0A140TA33_HUMAN Tenascin-X OS=Homo sapiens OX=9606 GN= |
| 820.51 | 141 | 131 | 271 | E9PH57\|E9PH57_HUMAN Transcription factor 4 OS=Homo sapiens OX=9606 |
| 820.08 | 505 | 439 | 943 | F5GZK2\|F5GZK2_HUMAN Collagen alpha-1(XXI) chain OS=Homo sapiens OX |
| 819.38 | 35 | 917 | 951 | A0A140TA41\|A0A140TA41_HUMAN Tenascin-X OS=Homo sapiens OX=9606 GN= |
| 817.56 | 141 | 29 | 169 | P15884\|ITF2_HUMAN Transcription factor 4 OS=Homo sapiens OX=9606 G |
| 815.38 | 35 | 917 | 951 | P22105\|TENX_HUMAN Tenascin-X OS=Homo sapiens OX=9606 GN=TNXB PE=1 |
| 812.92 | 30 | 801 | 830 | A0A087WTU9\|A0A087WTU9_HUMAN Alstrom syndrome protein 1 OS=Homo sap |
| 812.30 | 141 | 29 | 169 | H3BTP3\|H3BTP3_HUMAN Transcription factor 4 OS=Homo sapiens OX=9606 |
| 809.01 | 41 | 3038 | 3078 | Q7Z407\|CSMD3_HUMAN CUB and sushi domain-containing protein 3 OS=Ho |
| 808.53 | 35 | 917 | 951 | A0A140TA52\|A0A140TA52_HUMAN Tenascin-X OS=Homo sapiens OX=9606 GN= |
| 807.15 | 110 | 416 | 525 | A0A2R8YDL9\|A0A2R8YDL9_HUMAN Methyl-CpG-binding domain protein 5 OS |
| 801.91 | 145 | 1 | 145 | H3BPJ7\|H3BPJ7_HUMAN Transcription factor 4 OS=Homo sapiens OX=9606 |
| 799.98 | 380 | 416 | 795 | A0A1B0GWJ2\|A0A1B0GWJ2_HUMAN AT-rich interactive domain-containing |
| 798.99 | 127 | 2560 | 2686 | O15417\|TNC18_HUMAN Trinucleotide repeat-containing gene 18 protein |
| 798.99 | 127 | 2560 | 2686 | H9KVB4\|H9KVB4_HUMAN Trinucleotide repeat-containing gene 18 protei |
| 791.12 | 237 | 126 | 362 | Q96F45\|ZN503_HUMAN Zinc finger protein 503 OS=Homo sapiens OX=9606 |
| 791.12 | 93 | 165 | 257 | O15027\|SC16A_HUMAN Protein transport protein Sec16A OS=Homo sapien |
| 789.84 | 149 | 579 | 727 | A0A2R8YGI3\|A0A2R8YGI3_HUMAN Collagen alpha-1(XIII) chain OS=Homo s |
| 788.40 | 323 | 27 | 349 | A0A0G2JMC4\|A0A0G2JMC4_HUMAN Mucin-21 OS=Homo sapiens OX=9606 GN=MU |
| 787.13 | 142 | 483 | 624 | O14654\|IRS4_HUMAN Insulin receptor substrate 4 OS=Homo sapiens OX= |
| 786.68 | 141 | 29 | 169 | A0A1B0GVB8\|A0A1B0GVB8_HUMAN Transcription factor 4 OS=Homo sapiens |

| | | | | |
|---|---|---|---|---|
| 784.58 | 216 | 465 | 680 | A0A6E1W314\|A0A6E1W314_HUMAN Collagen alpha-1(XIII) chain OS=Homo s |
| 784.23 | 205 | 992 | 1196 | B5MCN7\|B5MCN7_HUMAN Nuclear receptor coactivator 1 OS=Homo sapiens |
| 783.69 | 149 | 568 | 716 | Q5TAT6\|CODA1_HUMAN Collagen alpha-1(XIII) chain OS=Homo sapiens OX |
| 779.79 | 30 | 843 | 872 | A0A087WV20\|A0A087WV20_HUMAN Alstrom syndrome protein 1 OS=Homo sap |
| 779.07 | 144 | 61 | 204 | Q9UI36\|DACH1_HUMAN Dachshund homolog 1 OS=Homo sapiens OX=9606 GN= |
| 776.51 | 321 | 27 | 347 | A0A140TA51\|A0A140TA51_HUMAN Mucin-21 OS=Homo sapiens OX=9606 GN=MU |
| 775.34 | 161 | 643 | 803 | A6NF01\|P121B_HUMAN Putative nuclear envelope pore membrane protein |
| 774.85 | 124 | 713 | 836 | Q5SYE7\|NHSL1_HUMAN NHS-like protein 1 OS=Homo sapiens OX=9606 GN=N |
| 772.87 | 67 | 27 | 93 | Q8IWZ3\|ANKH1_HUMAN Ankyrin repeat and KH domain-containing protein |
| 771.21 | 47 | 2226 | 2272 | O15018\|PDZD2_HUMAN PDZ domain-containing protein 2 OS=Homo sapiens |
| 771.17 | 188 | 1273 | 1460 | E7EWN3\|E7EWN3_HUMAN Histone-lysine N-methyltransferase SETD5 OS=Ho |
| 770.13 | 216 | 383 | 598 | A0A669KB55\|A0A669KB55_HUMAN Collagen alpha-1(XIII) chain (Fragment |
| 768.26 | 107 | 907 | 1013 | Q8IZL2\|MAML2_HUMAN Mastermind-like protein 2 OS=Homo sapiens OX=96 |
| 768.25 | 516 | 120 | 635 | P27658\|CO8A1_HUMAN Collagen alpha-1(VIII) chain OS=Homo sapiens OX |
| 764.83 | 202 | 672 | 873 | Q92585\|MAML1_HUMAN Mastermind-like protein 1 OS=Homo sapiens OX=96 |
| 764.69 | 70 | 534 | 603 | A0A2R8YFX5\|A0A2R8YFX5_HUMAN Neuron navigator 3 OS=Homo sapiens OX= |
| 764.55 | 137 | 327 | 463 | P55197\|AF10_HUMAN Protein AF-10 OS=Homo sapiens OX=9606 GN=MLLT10 |
| 764.39 | 54 | 352 | 405 | Q9H195\|MUC3B_HUMAN Mucin-3B (Fragments) OS=Homo sapiens OX=9606 GN |
| 761.93 | 157 | 196 | 352 | P22626\|ROA2_HUMAN Heterogeneous nuclear ribonucleoproteins A2/B1 O |
| 761.72 | 91 | 46 | 136 | Q2KJY2\|KI26B_HUMAN Kinesin-like protein KIF26B OS=Homo sapiens OX= |
| 758.09 | 285 | 2119 | 2403 | P12111\|CO6A3_HUMAN Collagen alpha-3(VI) chain OS=Homo sapiens OX=9 |
| 757.78 | 156 | 716 | 871 | Q9NTZ6\|RBM12_HUMAN RNA-binding protein 12 OS=Homo sapiens OX=9606 |
| 755.58 | 126 | 75 | 200 | Q6L8H1\|KRA54_HUMAN Keratin-associated protein 5-4 OS=Homo sapiens |
| 752.27 | 93 | 165 | 257 | F1T0I1\|F1T0I1_HUMAN Protein transport protein sec16 OS=Homo sapien |
| 748.32 | 194 | 781 | 974 | O94913\|PCF11_HUMAN Pre-mRNA cleavage complex 2 protein Pcf11 OS=Ho |
| 744.20 | 149 | 1691 | 1839 | P16112\|PGCA_HUMAN Aggrecan core protein OS=Homo sapiens OX=9606 GN |
| 744.20 | 149 | 1691 | 1839 | H0YMF1\|H0YMF1_HUMAN Aggrecan core protein OS=Homo sapiens OX=9606 |
| 744.18 | 149 | 1672 | 1820 | A0A087X1T7\|A0A087X1T7_HUMAN Aggrecan core protein OS=Homo sapiens |
| 743.54 | 155 | 1 | 155 | Q2M2I5\|K1C24_HUMAN Keratin, type I cytoskeletal 24 OS=Homo sapiens |
| 738.90 | 188 | 1254 | 1441 | Q9C0A6\|SETD5_HUMAN Histone-lysine N-methyltransferase SETD5 OS=Hom |
| 738.12 | 33 | 325 | 357 | A0A087X0K4\|A0A087X0K4_HUMAN CUB and sushi domain-containing protei |
| 736.93 | 126 | 1718 | 1843 | Q8IZD2\|KMT2E_HUMAN Inactive histone-lysine N-methyltransferase 2E |
| 736.65 | 203 | 408 | 610 | A0A669KB28\|A0A669KB28_HUMAN Collagen alpha-1(XIII) chain (Fragment |
| 736.00 | 110 | 416 | 525 | Q9P267\|MBD5_HUMAN Methyl-CpG-binding domain protein 5 OS=Homo sapi |
| 736.00 | 110 | 416 | 525 | A0A1B0GW10\|A0A1B0GW10_HUMAN Methyl-CpG-binding domain protein 5 OS |

| | | | | |
|---|---|---|---|---|
| 734.22 | 167 | 229 | 395 | P15923\|TFE2_HUMAN Transcription factor E2-alpha OS=Homo sapiens OX |
| 732.11 | 73 | 88 | 160 | A0A2R8Y5P9\|A0A2R8Y5P9_HUMAN Protein Shroom3 OS=Homo sapiens OX=960 |
| 731.60 | 171 | 1455 | 1625 | Q05707\|COEA1_HUMAN Collagen alpha-1(XIV) chain OS=Homo sapiens OX= |
| 730.82 | 463 | 182 | 644 | A8MWQ5\|A8MWQ5_HUMAN Collagen alpha-1(XXV) chain OS=Homo sapiens OX |
| 730.29 | 111 | 1287 | 1397 | A0A6Q8PFM0\|A0A6Q8PFM0_HUMAN Serine/threonine-protein kinase WNK1 ( |
| 729.57 | 149 | 1215 | 1363 | A0A5K1VW97\|A0A5K1VW97_HUMAN Aggrecan core protein (Fragment) OS=Ho |
| 728.34 | 474 | 196 | 669 | A0A2R8Y760\|A0A2R8Y760_HUMAN Collagen alpha-1(XXV) chain OS=Homo sa |
| 727.41 | 174 | 230 | 403 | B4DGI9\|B4DGI9_HUMAN Transcription factor 12 (Fragment) OS=Homo sap |
| 727.02 | 107 | 175 | 281 | Q3L8U1\|CHD9_HUMAN Chromodomain-helicase-DNA-binding protein 9 OS=H |
| 725.68 | 115 | 815 | 929 | Q9H2D6\|TARA_HUMAN TRIO and F-actin-binding protein OS=Homo sapiens |
| 725.42 | 285 | 1512 | 1796 | E7ENL6\|E7ENL6_HUMAN Collagen alpha-3(VI) chain OS=Homo sapiens OX= |
| 725.23 | 95 | 360 | 454 | Q96JK9\|MAML3_HUMAN Mastermind-like protein 3 OS=Homo sapiens OX=96 |
| 725.18 | 62 | 733 | 794 | Q9C0C2\|TB182_HUMAN 182 kDa tankyrase-1-binding protein OS=Homo sap |
| 718.21 | 145 | 401 | 545 | Q9ULJ6\|ZMIZ1_HUMAN Zinc finger MIZ domain-containing protein 1 OS= |
| 717.59 | 94 | 484 | 577 | Q7Z794\|K2C1B_HUMAN Keratin, type II cytoskeletal 1b OS=Homo sapien |
| 717.14 | 76 | 1145 | 1220 | O43166\|SI1L1_HUMAN Signal-induced proliferation-associated 1-like |
| 715.72 | 125 | 365 | 489 | P54259\|ATN1_HUMAN Atrophin-1 OS=Homo sapiens OX=9606 GN=ATN1 PE=1 |
| 714.37 | 73 | 169 | 241 | Q8TF72\|SHRM3_HUMAN Protein Shroom3 OS=Homo sapiens OX=9606 GN=SHRO |
| 711.74 | 71 | 944 | 1014 | Q8IZF6\|AGRG4_HUMAN Adhesion G-protein coupled receptor G4 OS=Homo |
| 710.77 | 193 | 325 | 517 | Q8NCA5\|FA98A_HUMAN Protein FAM98A OS=Homo sapiens OX=9606 GN=FAM98 |
| 710.25 | 49 | 2698 | 2746 | Q15751\|HERC1_HUMAN Probable E3 ubiquitin-protein ligase HERC1 OS=H |
| 707.49 | 216 | 409 | 624 | A0A669KB16\|A0A669KB16_HUMAN Collagen alpha-1(XIII) chain OS=Homo s |
| 707.14 | 41 | 2384 | 2424 | Q7Z7M0\|MEGF8_HUMAN Multiple epidermal growth factor-like domains p |
| 706.93 | 87 | 14 | 100 | P08047\|SP1_HUMAN Transcription factor Sp1 OS=Homo sapiens OX=9606 |
| 702.59 | 87 | 408 | 494 | E9PNV5\|E9PNV5_HUMAN Neuron navigator 2 (Fragment) OS=Homo sapiens |
| 702.45 | 236 | 406 | 641 | E7ES50\|E7ES50_HUMAN Collagen alpha-1(XIII) chain OS=Homo sapiens O |
| 701.61 | 122 | 993 | 1114 | Q5TGY3\|AHDC1_HUMAN AT-hook DNA-binding motif-containing protein 1 |
| 701.31 | 137 | 819 | 955 | Q8IWN7\|RP1L1_HUMAN Retinitis pigmentosa 1-like 1 protein OS=Homo s |
| 700.85 | 126 | 638 | 763 | Q5T1Z8\|Q5T1Z8_HUMAN Pumilio homolog 1 OS=Homo sapiens OX=9606 GN=P |
| 699.46 | 167 | 258 | 424 | X6REB3\|X6REB3_HUMAN Transcription factor E2-alpha OS=Homo sapiens |
| 697.77 | 210 | 448 | 657 | A0A669KAZ4\|A0A669KAZ4_HUMAN Collagen alpha-1(XIII) chain OS=Homo s |
| 696.33 | 37 | 2779 | 2815 | O75592\|MYCB2_HUMAN E3 ubiquitin-protein ligase MYCBP2 OS=Homo sapi |
| 696.21 | 72 | 1290 | 1361 | O15021\|MAST4_HUMAN Microtubule-associated serine/threonine-protein |
| 693.40 | 413 | 1 | 413 | A0A3B3ITG7\|A0A3B3ITG7_HUMAN Collagen alpha-1(IV) chain (Fragment) |
| 691.43 | 233 | 291 | 523 | P0CG12\|DERPC_HUMAN Decreased expression in renal and prostate canc |

| | | | | |
|---|---|---|---|---|
| 689.68 | 93 | 325 | 417 | Q8NF64\|ZMIZ2_HUMAN Zinc finger MIZ domain-containing protein 2 OS= |
| 689.68 | 93 | 325 | 417 | A0A087X127\|A0A087X127_HUMAN Zinc finger MIZ domain-containing prot |
| 689.34 | 119 | 1540 | 1658 | Q71F56\|MD13L_HUMAN Mediator of RNA polymerase II transcription sub |
| 689.34 | 119 | 1540 | 1658 | A0A3B3IRX3\|A0A3B3IRX3_HUMAN Mediator of RNA polymerase II transcri |
| 685.05 | 33 | 325 | 357 | Q7Z408\|CSMD2_HUMAN CUB and sushi domain-containing protein 2 OS=Ho |
| 684.01 | 62 | 514 | 575 | A0A3B3IRW6\|A0A3B3IRW6_HUMAN Glutamine and serine-rich protein 1 OS |
| 683.54 | 263 | 100 | 362 | H0Y2R3\|H0Y2R3_HUMAN AT-rich interactive domain-containing protein |
| 683.38 | 401 | 489 | 889 | F8WDM8\|F8WDM8_HUMAN Collagen alpha-1(XXIV) chain OS=Homo sapiens O |
| 682.85 | 126 | 602 | 727 | Q14671\|PUM1_HUMAN Pumilio homolog 1 OS=Homo sapiens OX=9606 GN=PUM |
| 682.85 | 126 | 603 | 728 | Q5T1Z4\|Q5T1Z4_HUMAN Pumilio homolog 1 OS=Homo sapiens OX=9606 GN=P |
| 682.66 | 97 | 2039 | 2135 | Q96RV3\|PCX1_HUMAN Pecanex-like protein 1 OS=Homo sapiens OX=9606 G |
| 680.87 | 59 | 1206 | 1264 | P10071\|GLI3_HUMAN Transcriptional activator GLI3 OS=Homo sapiens O |
| 677.41 | 94 | 2461 | 2554 | Q12830\|BPTF_HUMAN Nucleosome-remodeling factor subunit BPTF OS=Hom |
| 677.13 | 65 | 1 | 65 | A0A088AWL3\|A0A088AWL3_HUMAN Nuclear receptor corepressor 1 OS=Homo |
| 676.89 | 458 | 196 | 653 | Q9BXS0\|COPA1_HUMAN Collagen alpha-1(XXV) chain OS=Homo sapiens OX= |
| 676.19 | 56 | 2338 | 2393 | O75376\|NCOR1_HUMAN Nuclear receptor corepressor 1 OS=Homo sapiens |
| 676.14 | 193 | 261 | 453 | P02671\|FIBA_HUMAN Fibrinogen alpha chain OS=Homo sapiens OX=9606 G |
| 674.88 | 37 | 2741 | 2777 | A0A499FJI4\|A0A499FJI4_HUMAN RCR-type E3 ubiquitin transferase OS=H |
| 671.51 | 34 | 270 | 303 | P02751\|FINC_HUMAN Fibronectin OS=Homo sapiens OX=9606 GN=FN1 PE=1 |
| 671.21 | 85 | 45 | 129 | O94916\|NFAT5_HUMAN Nuclear factor of activated T-cells 5 OS=Homo s |
| 670.98 | 63 | 65 | 127 | Q6L8H4\|KRA51_HUMAN Keratin-associated protein 5-1 OS=Homo sapiens |
| 670.25 | 140 | 334 | 473 | A0A6Q8PH46\|A0A6Q8PH46_HUMAN Mucin-19 (Fragment) OS=Homo sapiens OX |
| 670.05 | 107 | 737 | 843 | A0A087X0G5\|A0A087X0G5_HUMAN Mastermind-like protein 2 OS=Homo sapi |
| 667.88 | 84 | 2268 | 2351 | Q9Y520\|PRC2C_HUMAN Protein PRRC2C OS=Homo sapiens OX=9606 GN=PRRC2 |
| 667.59 | 77 | 1077 | 1153 | P49792\|RBP2_HUMAN E3 SUMO-protein ligase RanBP2 OS=Homo sapiens OX |
| 667.39 | 89 | 1984 | 2072 | Q8WYB5\|KAT6B_HUMAN Histone acetyltransferase KAT6B OS=Homo sapiens |
| 667.04 | 359 | 305 | 663 | A0A0U1RRA7\|A0A0U1RRA7_HUMAN Collagen alpha-1(XI) chain (Fragment) |
| 665.90 | 84 | 2270 | 2353 | E7EPN9\|E7EPN9_HUMAN Protein PRRC2C OS=Homo sapiens OX=9606 GN=PRRC |
| 665.83 | 59 | 1147 | 1205 | A0A2R8YGX0\|A0A2R8YGX0_HUMAN Transcriptional activator GLI3 OS=Homo |
| 663.48 | 36 | 2997 | 3032 | Q96JQ0\|PCD16_HUMAN Protocadherin-16 OS=Homo sapiens OX=9606 GN=DCH |
| 663.44 | 62 | 521 | 582 | A0A0A0MQR4\|A0A0A0MQR4_HUMAN Protein capicua homolog OS=Homo sapien |
| 661.18 | 76 | 809 | 884 | P10070\|GLI2_HUMAN Zinc finger protein GLI2 OS=Homo sapiens OX=9606 |
| 660.58 | 62 | 521 | 582 | Q96RK0\|CIC_HUMAN Protein capicua homolog OS=Homo sapiens OX=9606 G |
| 658.62 | 77 | 1 | 77 | A0A3F2YNZ0\|A0A3F2YNZ0_HUMAN Protein transport protein sec16 OS=Hom |
| 656.28 | 110 | 416 | 525 | A0A0D9SG23\|A0A0D9SG23_HUMAN Methyl-CpG-binding domain protein 5 OS |

| | | | | |
|---|---|---|---|---|
| 656.18 | 50 | 3262 | 3311 | Q9NYQ7\|CELR3_HUMAN Cadherin EGF LAG seven-pass G-type receptor 3 O |
| 656.10 | 77 | 1266 | 1342 | Q68CP9\|ARID2_HUMAN AT-rich interactive domain-containing protein 2 |
| 656.10 | 77 | 1240 | 1316 | F8WCU9\|F8WCU9_HUMAN AT-rich interactive domain-containing protein |
| 654.13 | 69 | 552 | 620 | Q86YV5\|PRAG1_HUMAN Inactive tyrosine-protein kinase PRAG1 OS=Homo |
| 650.98 | 84 | 240 | 323 | Q15714\|T22D1_HUMAN TSC22 domain family protein 1 OS=Homo sapiens O |
| 648.63 | 179 | 572 | 750 | A0A087X0K0\|A0A087X0K0_HUMAN Collagen alpha-1(XV) chain OS=Homo sap |
| 647.08 | 77 | 876 | 952 | F8W108\|F8W108_HUMAN AT-rich interactive domain-containing protein |
| 646.71 | 66 | 774 | 839 | P55198\|AF17_HUMAN Protein AF-17 OS=Homo sapiens OX=9606 GN=MLLT6 P |
| 646.50 | 77 | 1 | 77 | Q92945\|FUBP2_HUMAN Far upstream element-binding protein 2 OS=Homo |
| 645.31 | 36 | 4719 | 4754 | Q96RW7\|HMCN1_HUMAN Hemicentin-1 OS=Homo sapiens OX=9606 GN=HMCN1 P |
| 644.29 | 182 | 371 | 552 | H0Y5N9\|H0Y5N9_HUMAN Collagen alpha-1(XII) chain (Fragment) OS=Homo |
| 643.38 | 29 | 1393 | 1421 | E5RIG2\|E5RIG2_HUMAN CUB and sushi domain-containing protein 1 OS=H |
| 642.98 | 122 | 906 | 1027 | A0A669KBM4\|A0A669KBM4_HUMAN DNA-binding protein RFX7 OS=Homo sapie |
| 642.06 | 77 | 1 | 77 | A0A3F2YNX0\|A0A3F2YNX0_HUMAN Protein transport protein sec16 OS=Hom |
| 640.12 | 29 | 1392 | 1420 | Q96PZ7\|CSMD1_HUMAN CUB and sushi domain-containing protein 1 OS=Ho |
| 639.00 | 179 | 586 | 764 | P39059\|COFA1_HUMAN Collagen alpha-1(XV) chain OS=Homo sapiens OX=9 |
| 637.56 | 91 | 109 | 199 | P31942\|HNRH3_HUMAN Heterogeneous nuclear ribonucleoprotein H3 OS=H |
| 636.90 | 66 | 1 | 66 | Q6ZRS2\|SRCAP_HUMAN Helicase SRCAP OS=Homo sapiens OX=9606 GN=SRCAP |
| 635.17 | 29 | 1393 | 1421 | F8W9C3\|F8W9C3_HUMAN CUB and sushi domain-containing protein 1 OS=H |
| 635.14 | 244 | 426 | 669 | F6M2K2\|F6M2K2_HUMAN Nuclear receptor coactivator 6 OS=Homo sapiens |
| 635.13 | 41 | 728 | 768 | E7EVZ1\|E7EVZ1_HUMAN Zinc finger homeobox protein 4 OS=Homo sapiens |
| 631.47 | 171 | 416 | 586 | A0A0A0MQT7\|A0A0A0MQT7_HUMAN Collagen alpha-1(XIV) chain OS=Homo sa |
| 630.89 | 227 | 163 | 389 | Q96E39\|RMXL1_HUMAN RNA binding motif protein, X-linked-like-1 OS=H |
| 630.73 | 354 | 1403 | 1756 | A8TX70\|CO6A5_HUMAN Collagen alpha-5(VI) chain OS=Homo sapiens OX=9 |
| 630.73 | 354 | 1403 | 1756 | E9PAL5\|E9PAL5_HUMAN Collagen alpha-5(VI) chain OS=Homo sapiens OX= |
| 630.59 | 40 | 874 | 913 | C9JG08\|C9JG08_HUMAN Uncharacterized protein C2orf16 OS=Homo sapien |
| 630.14 | 41 | 728 | 768 | Q86UP3\|ZFHX4_HUMAN Zinc finger homeobox protein 4 OS=Homo sapiens |
| 628.75 | 103 | 1275 | 1377 | O14513\|NCKP5_HUMAN Nck-associated protein 5 OS=Homo sapiens OX=960 |
| 628.75 | 103 | 1275 | 1377 | A0A0A0MS79\|A0A0A0MS79_HUMAN Nck-associated protein 5 OS=Homo sapie |
| 628.64 | 63 | 382 | 444 | G3V5H7\|G3V5H7_HUMAN SKI family transcriptional corepressor 1 OS=Ho |
| 628.47 | 74 | 54 | 127 | H7C269\|H7C269_HUMAN Trinucleotide repeat-containing gene 6A protei |
| 625.32 | 63 | 410 | 472 | P84550\|SKOR1_HUMAN SKI family transcriptional corepressor 1 OS=Hom |
| 625.11 | 244 | 426 | 669 | F6M2K4\|F6M2K4_HUMAN Nuclear receptor coactivator 6 OS=Homo sapiens |
| 624.99 | 122 | 809 | 930 | Q2KHR2\|RFX7_HUMAN DNA-binding protein RFX7 OS=Homo sapiens OX=9606 |
| 624.64 | 114 | 844 | 957 | Q8NET4\|RTL9_HUMAN Retrotransposon Gag-like protein 9 OS=Homo sapie |

| | | | | |
|---|---|---|---|---|
| 624.48 | 87 | 875 | 961 | A0A2R8YDS2\|A0A2R8YDS2_HUMAN Ras/Rap GTPase-activating protein SynG |
| 623.57 | 462 | 32 | 493 | A0A087X1E1\|A0A087X1E1_HUMAN Collagen alpha-1(XXV) chain OS=Homo sa |
| 620.50 | 110 | 507 | 616 | Q6AI39\|BICRL_HUMAN BRD4-interacting chromatin-remodeling complex-a |
| 620.35 | 64 | 788 | 851 | Q9ULM3\|YETS2_HUMAN YEATS domain-containing protein 2 OS=Homo sapie |
| 620.16 | 77 | 1 | 77 | A0A087WTP3\|A0A087WTP3_HUMAN Far upstream element-binding protein 2 |
| 619.50 | 104 | 267 | 370 | Q96PE2\|ARHGH_HUMAN Rho guanine nucleotide exchange factor 17 OS=Ho |
| 618.95 | 35 | 917 | 951 | A0A140T956\|A0A140T956_HUMAN Tenascin-X (Fragment) OS=Homo sapiens |
| 618.51 | 379 | 294 | 672 | A0A1B0GV63\|A0A1B0GV63_HUMAN AT-rich interactive domain-containing |
| 616.85 | 71 | 713 | 783 | A0A669KBC5\|A0A669KBC5_HUMAN Protein unc-80 homolog OS=Homo sapiens |
| 616.16 | 90 | 194 | 283 | H0Y4U1\|H0Y4U1_HUMAN Tensin-1 OS=Homo sapiens OX=9606 GN=TNS1 PE=1 |
| 615.33 | 87 | 920 | 1006 | B7ZCA0\|B7ZCA0_HUMAN Ras/Rap GTPase-activating protein SynGAP OS=Ho |
| 614.92 | 37 | 2223 | 2259 | A0A087WXI2\|A0A087WXI2_HUMAN IgGFc-binding protein OS=Homo sapiens |
| 614.79 | 71 | 713 | 783 | Q8N2C7\|UNC80_HUMAN Protein unc-80 homolog OS=Homo sapiens OX=9606 |
| 614.79 | 71 | 713 | 783 | A0A669KAW8\|A0A669KAW8_HUMAN Protein unc-80 homolog OS=Homo sapiens |
| 614.14 | 167 | 178 | 344 | A0A0A0MRB7\|A0A0A0MRB7_HUMAN Transcription factor E2-alpha OS=Homo |
| 613.57 | 291 | 1 | 291 | F8WC90\|F8WC90_HUMAN RNA-binding protein EWS (Fragment) OS=Homo sap |
| 612.12 | 87 | 934 | 1020 | Q96PV0\|SYGP1_HUMAN Ras/Rap GTPase-activating protein SynGAP OS=Hom |
| 611.74 | 87 | 934 | 1020 | A0A2R8Y6T2\|A0A2R8Y6T2_HUMAN Ras/Rap GTPase-activating protein SynG |
| 611.53 | 83 | 179 | 261 | Q9P2D1\|CHD7_HUMAN Chromodomain-helicase-DNA-binding protein 7 OS=H |
| 611.28 | 87 | 919 | 1005 | A0A0A0MQZ2\|A0A0A0MQZ2_HUMAN Ras/Rap GTPase-activating protein SynG |
| 610.35 | 174 | 64 | 237 | F5GY10\|F5GY10_HUMAN Transcription factor 12 OS=Homo sapiens OX=960 |
| 610.21 | 63 | 371 | 433 | G3V3E1\|G3V3E1_HUMAN SKI family transcriptional corepressor 1 OS=Ho |
| 610.13 | 94 | 203 | 296 | Q5JU85\|IQEC2_HUMAN IQ motif and SEC7 domain-containing protein 2 O |
| 608.34 | 61 | 1364 | 1424 | A0A494C0D3\|A0A494C0D3_HUMAN Protein PRRC2B (Fragment) OS=Homo sapi |
| 608.25 | 122 | 119 | 240 | E7EX21\|E7EX21_HUMAN Collagen alpha-1(XIII) chain OS=Homo sapiens O |
| 607.09 | 37 | 331 | 367 | Q9UGM3\|DMBT1_HUMAN Deleted in malignant brain tumors 1 protein OS= |
| 606.34 | 60 | 1153 | 1212 | A0A1U9X989\|A0A1U9X989_HUMAN NOTCH4 OS=Homo sapiens OX=9606 GN=NOTC |
| 604.97 | 60 | 1154 | 1213 | Q99466\|NOTC4_HUMAN Neurogenic locus notch homolog protein 4 OS=Hom |
| 604.29 | 59 | 2398 | 2456 | F5GXF5\|F5GXF5_HUMAN Nucleosome-remodeling factor subunit BPTF (Fra |
| 604.18 | 100 | 7935 | 8034 | A6NGQ3\|A6NGQ3_HUMAN Non-specific serine/threonine protein kinase O |
| 603.98 | 29 | 1254 | 1282 | F5GZ18\|F5GZ18_HUMAN CUB and sushi domain-containing protein 1 OS=H |
| 603.98 | 114 | 966 | 1079 | Q9NZP6\|NPAP1_HUMAN Nuclear pore-associated protein 1 OS=Homo sapie |
| 603.41 | 60 | 1156 | 1215 | A0A140T9R5\|A0A140T9R5_HUMAN NOTCH4 OS=Homo sapiens OX=9606 GN=NOTC |
| 603.14 | 37 | 331 | 367 | A0A590UJ76\|A0A590UJ76_HUMAN Deleted in malignant brain tumors 1 pr |
| 603.11 | 50 | 104 | 153 | P15502\|ELN_HUMAN Elastin OS=Homo sapiens OX=9606 GN=ELN PE=1 SV=4 |

| | | | | |
|---|---|---|---|---|
| 602.98 | 228 | 163 | 390 | P38159\|RBMX_HUMAN RNA-binding motif protein, X chromosome OS=Homo |
| 600.93 | 122 | 809 | 930 | H0YLX2\|H0YLX2_HUMAN DNA-binding protein RFX7 OS=Homo sapiens OX=96 |
| 600.77 | 93 | 293 | 385 | E7EWM3\|E7EWM3_HUMAN Zinc finger MIZ domain-containing protein 2 OS |
| 600.11 | 133 | 1 | 133 | A0A2R8YET7\|A0A2R8YET7_HUMAN Eyes absent homolog OS=Homo sapiens OX |
| 599.91 | 41 | 2308 | 2348 | H7BXX0\|H7BXX0_HUMAN CUB and sushi domain-containing protein 3 (Fra |
| 599.66 | 65 | 499 | 563 | P04259\|K2C6B_HUMAN Keratin, type II cytoskeletal 6B OS=Homo sapien |
| 597.89 | 65 | 499 | 563 | P02538\|K2C6A_HUMAN Keratin, type II cytoskeletal 6A OS=Homo sapien |
| 597.69 | 96 | 1291 | 1386 | A6NEM2\|A6NEM2_HUMAN Host cell factor 1 OS=Homo sapiens OX=9606 GN= |
| 597.42 | 66 | 126 | 191 | Q9ULD9\|ZN608_HUMAN Zinc finger protein 608 OS=Homo sapiens OX=9606 |
| 597.34 | 84 | 203 | 286 | A0A6Q8PFR7\|A0A6Q8PFR7_HUMAN IQ motif and SEC7 domain-containing pr |
| 597.13 | 303 | 49 | 351 | A0A2R8Y6K8\|A0A2R8Y6K8_HUMAN Mucin-19 (Fragment) OS=Homo sapiens OX |
| 596.79 | 69 | 940 | 1008 | A0A590UJ96\|A0A590UJ96_HUMAN Uncharacterized protein OS=Homo sapien |
| 596.13 | 94 | 2322 | 2415 | A0A2R8Y7Q1\|A0A2R8Y7Q1_HUMAN Nucleosome-remodeling factor subunit B |
| 595.61 | 97 | 1519 | 1615 | Q9UHV7\|MED13_HUMAN Mediator of RNA polymerase II transcription sub |
| 594.85 | 30 | 2415 | 2444 | P46531\|NOTC1_HUMAN Neurogenic locus notch homolog protein 1 OS=Hom |
| 594.48 | 260 | 1 | 260 | H7BXV5\|H7BXV5_HUMAN Collagen alpha-1(XVIII) chain (Fragment) OS=Ho |
| 594.14 | 65 | 499 | 563 | P48668\|K2C6C_HUMAN Keratin, type II cytoskeletal 6C OS=Homo sapien |
| 593.71 | 56 | 122 | 177 | P08151\|GLI1_HUMAN Zinc finger protein GLI1 OS=Homo sapiens OX=9606 |
| 592.95 | 129 | 164 | 292 | A0A2R8YGM9\|A0A2R8YGM9_HUMAN Eyes absent homolog OS=Homo sapiens OX |
| 592.50 | 224 | 261 | 484 | A0A669KB39\|A0A669KB39_HUMAN Collagen alpha-1(XIII) chain (Fragment |
| 590.14 | 96 | 1291 | 1386 | P51610\|HCFC1_HUMAN Host cell factor 1 OS=Homo sapiens OX=9606 GN=H |
| 589.63 | 101 | 467 | 567 | Q9ULI3\|HEG1_HUMAN Protein HEG homolog 1 OS=Homo sapiens OX=9606 GN |
| 589.16 | 103 | 2461 | 2563 | O75962\|TRIO_HUMAN Triple functional domain protein OS=Homo sapiens |
| 589.08 | 40 | 2269 | 2308 | Q5T1R4\|ZEP3_HUMAN Transcription factor HIVEP3 OS=Homo sapiens OX=9 |
| 588.79 | 77 | 146 | 222 | Q9UIF8\|BAZ2B_HUMAN Bromodomain adjacent to zinc finger domain prot |
| 587.63 | 119 | 1350 | 1468 | H0YHC1\|H0YHC1_HUMAN Mediator of RNA polymerase II transcription su |
| 587.09 | 74 | 600 | 673 | Q8N2Y8\|RUSC2_HUMAN Iporin OS=Homo sapiens OX=9606 GN=RUSC2 PE=1 SV |
| 586.73 | 29 | 4680 | 4708 | Q6V0I7\|FAT4_HUMAN Protocadherin Fat 4 OS=Homo sapiens OX=9606 GN=F |
| 586.72 | 109 | 981 | 1089 | H7BY37\|H7BY37_HUMAN Histone-lysine N-methyltransferase 2C (Fragmen |
| 586.39 | 100 | 1407 | 1506 | Q8NEV8\|EXPH5_HUMAN Exophilin-5 OS=Homo sapiens OX=9606 GN=EXPH5 PE |
| 586.29 | 97 | 124 | 220 | Q9HCD6\|TANC2_HUMAN Protein TANC2 OS=Homo sapiens OX=9606 GN=TANC2 |
| 585.86 | 74 | 1790 | 1863 | Q5HYC2\|K2026_HUMAN Uncharacterized protein KIAA2026 OS=Homo sapien |
| 584.09 | 61 | 206 | 266 | Q7Z5J4\|RAI1_HUMAN Retinoic acid-induced protein 1 OS=Homo sapiens |
| 583.26 | 60 | 1154 | 1213 | A0A140T8Y6\|A0A140T8Y6_HUMAN NOTCH4 OS=Homo sapiens OX=9606 GN=NOTC |
| 581.94 | 29 | 4682 | 4710 | A0A6Q8JR05\|A0A6Q8JR05_HUMAN Protocadherin Fat 4 OS=Homo sapiens OX |

| | | | | |
|---|---|---|---|---|
| 581.32 | 99 | 480 | 578 | Q99700\|ATX2_HUMAN Ataxin-2 OS=Homo sapiens OX=9606 GN=ATXN2 PE=1 S |
| 580.27 | 62 | 385 | 446 | Q2KHR3\|QSER1_HUMAN Glutamine and serine-rich protein 1 OS=Homo sap |
| 579.45 | 72 | 247 | 318 | O60299\|LZTS3_HUMAN Leucine zipper putative tumor suppressor 3 OS=H |
| 577.86 | 81 | 1549 | 1629 | Q6N021\|TET2_HUMAN Methylcytosine dioxygenase TET2 OS=Homo sapiens |
| 577.86 | 81 | 1570 | 1650 | E7EQS8\|E7EQS8_HUMAN Methylcytosine dioxygenase TET OS=Homo sapiens |
| 577.43 | 71 | 64 | 134 | Q96T58\|MINT_HUMAN Msx2-interacting protein OS=Homo sapiens OX=9606 |
| 577.02 | 184 | 347 | 530 | Q86VE3\|SATL1_HUMAN Spermidine/spermine N(1)-acetyltransferase-like |
| 577.02 | 184 | 347 | 530 | A0A2R8YFQ0\|A0A2R8YFQ0_HUMAN Spermidine/spermine N(1)-acetyltransfe |
| 575.93 | 119 | 935 | 1053 | A0A3B3IS46\|A0A3B3IS46_HUMAN Mediator of RNA polymerase II transcri |
| 575.60 | 307 | 51 | 357 | O95429\|BAG4_HUMAN BAG family molecular chaperone regulator 4 OS=Ho |
| 575.27 | 64 | 657 | 720 | P15822\|ZEP1_HUMAN Zinc finger protein 40 OS=Homo sapiens OX=9606 G |
| 575.13 | 77 | 833 | 909 | A0A590UJW6\|A0A590UJW6_HUMAN Zinc finger CCHC domain-containing pro |
| 573.38 | 132 | 173 | 304 | Q13151\|ROA0_HUMAN Heterogeneous nuclear ribonucleoprotein A0 OS=Ho |
| 573.38 | 143 | 1 | 143 | O15534\|PER1_HUMAN Period circadian protein homolog 1 OS=Homo sapie |

### Supporting Figures

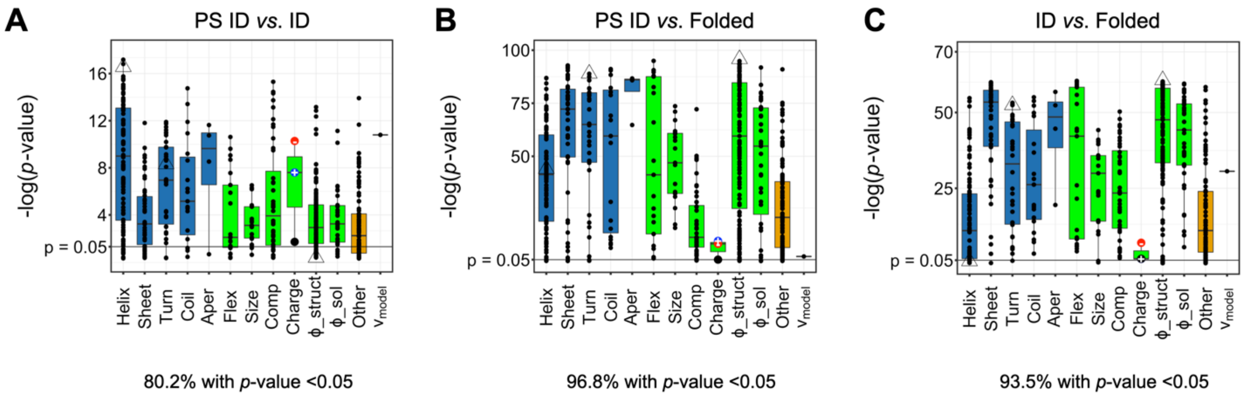

**Figure S1. Comparing means in the sequence sets using a nonparametric test.** *A-C*, *p*-values calculated by the Mann-Whitney *U*-test, shown as -log(*p*-value), compares set means in 567 amino acid scales and $v_{model}$. Here, the use of colors and symbols are identical to that used in Figure 2, where conformation-based scales are grouped by type and highlighted by blue boxplots, and physicochemical-based scales are grouped by type and highlighted by green boxplots. Scales (e.g., refractivity, crystal melting point) that did not easily map into a conformation-based or physicochemical-based group were combined separately (Other; orange boxplot). Boxplots show the dataset median (50$^{th}$ percentile) with the central bar, and the vertical width spans the 25$^{th}$ to 75$^{th}$ percentiles. Open triangles highlight the smallest *p*-value from Welch's *t*-test when comparing means in the PS ID and ID sets, which was from an α-helix propensity scale, the smallest *p*-value from Welch's *t*-test when comparing means in either ID set with the folded set, which was from a structure-based hydrophobicity scale, and the β-turn propensity scale used in ParSe (also provided for reference).

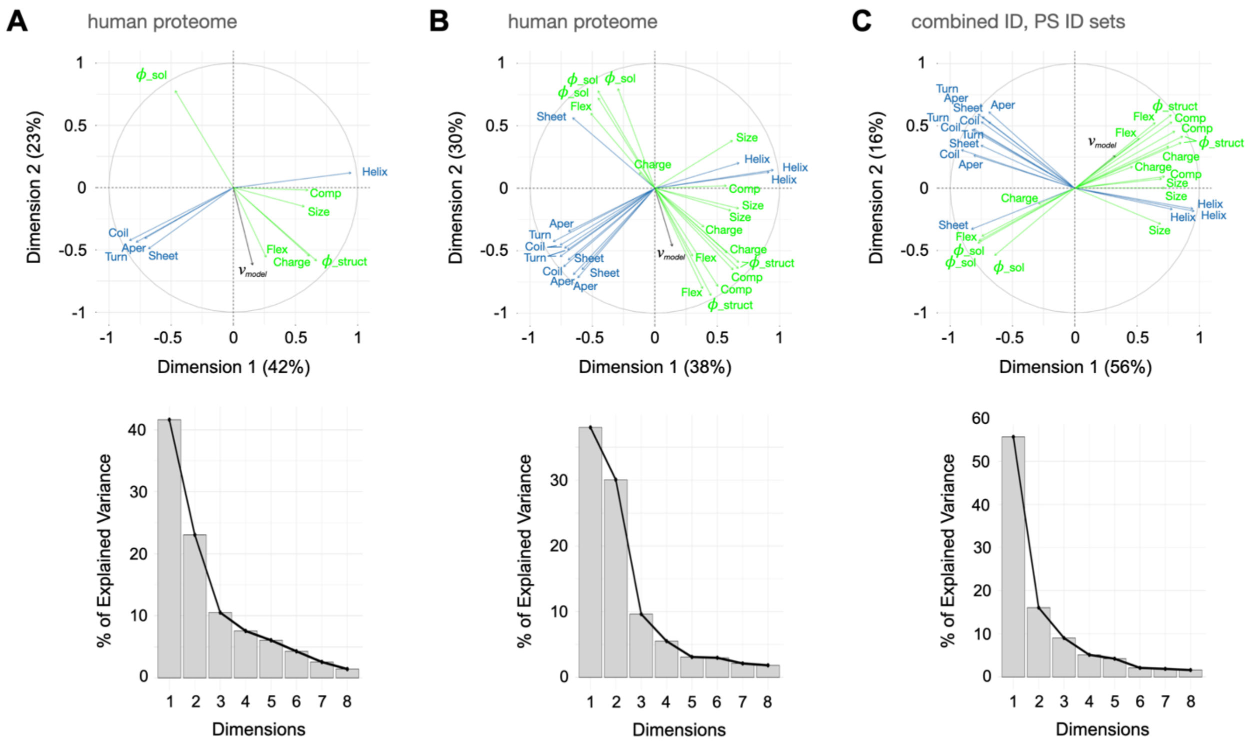

**Figure S2. Modes of variance in the sequence sets arising from different amino acid property scales.** *A-C*, bidimensional plots (top figures) from PCA showing the modes of variance in the human proteome and the combined ID sets (PS ID and ID) arising from conformation- (blue arrows) and physicochemical-based (green arrows) scales relative to the two principal components of variance, given as Dimension 1 and Dimension 2. Scree plots (bottom figures) showing the percent of the total variance in the human proteome and in the combined set of ID sequences that is captured by each principal component (i.e., dimension).

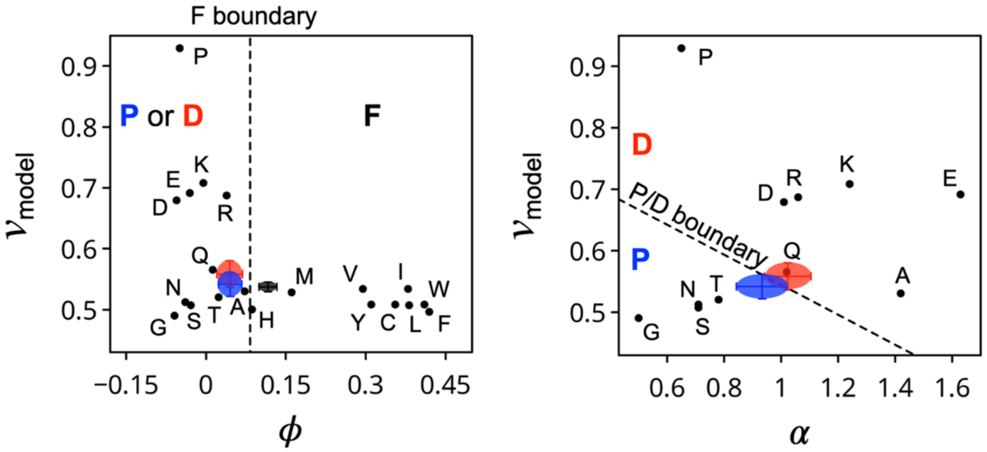

**Figure S3. Comparing hydrophobicity, α-helix propensity, and $v_{model}$ in homopolymers.** Hydrophobicity ($\phi$) and α-helix propensity (α) were calculated using the scales from Vendruscolo and coworkers (14) and Tanaka and Scheraga (15), respectively, in homopolymers ($N$ = 100) where amino acid type is identified by its one-letter code. Filled circles show the mean and standard deviation in $\phi$, α, and $v_{model}$ in the PS ID (blue), ID (red), and folded sets (black).

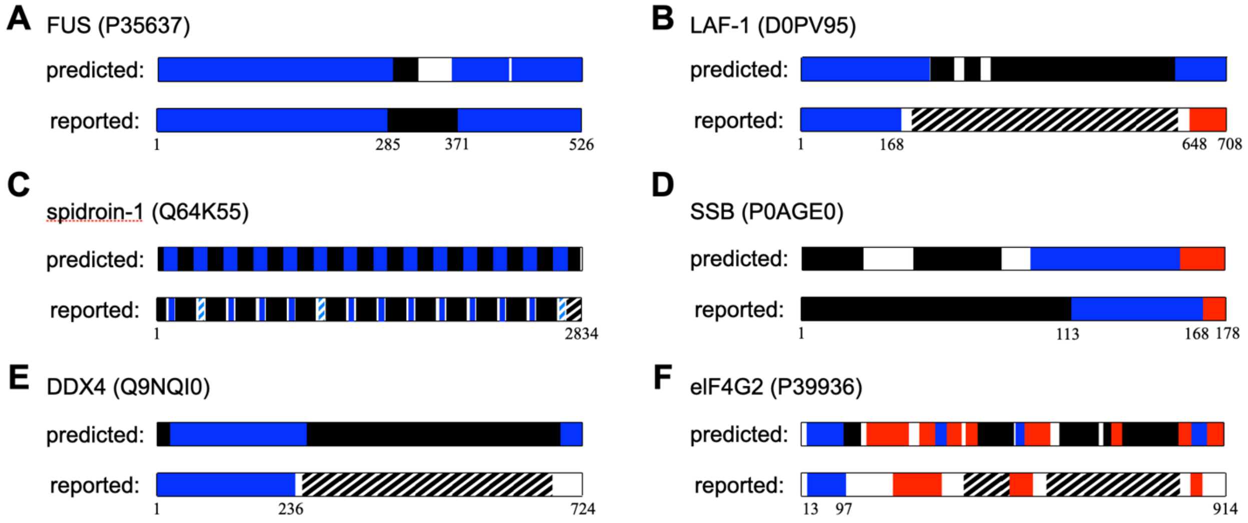

**Figure S4. Predicting protein regions that drive LLPS.** ParSe v2 was applied to the whole sequences of proteins with diverse reported mechanisms driving LLPS. The proteins are identified by name and UniProt accession number. Contiguous regions ($N \geq 20$) that were 90% of only one label, P, D, or F were colored blue, red, or black, respectively, to represent predicted PS, ID, or folded regions. Striped represents ≥50% identity to a known PS IDR (blue) or folded protein (black).

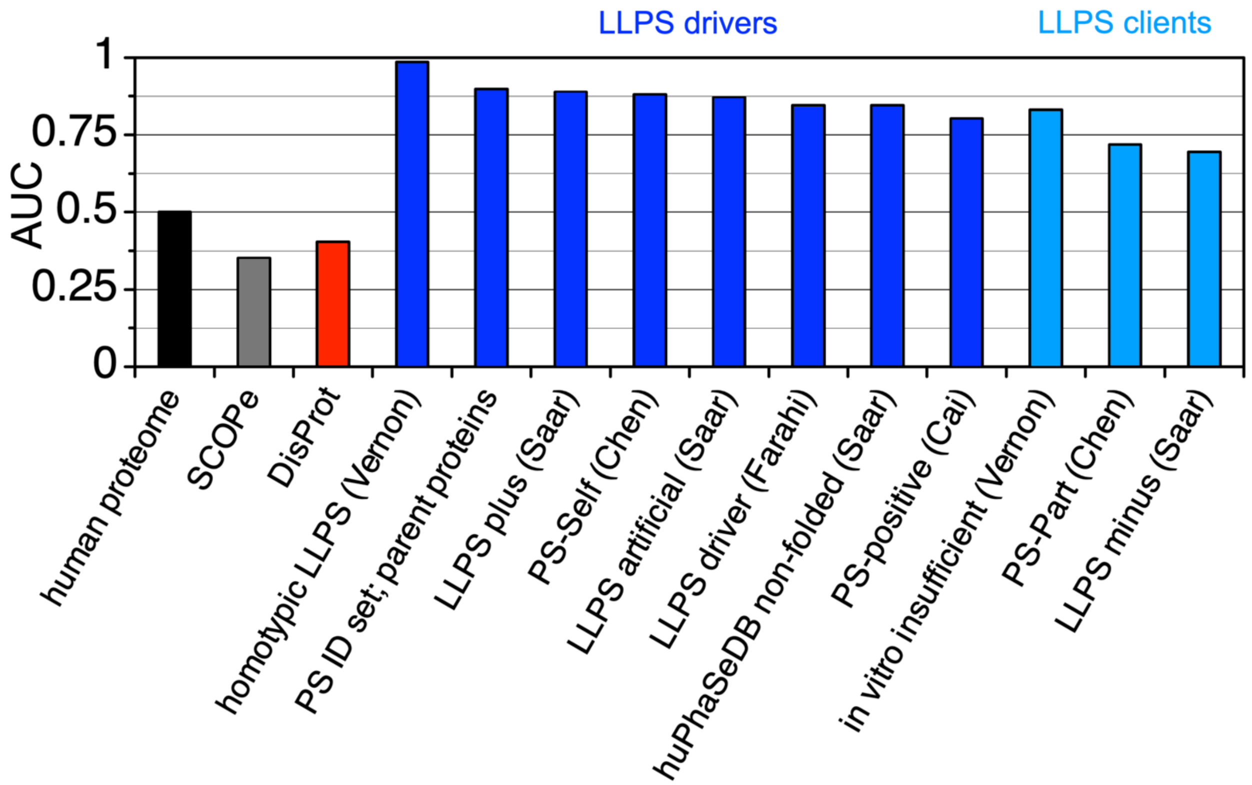

**Figure S5. LLPS driver sequences have AUC >0.8 when compared against the human proteome.** AUC calculations used the human proteome as the comparison set, and recall was based on the summed P classifier distance, as described in Figure 4. AUC values for SCOPe (grey) and DisProt (red) are reproduced from Figure 4C. LLPS driver sets (blue) are from Vernon et al (3), representing a set of proteins that have been verified *in vitro* to exhibit homotypic phase separation behavior (referred to as "*in vitro* sufficient" by Vernon), the parent proteins of the PS ID sequence set from the current study, and LLPS driver sets from Saar et al (16), Chen et al (17), Farahi et al (18), and Cai et al (19), where the sets are identified by the names used for these sets in each study. For comparison, light blue shows AUC for protein sets thought to have lower potential for phase separation (compared to the driver sets) because these proteins require partners (Vernon (3) and Chen sets (17)) and/or relatively high protein concentrations (>100 µM; Saar set (16)) for LLPS.

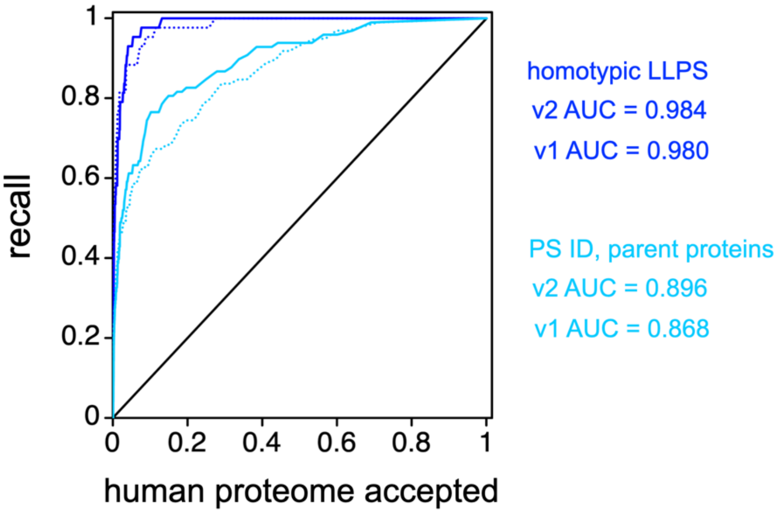

**Figure S6. ParSe v2 shows improved recall compared to the original version.** Homotypic LLPS is the Vernon et al set of proteins that have been verified *in vitro* to exhibit homotypic phase separation behavior (3). Solid lines are ParSe v2 results, while stippled lines are from the original ParSe algorithm. Data in this figure is a reproduction of the results in Figure 4A. Calculated AUC values are indicated to the right of the figure.

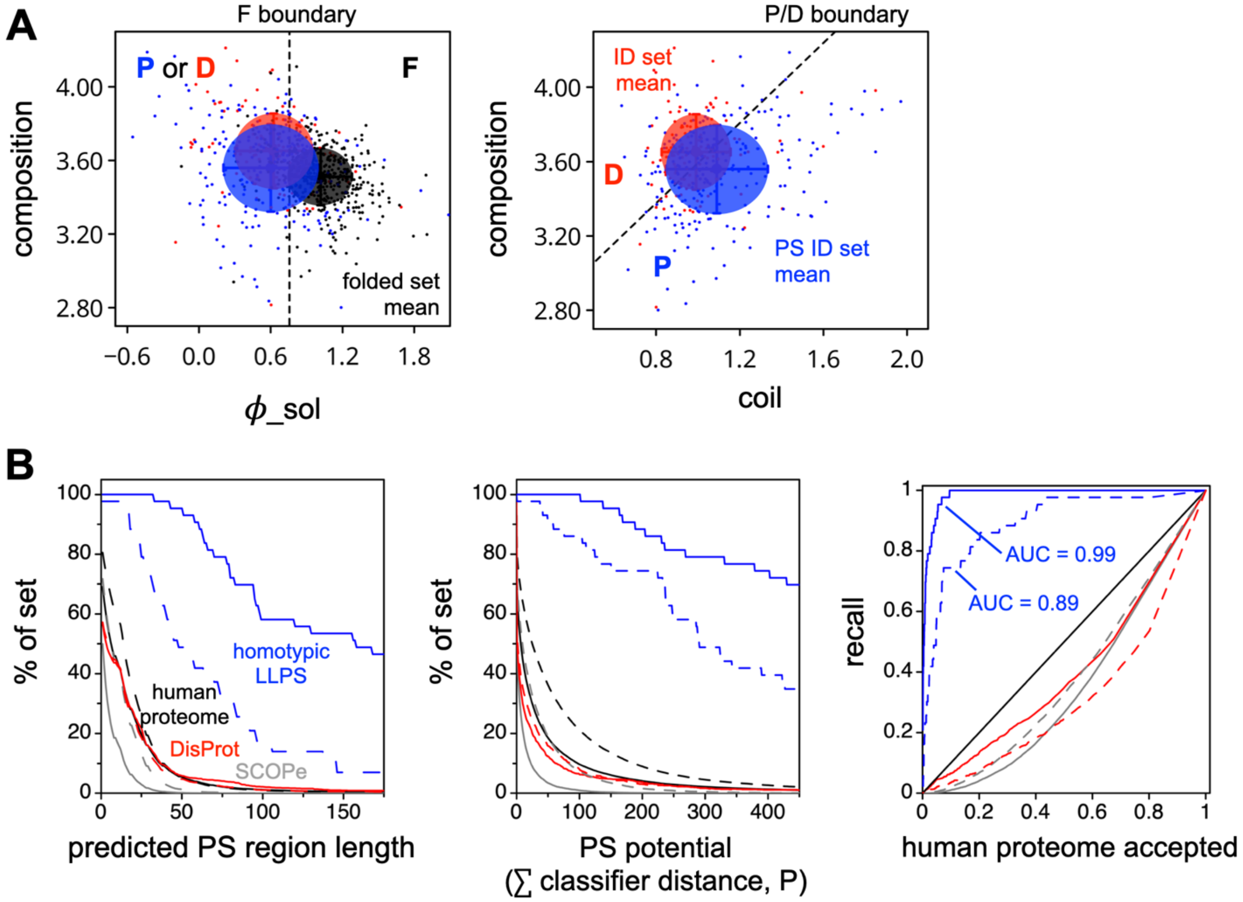

**Figure S7. ParSe v2 shows reduced recall when using scales with weaker predictive value.** *A*, the structure-based hydrophobicity scale from Vendruscolo and coworkers (14) was substituted for a solution-based hydrophobicity scale from Wilce et al (20) with *t*-test *p*-values of 3.4E-21 and 1.7E-18 when comparing means in the folded and PS ID and folded and ID sets, respectively. This solution-based hydrophobicity scale was used to identify F windows from P or D. A composition-based scale from Jukes et al (21), with a *t*-test *p*-value of 7.4E-08 when comparing means in the PS ID and ID sets, and a coil propensity scale from Isogai et al (22), with a *t*-test *p*-value of 4.6E-08 when comparing means in the PS ID and ID sets, were used to identify P windows from D. *B*, when using these weaker scales with ParSe v2 (dashed lines), the overall predictive value, as judged by AUC (left-most figure), decreased relative to ParSe v2 when using the top-performing scales (solid lines).

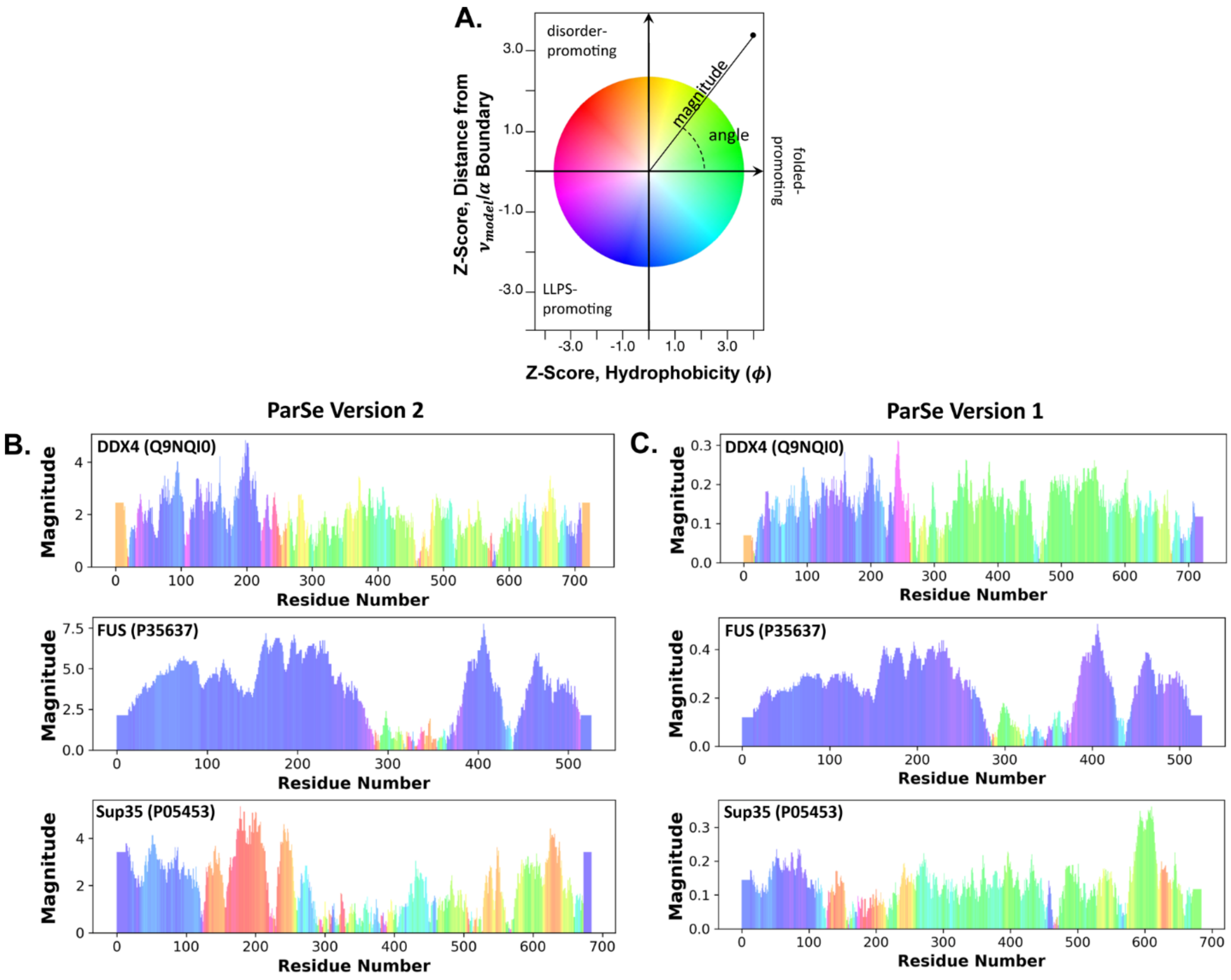

**Figure S8. ParSe v2 sequence predictions exhibit the same LLPS patterns as ParSe v1 predictions.** *A*, a modified color wheel scheme where each amino acid window is assigned a normalized hydrophobicity (x-axis) and normalized distance relative to the boundary line between α-helix propensity and $v_{model}$ ($v_{model}$ = -0.244•α-helix propensity + 0.789, see text). Positive y-axis values correspond to D-labeled windows (disorder-promoting, to the right of the P/D boundary in Figure 3B), and negative values correspond to P-labeled windows (LLPS-promoting, to the left of the P/D boundary in Figure 3B). A Z-score is used to normalize distances relative to the statistical distribution of the training sets (P, D, and F). As before (main text, Figure 1B), green regions correspond to F windows, blue/purple regions correspond to P windows, and red regions correspond to D windows. The magnitude represents the distance from the average hydrophobicity and $v_{model}/\alpha$ metrics. *B*,*C* color wheel predictions for Ddx4, FUS, and Sup35 based on Parse v2 (*B*) and ParSe v1 (*C*). UniProt IDs are given in parentheses. The color and magnitude for each residue window are mapped as described in (*A*). As before, each sequence partitions into regions that are mostly folded (green), disordered (red), and LLPS-promoting (blue/purple).

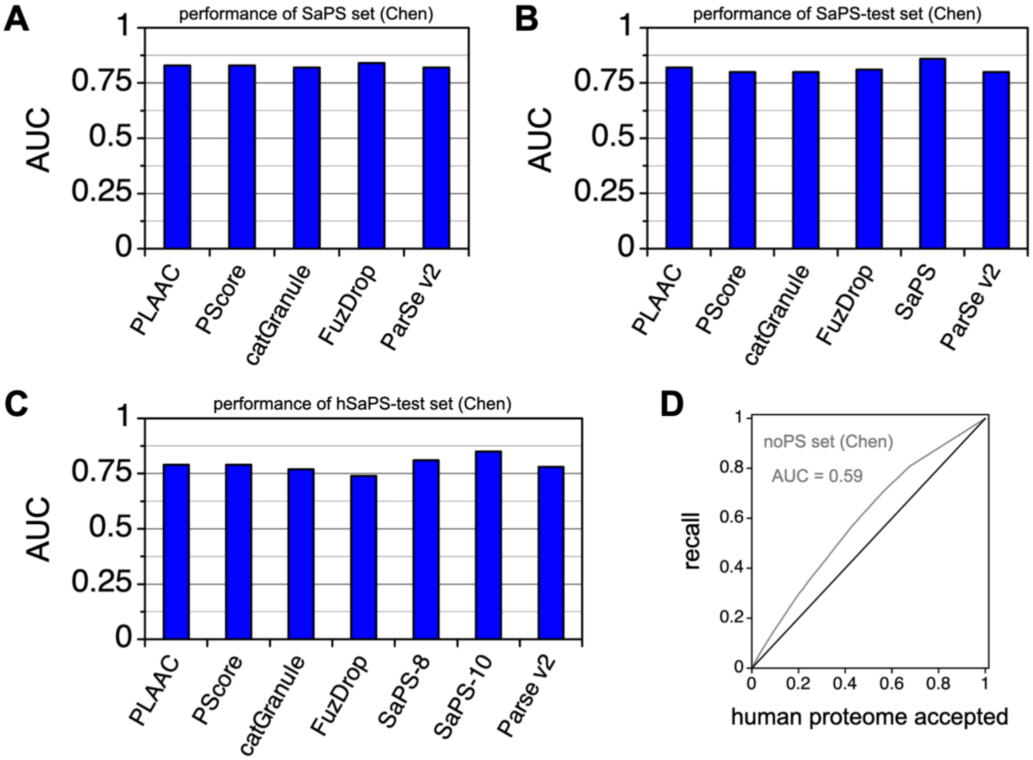

**Figure S9. ParSe v2 shows similar predictive accuracy as other LLPS predictors.** AUC values for PLAAC, PScore, catGranule, FuzDrop, SaPS, SaPS-8, SaPS-10 are reproduced from the scores given in Figures 1D, 2E, and S2B in Chen et al (17). AUC values for ParSe v2 used recall based on the summed P classifier distance, as described in Figure 4. *A*, SaPS, *B*, SaPS-test, and *C*, hSaPS-test sets were evaluated against the NoPS set. These sequence sets were obtained from Chen et al (17). *D*, recall in the NoPS set compared to the human proteome gives AUC >0.5, indicating that ParSe v2 predicts the NoPS set is enriched in PS regions.

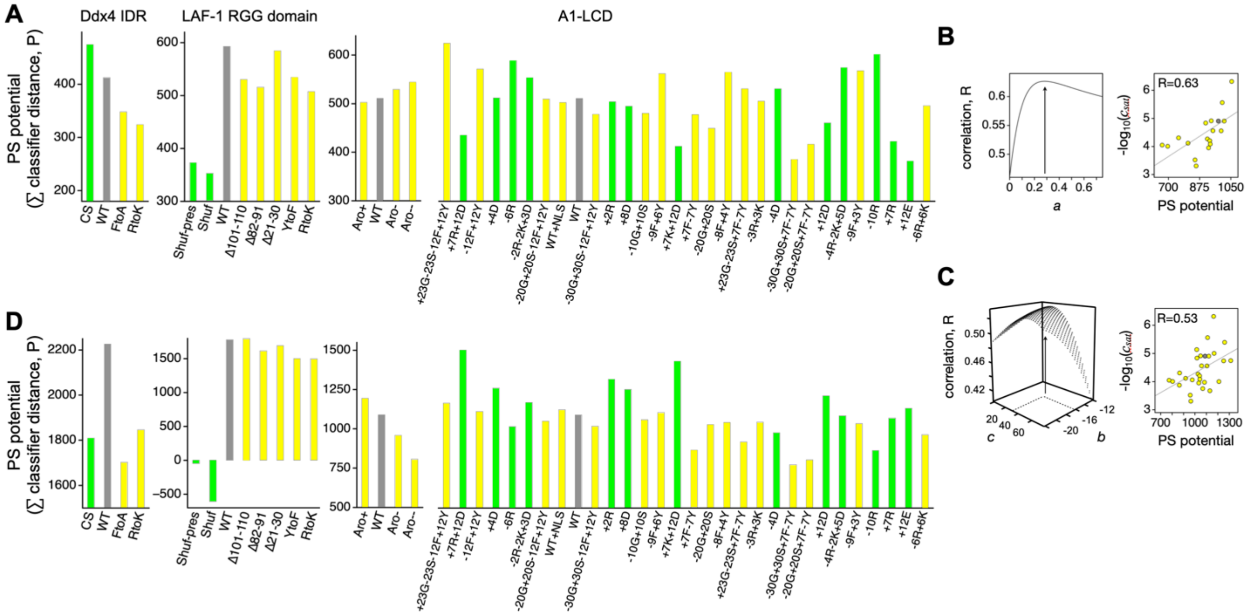

**Figure S10. Predicting mutation effects on phase separation behavior by training against $c_{sat}$.** *A*, the summed classifier distance of P-labeled positions was used to calculate a phase-separating (PS) potential from sequence. Mutants were grouped by experimental study and colored grey for wildtype (WT), yellow for mutants with both *NCPR* and *SCD* identical to the WT values, and green otherwise (non-WT *NCPR* and *SCD*). Placement left-to-right within a study follows the reported PS potential in rank, from high-to-low, for comparison to the predicted PS potential. A1-LCD mutants used $c_{sat}$ to establish rank. *B*, A1-LCD mutants with *NCPR* and *SCD* matching the WT values were used to fix *a* in Equation 3 by optimizing the correlation of Parse-calculated PS potential (including $U_\pi$) to -log$_{10}$($c_{sat}$); the right figure shows the optimal correlation. *C*, similarly, all A1-LCD mutants with experimental $c_{sat}$ were then used to fix *b* and *c* in Equation 4 by optimizing the correlation of ParSe-calculated PS potential (including $U_\pi$ and $U_q$) to -log$_{10}$($c_{sat}$); the right figure shows the optimal correlation. *D*, ParSe-calculated PS potentials (including $U_\pi$ and $U_q$ optimized to -log$_{10}$($c_{sat}$)) for the mutant and WT sequences.

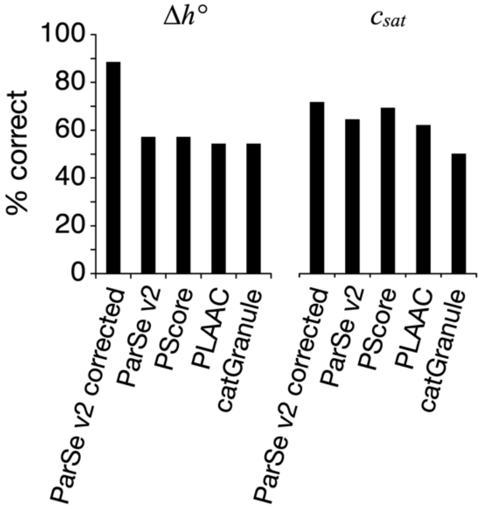

**Figure S11. ParSe v2 and other LLPS predictors show similar accuracy for predicting mutation effects.** Each predictor was used to rank the mutant sequences in order of phase separation potential, for the set of mutants shown in Figures 5 (rank determined by $\Delta h°$) and S10 (rank determined by $c_{sat}$). Percent correct is the number of mutant sequences that correctly predicted an increase or decrease relative to the wildtype sequence (by the specified predictor), divided by the total number of mutants and given as a percentage. Granule propensity was used for the catGranule score and LLR was used for the PLAAC score. "ParSe v2 corrected" refers to PS potential (sum of P-labeled windows) including $U_\pi$ and $U_q$.

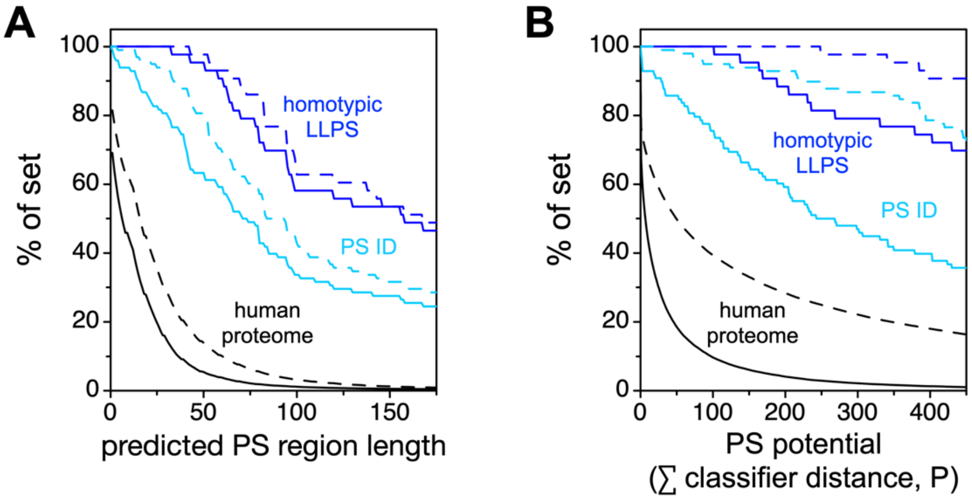

**Figure S12. $U_\pi$ and $U_q$ effects on ParSe predicted PS regions and potential.** *A*, ParSe v2 (solid lines) and ParSe v2 including $U_\pi$ and $U_q$ in the calculations (dashed lines) were used to identify regions in proteins that were ≥90% labeled P, which are referred to as phase-separating, PS, regions. Shown by the y-axis is the percent of proteins in a set with PS regions at least as long as the length indicated by the x-axis. The human proteome (UniProt reference proteome UP000005640) is given by black lines; a set of *in vitro* sufficient homotypic LLPS proteins by blue lines; and the full sequences of the proteins in the PS ID set by light blue lines. *B*, the summed P classifier distance was calculated for the protein sets in panel A, using both ParSe v2 (solid lines) and ParSe v2 including $U_\pi$ and $U_q$ (dashed lines). Shown by the y-axis is the percent of proteins in a set with a summed P classifier distance at least as much as the value indicated by the x-axis. Lines were colored using the same coloring scheme as in panel A.
